# Mapping chromothripsis across human cancer cell lines reveals genome evolution and therapeutic vulnerabilities

**DOI:** 10.64898/2026.10.30.763015

**Authors:** Gil Leor, Tal Ben-Yishay, Adi Tarrab, Yonatan Eliezer, Tom Winkler, Michal Goetz, Linoy Raz, Eli Reuveni, Ron Shamir, Uri Ben-David

## Abstract

Chromothripsis is a catastrophic chromosomal rearrangement found in ∼40% of primary tumors, yet its landscape in cancer cell lines (CCLs) remains uncharacterized. To address this gap, we developed a pipeline to detect chromothripsis-like events from copy-number data alone, and mapped their occurrence across 1,646 human CCLs. Chromothripsis patterns were broadly conserved between primary tumors and CCLs, but CCLs displayed some distinct features and a significantly higher prevalence of chromothripsis. Chromothripsis continued to emerge during culture propagation, likely contributing to its increased prevalence in CCLs. Integrating genomic and functional data revealed selective vulnerabilities: chromothriptic cells were resistant to DNA damage-inducing agents but highly sensitive to histone deacetylase inhibitors (HDACi). Using isogenic models, we validated that HDACi could synergistically sensitize chromothriptic cells to DNA damage-inducing agents. Our study establishes the first comprehensive resource of chromothripsis in CCLs and reveals functional consequences and therapeutic vulnerabilities of chromothripsis-positive cancers.

## Introduction

Chromosomal instability (CIN) is a well-established hallmark of cancer, encompassing a broad spectrum of chromosomal aberrations that disrupt cellular homeostasis and genomic integrity^1^. Chromothripsis (CT) is a major manifestation of CIN, characterized by the shattering and massive chromosomal rearrangements of one or more chromosomes, which result in oscillating copy number (CN) states^2–6^. This catastrophic event can profoundly affect gene and protein expression patterns, constituting an evolutionary “burst” that can influence the tumorigenic process by simultaneously affecting hundreds of genes^2–6^. Primary mechanisms that induce CT include rupture of micronuclei envelopes^7–10^, chromosome bridges^7,11,12^, and severe telomere attrition^7,12–14^. These events are associated with extensive double strand breaks (DSB), which can overwhelm the DNA damage repair machineries^15,16^.

Recent studies have extensively characterized CT in human tumors^17,18^, revealing that CT affects nearly half of all cancer cases and can form tumor-promoting genomic alterations, including clinically relevant druggable fusions^18^ and extra chromosomal DNA (ecDNA)^19^. This underscores the crucial role of CT in cancer pathogenesis and its therapeutic potential^17,18,20^. From a clinical perspective, CT tends to negatively impact patient survival across various cancer types, often correlating with poor prognostic outcomes^17,18^. Nonetheless, whether CT is associated with the response to specific anticancer therapeutics, and whether it could serve as a biomarker or a molecular target in cancer therapy, remains to be determined^20^.

Human cancer cell lines (CCLs), derived from various types of human tumors and propagated *in vitro*, are a crucial resource in cancer biology. The use of CCLs enables researchers to investigate cellular mechanisms, genetic mutations, and the effects of potential therapeutic agents under controlled conditions^21–31^. However, the use of CCLs is not without significant limitations, as cell lines are grown in isolation whereas tumors are a complicated ecosystem of cell types; the *in vitro* selection pressures differ from the *in vivo* ones, resulting in genomic evolution^32^; and the two-dimensional culture of CCLs does not mimic their growth within the complex tumor microenvironment. Although these constraints limit the use of CCLs in studying certain biological phenomena^33^, human CCLs remain a fundamental staple of cancer research, providing crucial insights into cancer cell-intrinsic traits and cancer cell vulnerabilities that can be translated to the clinic^22,29–31,34^. Importantly, while CT has been extensively studied in patient tumors^17,18^, the landscape of CT in CCLs remains poorly characterized.

Korbel et al. (2013) proposed several criteria for CT inference, including the presence of tens to hundreds of clustered rearrangements, oscillating CN states, and an enrichment of DNA breakpoints in specific chromosomal regions^3^. The measurement of these criteria heavily relies on whole-genome sequencing (WGS); precise information about breakpoint positions and genomic segment orientations is required. Consequently, several computational tools have been developed to detect CT^3,17,35^, most of which require whole-genome sequencing data. For example,

ShatterSeek, introduced by Cortés-Ciriano et al. (2020)^17^, relies on structural data for detecting CT events. Human CCLs, however, are often not characterized by WGS, limiting the ability to detect CT in CCLs using this approach. Fortunately, it has been suggested that CN data alone could be sufficient to detect CT-like patterns^36,37^, as alternating CN states, accompanied by alternating loss of heterozygosity (LOH), are a hallmark of this phenomenon^2,3^. CTLPScanner is a CT detection tool that only requires CN data^37^; based on likelihood ratio tests on CN alterations across the genome, it detects CT-like patterns. Importantly, since the criteria for CT are still evolving, no existing method is sufficiently accurate. To date, all major studies complemented automatic CT detection with careful manual inspection of suspected chromosomes using CN and/or SV _plots_17,35-37.

Here, we developed a new pipeline for detecting CT-like patterns (referred to as ‘CT events’ hereinafter) using CN data alone, improving upon current methods in parameters such as sensitivity, the exact determination of event borders, and the documentation of relevant structural and positional features (e.g., centromere or telomere involvement). We then mapped and characterized CT in a cohort of 1,646 human CCLs^21,22^, allowing the integration of this important genomic information into CCL data analyses and facilitating the selection of the most appropriate models for the study of CT *in vitro*. We also compared the incidence of CT in tumors^17,18^ to that of CCLs across lineages and individual chromosomes, highlighting similarities and differences between the human tumors and their cell line models. Comparing the CT landscapes between two large datasets of partly-overlapping CCLs^21,38^, we provide evidence for their genomic evolution throughout culture propagation. Together, these analyses shed light on the relevance, capabilities, and limitations of CCLs for modeling and studying CT. Next, using published omics data for our CCL cohort, we explored associations between CT and various genomic features, including other CIN manifestations and gene expression (GE)^21^. Finally, we harnessed large-scale genetic and chemical perturbation screens (e.g., CRISPR, RNAi, and drugs) that were previously performed in the CCLs^22–31^, in order to identify cellular vulnerabilities associated with CT, and experimentally validated clinically-relevant vulnerabilities in an isogenic system of CT-negative/CT-positive cells^8,39^. Specifically, we report that HDAC inhibition can selectively sensitize CT-positive cells to DNA damage-inducing agents, highlighting a combination drug therapy strategy of potential clinical relevance.

## Results

### A computational pipeline for chromothripsis detection

To map the landscape of CT in human CCLs, we utilized CN data derived from whole-exome sequencing (WES) or single-nucleotide polymorphism (SNP) arrays of 1,733 unique cell lines from the Cancer Cell Line Encyclopedia (CCLE) database^21,22^. We applied CTLPScanner^37^, a CN-based pipeline for CT-like pattern detection, to identify CT-like patterns across these human CCLs. As CTLPScanner does not accurately identify the boundaries of detected events, we modified the pipeline as follows (see **Methods** and **Supplementary Methods**): First, we ran CTLPScanner^37^ using lenient parameters that minimize the false negative calls at the expense of increasing the false positive (FP) rate. Subsequently, we used a log-likelihood ratio score to assess the confidence level of each interval within each CT-positive (CT+) event, and used this score to classify each detected event as positive (‘high-confidence’), borderline (“low-confidence”), or negative (CT- or ‘non-CT’). We then extended the genomic coordinates of the original CT-like event, as provided by CTLPScanner^37^, based on the prevalence and density of CN alterations (CNAs) in the proximity of the boundaries. Lastly, as done in previous works^17,18^, we visually inspected all positive and borderline CT-like events, and re-classified or border-adjusted them as necessary (**Fig. 1a**).

**Fig. 1.**
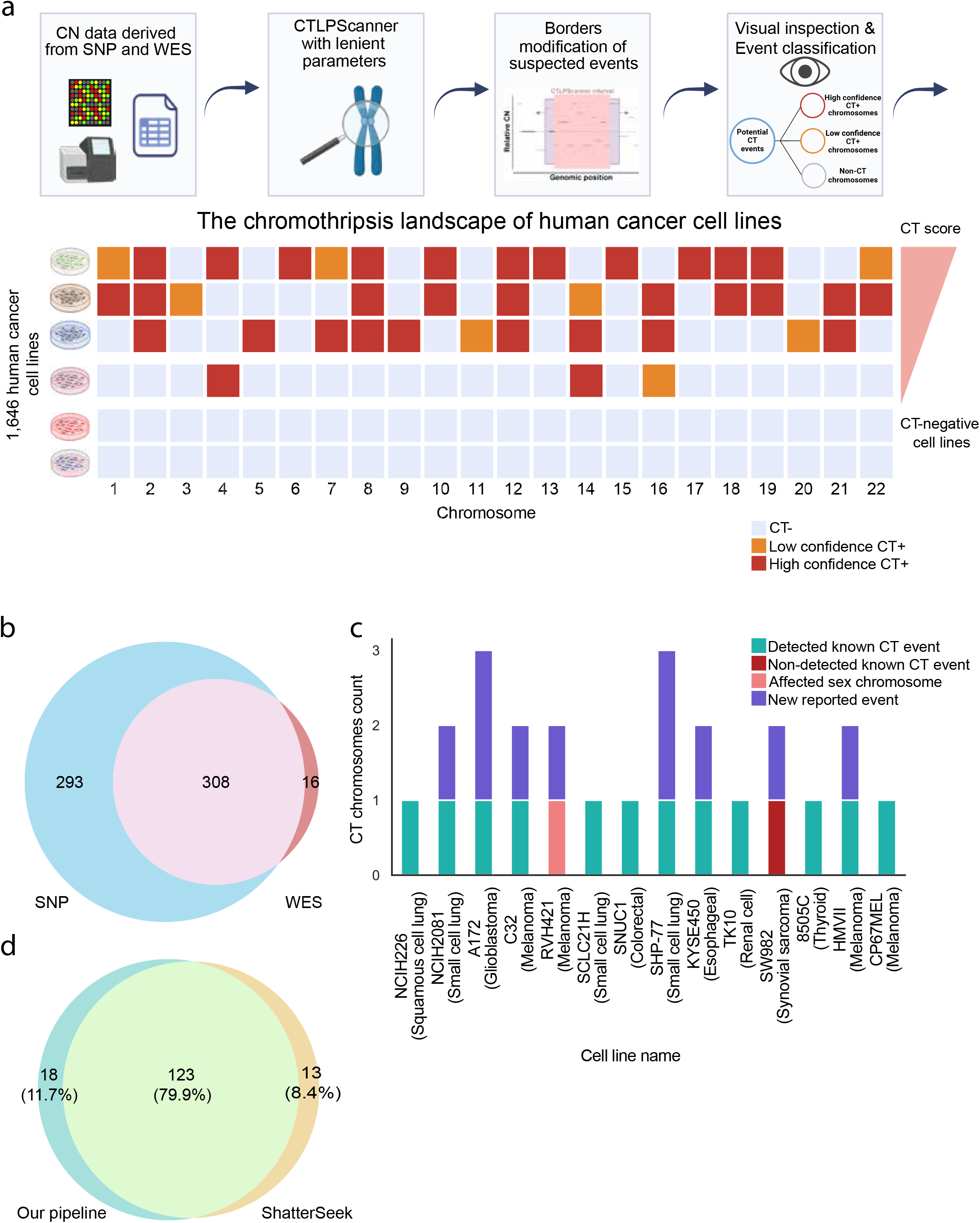
CN-based detection of chromothripsis-like patterns in human cancer cell lines. **(a)** Workflow for identifying chromothripsis-like events in human CCLs. CN data derived from SNP arrays and WES were analyzed using CTLPScanner^37^ with lenient parameters to detect putative CT events. Genomic intervals flagged as candidate events underwent further computational refinement to improve event boundary determination. Refined events were then classified into high-confidence CT+, low-confidence CT+, or CT negative (CT-) chromosomes. All high-confidence events were manually inspected for final calling of the presence and boundaries of CT-like events. This resulted in a comprehensive mapping of chromothripsis-like patterns across 1,646 cancer and 87 non-cancer human CCLs. **(b)** Venn diagram showing the overlap in CT detection based on SNP data vs. WES data. For this diagram, low-confidence events were considered positive. **(c)** Detection of CT events in 14 previously characterized human CCLs^2^. Known CT events that were successfully detected by our pipeline are marked in light blue, whereas the only previously detected event that was missed by our pipeline is marked in dark red. Events involving sex chromosomes (not included in our analysis) are indicated in pink, and events identified only by our pipeline are highlighted in purple. **(d)** Venn diagram showing the overlap in CT+ CCLs between our pipeline and the ShatterSeek^17^ algorithm across 325 human CCLs with available WGS data. Only high-confidence CT+ CCLs were considered for this analysis.

We also used visual inspection as a first assessment of the pipeline’s performance. Considering high-confidence events, precision was 0.91, recall was 0.96, and F1 score was 0.93 for the SNP data (**Extended Data Fig. 1a**); and precision was 0.77, recall was 0.93, and F1 score was 0.84 for the WES data (**Extended Data Fig. 1b**). To estimate the false negative (FN) rates, we manually inspected 400 random chromosomes classified as CT-by CTLPScanner^37^ (200 chromosomes per platform), and found a 0.5% FN rate for both SNP and WES data. When restricted to the subset of chromosomes flagged positive by CTLPScanner^37^ but reclassified as negative by our pipeline, the FN rate was similarly ∼0.5% for SNP data, but increased to ∼3% for WES data. Therefore, our automatic pipeline allowed us to capture the vast majority of CT-like events, and our manual inspection enabled us to exclude false positive calls, resulting in accurate mapping of CT-like CNA patterns across human CCLs. For downstream analyses, we mostly focused on CT-like events classified as ‘high-confidence’ events (**Methods** and **Supplementary Methods**).

As our analysis was based on both SNP and WES data, and our parameters were optimized for each platform separately, we next set out to compare the two platforms using 899 cell lines for which both data types were available. As expected, SNP-based CT calling was more sensitive than WES-based CT calling: with borderline events considered as positive, SNP data analysis detected ∼87% of the events identified by WES data, whereas WES data analysis detected only ∼18% of the events identified by SNP data; at the cell line level, SNP data detected ∼95% of the CT+ CCLs identified by WES data, whereas WES detected only ∼51% of the CT+ CCLs identified by SNP data, demonstrating the superior sensitivity of SNP-based inference (**Fig. 1b**). We conclude that both platforms can be used to accurately detect CT-like patterns (**Extended Data Fig. 1c**), with WES being less sensitive than SNP (as could be expected), explaining the higher FN rates for WES-based CT calling. Therefore, for downstream analyses, whenever both SNP and WES data were available for the same cell line, we used the SNP-based CT calling results.

### Comparison between our pipeline and existing chromothripsis-detection tools

To first compare our extended, optimized pipeline with the standard performance of CTLPScanner, we applied the tool’s recommended parameters^37^ (≥ 20 CNAs, LLR ≥ 8, and CN difference ≥ 0.3), visually inspected the called CT-like events, and compared the performance of the two pipelines. Considering only ‘high-confidence’ events, CTLPScanner^37^ showed similar precision to our pipeline (0.79 vs. 0.8) but lower recall (0.38 vs. 0.55) in the SNP data; and higher precision (0.92 vs. 0.78) but lower recall (0.27 vs. 0.35) in the WES data (**Extended Data Fig. 1d,e**). Our modified pipeline, therefore, increased the recall rate of CT-like events compared with the standard CTLPScanner pipeline^37^. Importantly, our pipeline also greatly improved the accuracy of boundary assignment for each detected event (**Extended Data Fig. 1f,g**).

As another validation of our pipeline, we asked whether our CT calling pipeline could accurately detect CT events that had been previously reported in the literature. Specifically, we examined 14 CCLs reported to be CT+ in the seminal study by Stephens et al. (2011)^2^. Our pipeline successfully detected 12 out of the 13 autosomal CT+ chromosomes (**Fig. 1c**, **Extended Data Fig. 2a-c**); sex chromosomes were excluded from our analysis). Moreover, for most CCLs, our analysis identified additional positive and borderline CT-like events, consistent with the original study’s focus on the most prominent events^2^.

A definitive detection of CT requires WGS data, which allows the identification and analysis of structural rearrangements, a defining feature of CT. However, WGS data are much less common than WES/SNP data, providing the rationale for our attempt to infer CT-like events based on CN data alone. To compare the performance of WGS-based CT calling to our CNA-based calling of CT-like events, we applied ShatterSeek^17^, a commonly-used tool for SV-based CT detection, to 325 CCLs for which both WES/SNP and WGS data were available^21^. ShatterSeek identified 288 high-confidence and 498 low-confidence CT+ chromosomes across 149 high-confidence and 101 low-confidence CT+ CCLs. Focusing on the high-confidence calls of CT+ CCLs, we observed 79.9% agreement between the pipelines (123 out of 154 CCLs), with 8.4% CT+ CCLs detected exclusively by ShatterSeek and 11.7% CT+ CCLs detected only by our pipeline (**Fig. 1d**). Comparing the specific CT events after excluding borderline calls (from at least one of the tools), ShatterSeek identified 246 high-confidence CT+ chromosomes, while our pipeline identified 437, with only 27.2% agreement in positive detections between the tools (**Extended Data Fig. 3a**). There are several probable reasons for this relatively low concordance, including the inherent inability of our pipeline to detect CN-neutral CT events^40^ (**Extended Data Fig. 3b**), as well as the inability of ShatterSeek to detect CT-like CN oscillations that are not accompanied by massive structural rearrangements (**Extended Data Fig. 3c**). Indeed, when visually inspecting events that were identified by our pipeline and not by ShatterSeek, we found that many of them showed clear CT-like CNA patterns (**Extended Data Fig. 3c**). Our pipeline, therefore, efficiently detects CT-like CNA patterns, and generally agrees with ShatterSeek^17^ whenever the CT event involves both SVs and CN oscillations.

### The chromothripsis landscape of human cancer cell lines

Our pipeline yielded a detailed map of CT-like events across human CCLs. For each event, we documented various features such as the event coordinates, whether events spanned centromeres or telomeres, the number of CN states and alterations, as well as additional characteristics (described below). These data are summarized in a detailed table of the CT-like events (**Supplementary Table 1**). In addition, we provide a cell line-level table that summarizes the CT status, metadata, and additional CIN features per cell line (**Supplementary Table 2**). A similar ShatterSeek-based CT events table is provided for the 325 CCLs for which WGS data were available (**Supplementary Table 3**). We also provide a detailed visual PDF report per cell line and chromosome, including ShatterSeek^17^ and Rawcopy^41^ plots when available (See **Extended Data Fig. 4a-d** for an example).

Out of the 1,646 human CCLs analyzed, high-confidence CT events were identified in 757 (46.0%) CCLs, and additional low-confidence (‘borderline’) CT events were identified in 202 (12.3%) CCLs (**Fig. 2a, Extended Data Fig. 5a**). The prevalence of CT-like events greatly varied across the 28 lineages (cancer types) that were analyzed (**Fig. 2a**). Some cancer types are vastly more chromothriptic than others (Pearson’s Chi-squared test, p-value < 2.2e-16), with esophagus and breast being the most chromothriptic (81.6% and 79.1%, respectively), and peripheral nervous system being the least chromothriptic (14.9%). Importantly, we also analyzed 41 cell lines of non-cancerous fibroblast origin. A CT-like event was detected with high-confidence only in one (2.4%) of the fibroblast cell lines, HS706T. Further examination revealed that this fibroblast cell line was very likely mislabelled in DepMap and actually originated from a bone cancer (as currently annotated in Cellosauros^42^ and CellModelPassposrt^38^). These findings demonstrate that CT is rare in non-cancer cells and further support the high specificity of our CT calling.

**Fig. 2.**
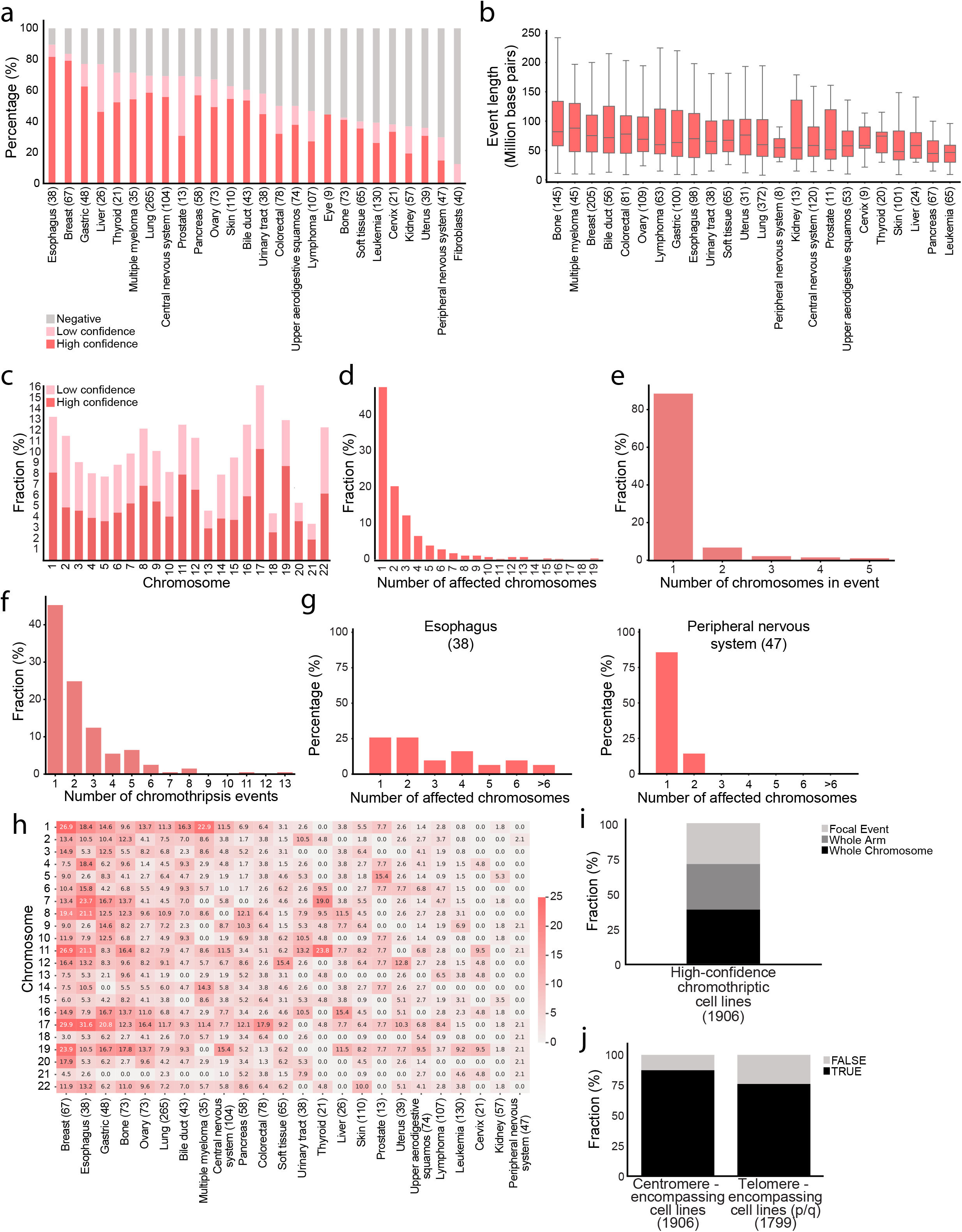
The landscape of chromothripsis across human cancer cell lines. **(a)** Chromothripsis prevalence across 1,639 human CCLs and 41 cancer-associated fibroblasts. Cell lineage (i.e., cancer type) affects chromothripsis prevalence (Pearson’s chi-squares test, p-value < 2.2e-16). The mean occurrence of high-confidence CT is 46.0%. 7 CCLs belonging to lineages with fewer than 5 samples were excluded from the plot. **(b)** CT length across lineages. The mean event length across all CCLs is 80 MBPs and the median is 62.9 MBPs. Lineage affects the mean length of CT events (One-way ANOVA, p-value < 0.0001). (**c)** CT prevalence across chromosomes. CT prevalence varies significantly across chromosomes (Chi-squared test, p-value < 2.2e-16). (**d),** Number of chromosomes affected by high-confidence CT across CCLs. 52.3% of the high-confidence CT+ CCLs involve two or more chromosomes. 32.0% of the high-confidence chromothriptic cells involve three or more chromosomes. 13.3% of the high-confidence chromothriptic cells involve five or more chromosomes. (**e)** Distribution of the number of chromosomes involved in individual CT events, defined using shared SVs between chromosomes within CT regions (≥2 SVs, 10 kb tolerance, transitivity applied). (**f)** Distribution of the number of distinct CT events per cell line, demonstrating a shorter “right tail” compared to the distribution of affected chromosomes (Fig. 2d), suggesting that cases with many affected chromosomes often stem from a smaller number of complex multi-chromosome CT events (means: 2.30 vs. 2.52, Mann-Whitney p-value = 0.24). **(g)** Number of chromosomes affected by high-confidence CT in esophagus and peripheral nervous system cancer types. Lineage affects the number of chromosomes involved in high-confidence CT (One-Way ANOVA, p-value = 8.3e-08). In esophagus cancer, the cancer type with the highest CT incidence (81.6%), 74.2% of the CT+ CCLs show at least two high-confidence chromothriptic chromosomes. In comparison, in peripheral nervous system cancer, with CT incidence of 14.9%, only 1 (14.3%) of the CT+ CCLs has >1 chromothriptic chromosomes. (**h)** CT prevalence across chromosomes and cancer types. The heatmap presents the prevalence of CT in each chromosome of each cancer type, the higher the prevalence the darker the color. CT prevalence across chromosomes is variable in most, but not all, cancer types (asterix reflects statistical significance of individual Pearson’s chi-squared tests for each lineage. ns = not significant). (**i)** The prevalence of whole-chromosome and chromosome-arm events out of all high-confidence CT+ events across cell lines. For determining whole p-arm and whole q-arm, only events that involve p and q arm, respectively, were assessed. For acrocentric chromosomes, whole q-arm events were considered as whole-arm. **(j)** Incidence of centromere (left) and telomere (right) involvement in high-confidence CT events in human CCLs. The majority of the high-confidence CT events involve the centromere and at least one telomere. 87.3% of all high-confidence events involve the centromere. 76% of the CCLs involve the (p or q) telomere. 107 acrocentric chromosomes (13, 14, 15, 21 and 22) with high-confidence CT were excluded from this analysis (due to the absence of copy number data for their p arms).

Next, we examined the genomic properties of the CT events themselves, focusing on their length, density (i.e., number of alterations within each event normalized to its length) and the number of copy number states (CNSs) that are involved (**Methods** and **Supplementary Methods**). When assessing the distribution of CT length, we observed that the mean CT-like event length across all CCLs varies between ∼8 million base pairs (MBPs) and ∼250 MBPs (mean: 80 MBPs; median: 62.9 MBPs), varying considerably across lineages (**Fig. 2b**; One-way ANOVA, p-value < 0.0001). Bone has the longest mean event length (96.6 MBPs) whereas leukemia has the shortest (54.8 MBPs). The mean percentage of the genome affected by CT-like events in high-confidence CT+ CCLs is 7.4%, and this number also varies across cancer types (**Extended Data Fig. 5b**; p-value = 2.01e-08), with the bone lineage presenting the highest percentage of the genome covered by CT (16.6%), and the peripheral nervous system presenting the lowest (3.1%). The mean number of alterations within a CT-like event is 35 (min: 7, max: 450). Although the length of CT-like events varied significantly across lineages (p-value < 0.001), the CT density did not vary as much (p-value = 0.04 & 0.20 for SNP & WES, respectively; **Extended Data Fig. 5c,d**). The mean number of CNSs (encompassing >3% of the event’s segments; **Methods** and **Supplementary Methods**) is 5.5, and the mean number of dominant CNSs (encompassing >15% of the event’s total length; **Methods** and **Supplementary Methods**) is 2.5. As with the other structural properties, the lineage significantly affects the mean number of CNSs (**Extended Data Fig. 6a,b**; p-value = 3.82e-07 and p-value = 2.81e-08, for all CNSs and dominant CNSs, respectively). When comparing CT prevalence between CCLs derived from primary tumors and those derived from metastases, only a slight increase is observed in metastasis-derived vs. primary-derived CCLs (**Extended Data Fig. 6c**, 54.2% vs. 48.7%, Chi-square, p-value = 0.233). Across cancer types, there was no significant difference in CT prevalence between metastasis-derived and primary-derived CCLs, with the notable exceptions of gastric and bone cancers, which showed significant (yet opposing) trends (**Extended Data Fig. 6d**). The similar prevalence of CT in metastasis- and primary tumor-derived CCLs mirrors its similar prevalence in metastases and primary tumors, as previously reported^2^.

### Chromosome-specific patterns of chromothripsis

Examination of CT prevalence throughout the genome revealed high variability across chromosomes (**Fig. 2c**; p-value < 2.2e-16), with chr17 being the most chromothriptic (10.3%), and chr21 being the least (1.4%). Similarly, CT prevalence varies greatly when considering chromosome-arms rather than entire chromosomes (**Extended Data Fig. 7a**; p-value < 2.2e-16); The most chromothriptic chromosome-arm is chr17q (10.1%), and the least chromothriptic chromosome-arm is chr20q (1.3%). We note that acrocentric chromosomes (13, 14, 15, 21 & 22) tend to be significantly less chromothriptic than non-acrocentric chromosomes (**Extended Data Fig. 7b**; 3.7% vs. 5.7% respectively, p-value = 2.53e-12). This observation is likely attributed to selection during cancer development, but may also result from chromosome-specific properties, including length, structure, and nuclear positioning, which predispose certain chromosomes to missegregation and micronucleation, key CT-initiating events^43–45^.

Importantly, 52.3% of the high-confidence CT+ CCLs exhibit CT in more than one chromosome, 32.0% of the high-confidence CT+ CCLs have three or more CT+ chromosomes, and, notably, 13.3% of the high-confidence CT+ CCLs have five or more CT+ chromosomes (**Fig. 2d**). In 16 CCLs, we observed CT-like patterns in >10 (!) chromosomes, and manual inspection of the CN landscapes of these CCLs suggests that these are all indeed CT-like events (**Extended Data Fig. 7c**).

Among the CT+ CCLs, 201 had WGS data with SV information, enabling the attribution of CT-affected chromosomes to individual CT events following Cortés-Ciriano et al.’s method (**Methods**)^17^. In this CCL subset, 552 affected chromosomes grouped into 460 events, revealing that most CT events (∼88%) affect only one chromosome, whereas ∼7% of the events involve two chromosomes, and ∼5% of the events involve three chromosomes or more (**Fig. 2e**). The distribution of affected chromosomes in this analysis resembles that of the full cohort (**Extended Data Fig. 7d**, **Fig. 2d**; means: 2.75 vs. 2.52, Mann-Whitney p-value = 0.47). Interestingly, when considering the number of CT events per cell line, the ‘right tail’ shortens (**Fig. 2f**; compare to **Fig. 2d** and **Extended Data Fig. 7d**), indicating that CCLs with multiple affected chromosomes often arise from fewer complex multi-chromosome CT events.

Like CT prevalence itself, the number of chromosomes affected by CT per cell line varies across cancer types (**Extended Data Fig. 7e**; one-way ANOVA, p-value = 8.36e-08): bone and breast cancers, for example, have 33.3% and 34.0% of their CCLs involving >6 chromothriptic chromosomes, respectively. As expected, the mean number of CT-affected chromosomes is highly correlated with the overall CT prevalence across cancer types (**Extended Data Fig. 8a**; Pearson’s R = 0.86, p-value = 1.1e-08), and with the overall fraction of the genome affected by CT (**Extended Data Fig. 8b**; Pearson’s R = 0.98; p-value = 5.44e-17). In esophageal cancer, the lineage with the highest CT incidence (81.6%), 74.2% of the CT+ CCLs show at least two high-confidence CT+ chromosomes. In comparison, in peripheral nervous system cancer, with CT incidence of 14.9%, only 1 (14.3%) of the CT+ CCLs has more than one high-confidence CT+ chromosome (**Fig. 2g**).

Cancer types differ not only in their tendency toward CT, and in the number of CT-affected chromosomes, but also in the involvement of specific chromosomes (**Fig. 2h**). This is more apparent in highly-chromothriptic cancer types. For example, chromosomes 1, 11, 17 and 19 are each involved in ∼25% of the CT-like events in breast cancer, whereas their relative involvement in CT-like events in gastric CCLs is much lower (**Fig. 2h**). Notably, some cancer types show a relatively high prevalence of CT-like events, without a strong inclination toward specific chromosomes. For example, over half of the CCLs are chromothriptic in cancers of the bile duct (57.5%), pancreas (64.7%) and urinary tract (51.5%), but there’s no enrichment for any specific chromosome (**Fig. 2h**; p-value > 0.05 for all chromosomes). Interestingly, although 57.5% of urinary tract CCLs are CT+, none of these events involves chromosome 17, the most chromothriptic chromosome overall (10.3% pan-cancer prevalence). Similar tissue-specific patterns were reported in other forms of chromosomal instability, such as aneuploidy and whole-genome doubling^46–48^.

We next assessed how many of the high-confidence CT events span the entire chromosome or chromosome-arm. We defined a whole-chromosome or chromosome-arm CT-like event as an event encompassing >80% of the chromosome or chromosome-arm, respectively. We found that 38.4% of the CT-like events encompass the entire chromosome, 32.4% of the CT-like events encompass the whole chromosome-arm, and 29.1% of the CT-like events are more focal (**Fig. 2i**). The tendency of CT to encompass the entire chromosome or chromosome-arm is both cancer-type-specific and chromosome-specific: some cancer types have a significantly higher portion of whole-chromosome events (**Extended Data Fig. 9a**; p-value = 4.1e-07). For example, bone cancer has a 60% whole-chromosome rate compared to only 20% in thyroid cancer. Similarly, some chromosomes tend to have a higher rate of whole-chromosome events, with chromosome 19 at the top (59.7% of the CT-like events) and chromosome 6 at the bottom (20.5%) (**Extended Data Fig. 9b**; p-value = 2.6e-34). The same is true when considering chromosome-arms rather than entire chromosomes (**Extended Data Fig. 9c**). We then analyzed the chromosomal position of high-confidence CT-like events. Across the human CCLs, 87.3% of events encompass the centromere, and 76% encompass the (p or q) telomere (**Fig. 2j**).

We note that the chromosome-bias in CT-like incidence, together with the tendency of CT in specific chromosomes to encompass the entire chromosome or chromosome-arm, likely contributes to the difference in CT event length across cancer types. For example, leukemia tends to involve smaller chromosomes in comparison to bone (p-value = 0.049; comparison of the relative prevalence of small chromosomes (12:22) vs. large chromosomes (1:11); **Fig. 2h**), in line with the smaller mean length of CT events in leukemia vs. bone cancer (54.8M vs. 96.6M, respectively; **Fig. 2b**). We also found a positive relationship between CT mean event length and the fraction of whole-chromosome events out of all CT-like events in the respective cancer type (**Extended Data Fig. 10a**; Pearson’s R = 0.45, p-value = 0.026). Interestingly, the larger the chromosome, the less likely it is to be entirely encompassed by a CT event (**Extended Data Fig. 10b**; Pearson’s R = -0.63, p-value = 0.0017).

### Chromothripsis association with chromosomal instability

We next sought to investigate the relationship between CT-like patterns and other CIN manifestations in human CCLs. First, we found that high-confidence CT+ CCLs tend to have higher aneuploidy scores (AS) than low-confidence and CT-CCLs (**Fig. 3a**; means: 18.2 vs. 14.9 & 12.6, respectively, One-way ANOVA, p-value < 2e-16). This is not surprising considering that both CT and aneuploidy are downstream consequences of chromosome mis-segregation and chromosome encapsulation in micronuclei^6,7^.

**Fig. 3.**
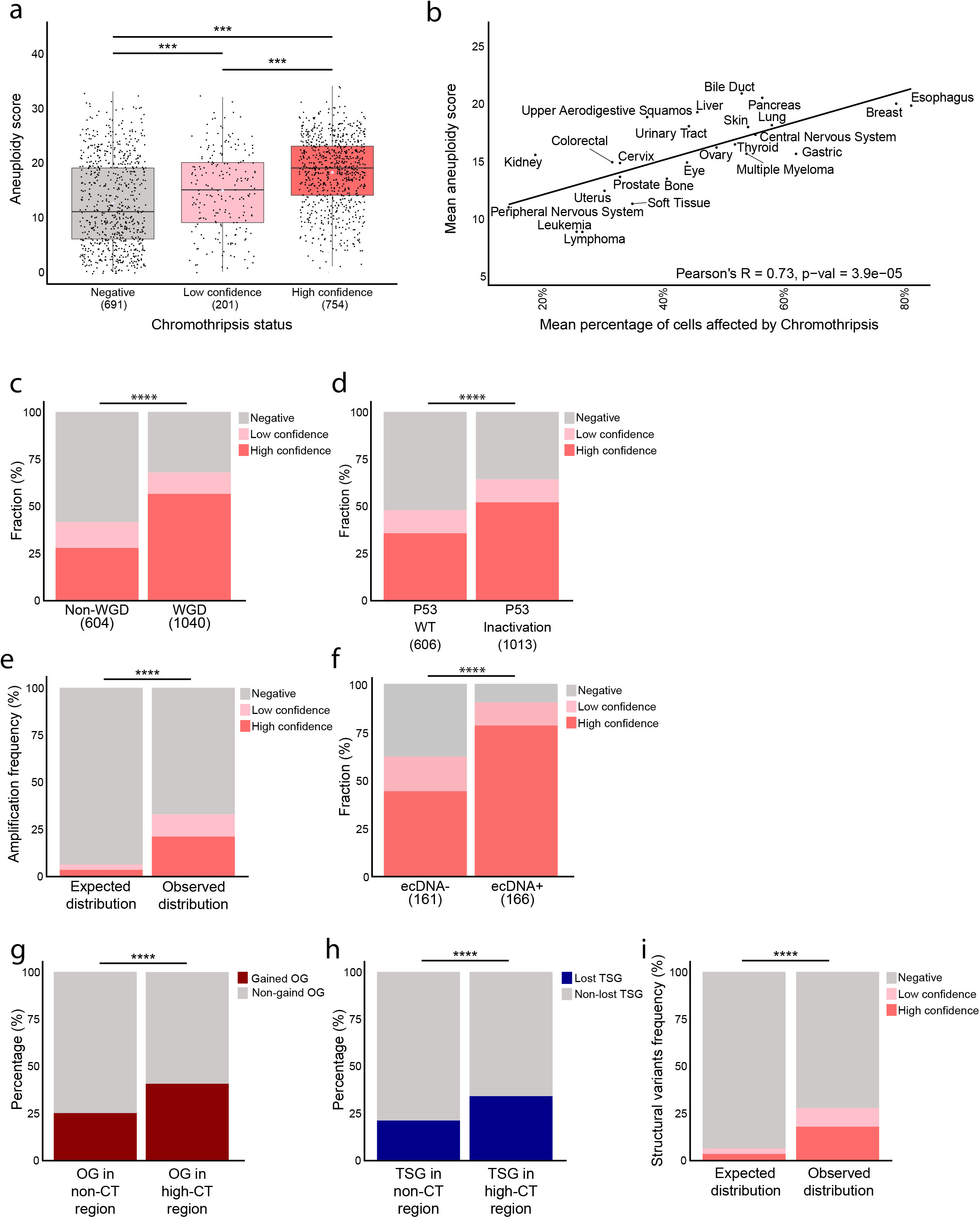
Chromothripsis association with other consequences of CIN. **(a)** Comparison of aneuploidy scores by CT status. High-confidence CT+ cells lines exhibit higher aneuploidy scores than low-confidence CT+ and CT-CCLs (Means: 18.2 vs. 14.9 & 12.6, respectively). **(b)** Correlation between CT prevalence and aneuploidy score across cancer types (Pearson’s R = 0.73, p-value = 5.7e-05). **(c),** Comparison of CT prevalence between WGD+ and WGD-CCLs. CT prevalence is significantly higher in WGD+ vs. WGD-CCLs (56.4% vs. 27.6%, Chi-square test, p-value = 6.5e-30). **(d)** Comparison of CT prevalence between *TP53*-deficient and *TP53*-proficient CCLs. CT prevalence is significantly higher in TP53-deficient cells lines (51.9% vs. 35.5%, Chi-square test, p-value = 9.5e-11). **(e)** Comparison of the expected vs. observed prevalence of focal amplifications within CT events. CT events are strongly enriched for the presence of focal amplifications, compared to the expected prevalence if the two phenomena were independent (Chi-square test, p-value < 0.0001). **(f)** Comparison of CT prevalence between ecDNA+ and ecDNA-CCLs. CT prevalence is significantly higher in ecDNA+ vs. ecDNA-CCLs (78% vs. 44%, Chi-square test, p-value = 1.53e-10). **(g)** Comparison of the fraction of OGs that are gained within vs. outside CT events, demonstrating a strong enrichment for OG gain within CT events (Chi-square test, p-value < 0.0001). **(h)** Comparison of the fraction of TSGs that are lost within vs. outside CT events, demonstrating a strong enrichment for TSG loss within CT events (Chi-square test, p-value < 0.0001). Boxplots: bar = median; box = 25-75th percentile; whiskers = range. **(i)** Comparison of the expected vs. observed prevalence of SVs within CT events. CT events are strongly enriched for the presence of SVs, compared to the expected prevalence if the two phenomena were independent (Chi-square test, p-value < 0.0001).

A positive correlation between mean CT prevalence and mean aneuploidy prevalence exists across cancer types (**Fig. 3b**, **Extended Data Fig. 11a**; Pearson’s R = 0.73, p-value = 5.7e-05). However, specific cancer types, such as bile duct, kidney, and pancreas do not demonstrate this association between CT and aneuploidy (**Extended Data Fig. 11a**; we note, however, that some cancer types include a small number of CCLs, reducing statistical power). Interestingly, at the chromosome level, we observed only a weak, statistically insignificant positive correlation between the prevalence of aneuploidy of a given chromosome and the prevalence of its chromothripsis in the same cancer type (**Extended Data Fig. 11b**; Pearson’s R = 0.302, p-value = 0.172), suggesting that the two phenomena may be alternative means for achieving the same outcome. However, at the individual cell line level, CT is significantly more common in aneuploid chromosomes (**Extended Data Fig. 11c**; paired *t* test, p-value = 1.06e-10). For example, chromosome-arm 1p is three times more likely to undergo CT when it is aneuploid than when it is diploid (10.4% vs. 3.7%, Chi-square q-value = 2.92e-06; **Extended Data Fig. 11d**). Overall, the strong association between CT and aneuploidy is in line with the important role of chromosome mis-segregation in the generation of both types of chromosomal aberrations^6,7^.

Next, we assessed whole-genome doubling (WGD; defined as genome-wide ploidy >2.5, as previously used in Voronina et al.^18^). CT-like events are much more common in WGD+ compared to WGD-CCLs (**Fig. 3c**; 56.4% vs. 27.6%, respectively, Chi-square, p-value < 2e-16). A plausible explanation for that would be that WGD can buffer the consequences of CNAs, allowing more tolerance towards chromothripsis^49,50^. We previously showed a similar positive relationship between WGD and aneuploidy in human CCLs^51^.

Various CIN manifestations are known to be associated with perturbations in the *TP53* tumor suppressor gene. We therefore assessed the association between CT and *TP53* inactivation^21^. Whereas TP53 perturbation is not necessary for CT, the prevalence of CT in *TP53*-mutated CCLs was significantly higher than that of CCLs with wild-type *TP53* (**Fig. 3d**; 51.9% vs. 35.5%, respectively, Chi-square, p-value = 9.5e-11). This trend was observed across cancer types (**Extended Data Fig. 12a**), but reached significance only in 3 of them, likely because the p53 pathway can be perturbed in various ways. A notable example is gastric cancer, in which 89.7% of the high-confidence CT+ CCLs were *TP53*-perturbed in comparison to only 30.0% of the CT-CCLs (p-value = 0.0009). We note that this association is expected, as *TP53* perturbation disrupts cell cycle checkpoints, allowing cells to keep proliferating despite substantial problems in replication and/or mitosis^52^. This observation is in line with previous findings in patient tumors identifying CT in 38% of the samples with *TP53* inactivation, compared to 24% of the samples with a wild-type *TP53*^17^.

Chromothripsis has been shown before to lead to focal amplifications, and to give rise to extrachromosomal DNA (ecDNA) and homogeneously staining regions (HSRs)^51^, both of which lead to DNA segment amplification and result in high-level CNAs. To study the association between CT and high-level amplifications, we followed the approach outlined by Cortés-Ciriano et al. (2020)^17^, defining high-level amplifications as regions with more than 10 copies, flanked by low copy number regions (**Methods**). We identified 2,240 amplified regions across 755 (45.87%) of the CCLs (**Supplementary Table 4**). Interestingly, high-level amplifications were significantly more prevalent within CT+ regions (**Fig. 3e**; Chi-square, p-value = 1.35e-112). To further investigate whether this association extends specifically to ecDNA, we leveraged AmpliconArchitect^53^-derived ecDNA calls from WGS data. ecDNA+ CCLs were significantly enriched for high-confidence CT in comparison to ecDNA-CCLs (**Fig. 3f**; 78% vs. 44%, respectively, Chi-square, p-value = 1.53e-10). These strong associations are consistent with the known consequences of CT, which involves extensive chromosomal fragmentation and rearrangement, providing a genomic environment conducive to the formation of ecDNA and focal amplifications^17–19^.

Next, we assessed the associations between CT and oncogene (OG) gains or tumor suppressor gene (TSG) losses. Specifically, we asked whether CT-like events would encompass *bona fide* cancer genes more than would be expected by chance. We found that both OG gains and TSG losses were highly enriched within CT+ regions (**Fig. 3g,h**; Chi-square, p-value < 0.00001), suggesting that CT-like events indeed facilitate OG amplification and TSG deletion. This pattern is consistent with the strong selection pressures that determine the fate of CT+ cells^2,3^.

Lastly, structural variants (SVs) such as translocations and inversions are also key manifestations of CIN, often reflecting major disruptions in mechanisms that preserve genome integrity. These events can rewire regulatory landscapes, create gene fusions, or isolate genomic segments, contributing to oncogenic processes. Indeed, we found a strong enrichment for SVs within CT+ regions (**Fig. 3i**; Chi-square, p-value < 0.00001), consistent with the structural complexity that characterizes CT^2,3,17^.

### Comparison of chromothripsis landscapes between human tumors and cell lines

A major question pertaining to CT in human CCLs is the extent to which they resemble CT in human tumors. We thus sought to compare the genomic landscape of CT-like events that we observed in the CCLs to that previously reported in patients’ tumors (the two seminal studies of Cortés-Ciriano et al. and Voronina et al.)^17,18^.

The prevalence of CT-like events across human CCLs is 46% (**Fig. 2a**), compared to 29% and 49% in tumors (in Cortés-Ciriano et al. (2020) and Voronina et al. (2020), respectively)^17,18^. These differences could potentially represent differences in the cohort composition. We therefore performed a matched analysis of CT prevalence, considering all cancer types for which >5 samples were available both in the cell line cohort and in the tumor cohort. This analysis confirmed that, across cancer types, the prevalence of CT-like events is higher in CCLs in comparison to tumors: 48.7% in CCLs vs. 31.0% in tumors based on the Cortés-Ciriano et al. study; or 56.8% in CCLs vs. 43% of tumors based on the Voronina et al. study (**Fig. 4a**, **Extended Data Fig. 13a**)^17,18^. The prevalence of CT-like events across cancer types positively correlates between CCLs and tumors (**Extended Data Fig. 13b,** Pearson’s R = 0.604, p-value = 0.0011 for Cortés-Ciriano et al. and Pearson’s R = 0.50, p-value = 0.056 for Voronina et al.)^17,18^. However, in cancer types with low CT prevalence in tumors, such as Non-Hodgkin Lymphoma (Lympho-BNHL), Chronic Lymphocytic Leukemia (Lymph-CLL) and Pilocytic Astrocytoma (CNS-PiloAstro), CT prevalence tends to be considerably higher in the CCLs (**Fig. 4a**, **Extended Data Fig. 13a**). This can be due to selection bottlenecks during cell line derivation^29^, or due to the *in vitro* genomic evolution of the CCLs (see below), as we previously observed a similar phenomenon for aneuploidy^32,54^. In both tumors and CCLs, the most chromothriptic cancer type was osteosarcoma. Indeed, CT was recently shown to be important for osteosarcoma tumorigenesis^55^.

**Fig. 4.**
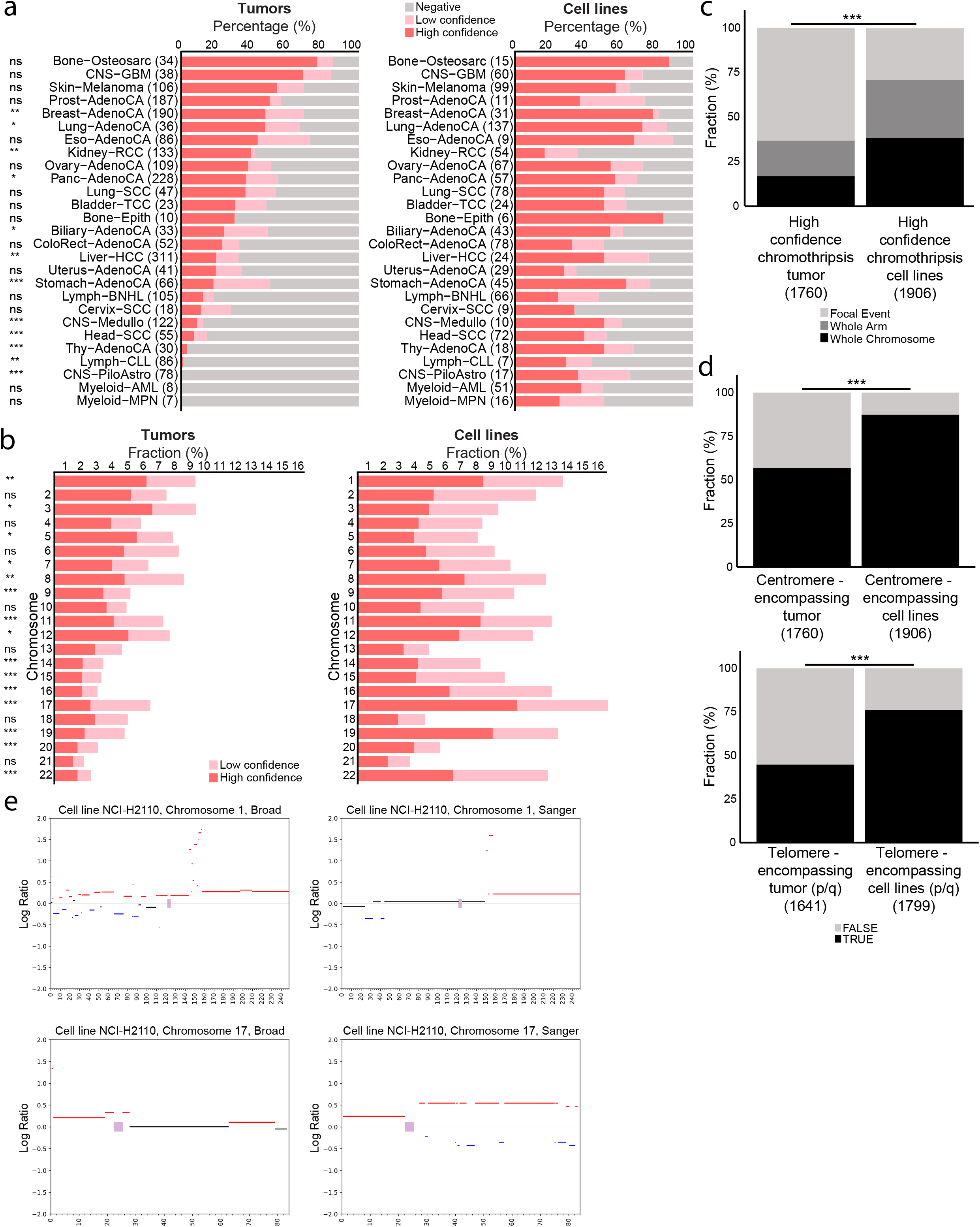
Comparison of CT-like patterns between human tumors and cancer cell lines. **(a)** Comparison of CT prevalence between human tumors (Cortés-Ciriano et al.)^17^ and CCLs across 27 matched cancer types. CT prevalence is significantly higher in CCLs compared to tumors (48.7% vs. 31.0%; p-value <0.001, chi-square test). Cancer types with fewer than 5 samples in either cell line or tumor cohorts were excluded from the analysis. **(b)** CT prevalence across chromosomes in tumors vs. CCLs. Most chromosomes are significantly more likely to be CT+ in CCLs vs. tumors (mean 5.26% vs. 3.29%, p-value =0.011). **(c)** Comparison of the fractions of whole-chromosome, chromosome-arm, and focal CT-like events in tumors (Cortés-Ciriano et al.)^17^ vs. CCLs. CCLs exhibit a higher relative prevalence of whole-chromosome and chromosome-arm CT-like events (38.5% vs. 16.8% and 32.4% vs. 19.9%, for whole-chromosome and chromosome-arm fractions, respectively). **(d)** Comparison of the fraction of CT-like events involving the centromeres (left) and the telomeres (right) in tumors (Cortés-Ciriano et al.)^17^ vs. CCLs. CCLs have a significantly higher involvement of centromeres and telomeres (p or q) in their CT-like events, 87.3% vs. 56.7% and 76.0% vs. 44.7, respectively; p-value <0.001). **(e)** CN landscapes for chromosomes 1 and 17 of the NCI-H2110 (ACH-000929) cell line from Broad^21^ (left) and Sanger^38^ (right) WES-based segmented data. Log2 CN ratios are shown, with segments colored by their ploidy (baseline ploidy in black), gains in red), and losses in blue). Clustered oscillating CN states consistent with CT-like events are present in one dataset but absent in the other, indicating genomic evolution of the CT landscapes during *in vitro* propagation of CCLs.

We note that while CCLs are more CT+ than tumors for most cancer types, a couple of notable exceptions do exist: renal cell carcinoma (Kidney-RCC), which shows a significantly higher CT incidence in tumors compared to CCLs (**Fig. 4a**; 39.1% vs. 16.7%, p-value = 0.003), and soft-tissue sarcoma (**Fig. 4a**, **Extended Data Fig. 13a**; 55% vs. 37%, p-value = 0.22).

Next, we compared the chromosomal prevalence of CT-like events between CCLs and tumors. The most CT+ chromosomes in tumors (chr1, chr12, chr17) also exhibited high CT prevalence in CCLs, and the acrocentric chromosomes were less likely to be CT+ in both data sets (**Fig. 4b**)^17^. However, in line with the higher prevalence of CT-like events in CCLs, many of the smaller chromosomes also exhibited high prevalence of CT in CCLs (but not in tumors), so overall the chromosome-specific CT prevalence was not significantly correlated between tumors and CCLs (**Extended Data Fig. 13c,** Pearson’s r = 0.158, p-value = 0.48). Furthermore, the number of chromosomes involved in CT events was higher in CCLs compared to tumors (**Extended Data Fig. 13d**, p-value = 0.046), and the fraction of samples in which CT involved >5 chromosomes was higher as well (9.41% in CCLs compared to 7.08% in tumors)^17^. Interestingly, however, some common CT events in tumors are not recurrently found in the CCLs. A prominent example is that of chromosome-arm 3p in kidney RCC: chr3p CT was shown to be a main driver of the loss of key TSGs, such as VHL and PBRM1^56^, in kidney-RCC. This recurrent chr3p CT is completely absent from the cell line cohort (**Fig. 2h**), potentially also explaining the lower overall CT incidence observed in kidney-RCC CCLs (in comparison to RCC tumors).

We next compared the chromosomal span of high-confidence CT-like events. CT-like events in CCLs were more likely to encompass an entire chromosome or chromosome-arm: 38.5% and 32.4% of events in CCLs encompass a whole chromosome and a whole chromosome-arm, respectively, compared with 16.8% and 19.9% of events in tumors (**Fig. 4c**, chi-square, p-value < 0.001). While this difference is consistent with the increased CT prevalence in CCLs vs. tumors, we note that it could also be affected by methodological differences (as CT events in the tumors were called by an analysis of WGS, resulting in higher resolution of CT boundaries)^17^. We then compared the positions of high-confidence CT-like events relative to centromeres and telomeres. CT-like events in CCLs tend to encompass the centromere and the telomere more often than tumors: in tumors, 56.7% of all high-confidence events encompass the centromere and 44.7% encompass the (p or q) telomere, compared to 87.3% and 76.0% (respectively) in CCLs (**Fig. 4d**, chi-square, p-value < 0.001)^17^

### *In vitro* evolution of chromothripsis landscapes

Clonal evolution during *in vitro* propagation of CCLs, due to ongoing CIN and genetic drifts, can lead to changes in their aneuploidy landscapes^32^. Whether the landscape of more complex chromosomal events, such as CT, also evolves during culture propagation, remains an open question. To investigate this, we inferred CT-like events in 998 matched CCLs, which were profiled by WES both at the Sanger institute^38^ and at the Broad Institute^21^, and compared the CT landscapes between these matched samples (**Methods**, **Supplementary Table 8**). Our pipeline identified CT-like events in 347 Broad CCLs spanning 493 chromosomes, versus 143 Sanger CCLs spanning 164 chromosomes, with overlap across 132 CCLs and 126 specific chromosomes (corresponding to 92.3% of the CCLs and 76.8% of the events identified in the Sanger data; **Extended Data Fig. 14a,b**).

The higher prevalence of CT detection in the Broad dataset might reflect, at least in part, systematic differences in CN inference, as the two datasets were segmented using distinct pipelines, resulting in the identification of more CN segments and more CN alterations in the Broad dataset (**Extended Data Fig. 14c,d**). However, manual inspection of discordant cases revealed multiple chromosomes with qualitatively distinct CN landscapes between datasets, including localized clusters of oscillating CN states present in one source and absent from the other (**Fig. 4e**, **Extended Data Fig. 14e-h**). These chromosome-level discrepancies cannot be explained by segmentation differences alone and indicate genomic evolution of CT-like events during *in vitro* propagation of CCLs, extending prior observations of dynamic genome evolution in cell line models^32^. This genomic evolution provides another mechanistic basis for variability among nominally identical CCLs across laboratories, and likely contributes to the increased prevalence of CT in CCLs in comparison to that of tumors, and it may therefore have important implications for the *in vitro* modeling of CT^32^.

### Gene expression and cellular vulnerabilities associated with chromothripsis

To identify gene expression programs and cellular vulnerabilities associated with CT, each cell line was assigned a CT score, defined as the fraction of the genome affected by CT. Utilizing these scores, we performed a comparative analysis of gene expression and functional dependencies between CCLs with high-CT scores and those with a CT score of 0 (i.e., CT-negative CCLs). To isolate the effect of CT from the effects of factors that are strongly associated with it, our analysis accounted for potential confounders, including the cancer type, *TP53* inactivation status, aneuploidy score, ecDNA-like amplifications, and the overall ploidy of each sample (**Methods**).

We defined high-CT CCLs as those at the top quartile of CT scores (corresponding to >4% of the genome affected by CT), and conducted differential gene expression analysis between high-CT and CT-negative (CT-) CCLs (**Extended Data Fig. 15a**). We identified 3,922 genes (20.4%) as significantly differentially expressed between the groups (**Supplementary Table 5**). We then performed gene set enrichment analysis (GSEA)^57,58^ based on fold-changes in gene expression, and identified 371 pathways (24.5%) that were significantly enriched (**Supplementary Table 6**). Among the top 50 significantly downregulated pathways, we observed translation-related processes and certain p53-associated pathways (**Fig. 5a**). In contrast, several of the top 50 significantly upregulated pathways were linked to enhanced DNA damage response (DDR), DNA replication, and activation of the proteasome pathway (**Fig. 5a**, **Extended Data Fig. 15b-d**). These changes are consistent with CT cells harbouring high levels of DNA damage, concomitant with a perturbed p53 pathway activity^10,59^. The elevation in the DDR is a possible mechanism to cope with the DNA damage involved in CT; these findings are in line with studies showing that CT is associated with widespread DNA damage, triggering the activation of DDR pathways^10,60^, as well as with our recent findings from highly-aneuploid cells^61,62^. The elevated replication activity observed in high-CT samples might be linked to their increased aggressiveness and enhanced proliferative capacity^17,18,59^.

**Fig. 5.**
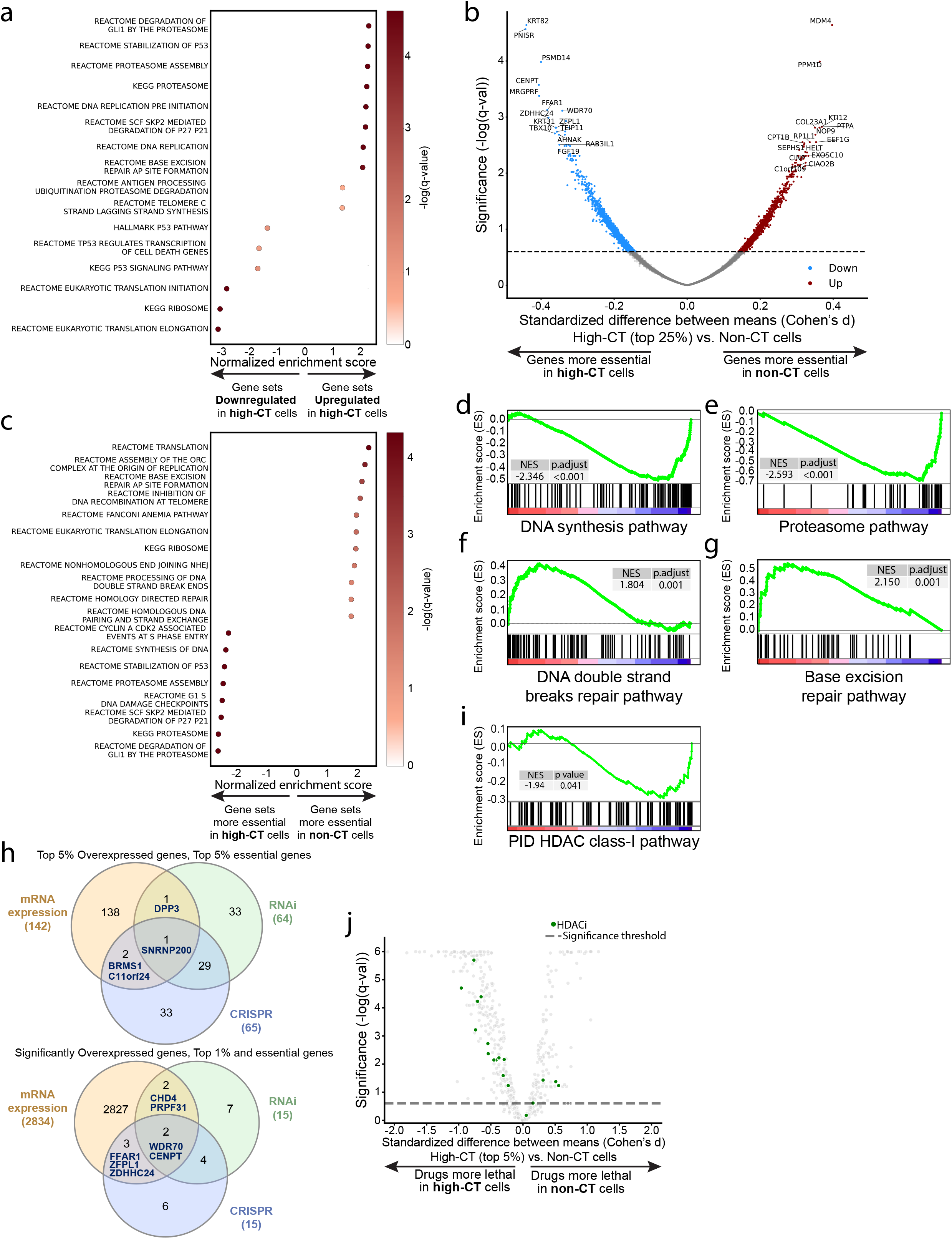
Gene expression programs and cellular vulnerabilities associated with CT. **(a)** GSEA^57,58^ of differentially expressed genes in high-CT vs. CT-CCLs, highlighting pathways among the top 50 most significantly upregulated and downregulated. Enriched pathways include those associated with *TP53*, DDR, proteasome, and ribosome functions. The x-axis shows normalized enrichment score (NES), with significance indicated by dot color **(b)** Volcano plot of the differential sensitivity to CRISPR-mediated gene knockdown. The x-axis shows the standardized difference in mean dependency scores between high-CT (top 25%) and CT-cells, with lower values indicating greater gene essentiality in high-CT cells, while the y-axis shows statistical significance (-log(q-value)), **(c)** GSEA of differential response to knockout in high-CT vs. CT-CCLs, highlighting pathways among the top 50 most significantly enriched and depleted. Enriched pathways include those associated with *TP53*, DNA damage response (DDR), proteasome, and ribosome functions. The x-axis represents the normalized enrichment score (NES), with significance indicated by dot color. **d-g,** Examples of GSEA plots on the ordered gene list, according to the fold change in essentiality score, **(d) ‘**DNA Synthesis’ gene set (NES = -2.346, FDR q-value < 0.001), **(e)** ‘Proteasome’ gene set (NES = -2.593, FDR q-value < 0.001), **(f)** ‘DNA DSB repair’ gene set (NES = 1.804, FDR q-value = 0.033), **(g)** ‘Base excision repair’ gene set (NES = 2.150, FDR q-value = 0.001). **(h)** Top: Venn diagram showing overlap of significantly 5% top overexpressed genes and the top 5% most essential genes from CRISPR and RNAi screens in high-CT vs. non-CT CCLs. Shared genes include *SNRNP200* (three modalities overlap), *BRMS1* and *C11orf24* (mRNA and CRISPR), and *DPP3* (mRNA and RNAi). Bottom: Venn diagram showing overlap of significantly overexpressed genes with the top 15 most essential genes (∼1%) in CRISPR and RNAi screens in high-CT vs. non-CT CCLs. The *WDR70* and *CENPT* genes shared across the three modalities, **(i)** GSEA of the ‘PID_HDAC_CLASSI_PATHWAY’ gene set, according to the fold change in CRISPR-based essentiality score (NES = -1.94, p-value = 0.041), **(j)** Volcano plots of differential drug sensitivity contrasting high-CT (top 5% in CT score) and non-CT cells tested in the CTD² drug screen^28–31^; HDAC inhibitors are marked in green. HDAC inhibitors are overrepresented among drugs that are significantly more lethal to high-CT cells, comparing HDAC inhibitors vs. all other drug classes (Fisher’s exact test, p-value = 0.021).

Next, we utilized RNAi and CRISPR screens^23–27^ to conduct a genome-wide comparison between high-CT and CT-CCLs, in order to identify differential vulnerabilities. Similar to the differential gene expression analysis, we compared the essentiality scores of each gene between the high-CT and CT-CCLs. The essentiality (=dependency) score represents the viability of the cells following gene perturbation (i.e., knockdown in the RNAi screen or knockout in the CRISPR screen); the lower the score, the more sensitive this cell line is to perturbation in that gene. **Fig. 5b** shows the differential response analyses of the CRISPR screens, and **Extended Data Fig. 16a** shows the differential response analyses of the RNAi screens. 2,552 (14.68%) and 3,839 (21.31%) genes exhibited significantly differential dependencies in the CRISPR and RNAi screens, respectively (**Supplementary Table 5**). We then performed GSEA^57,58^ based on the fold-change in gene essentiality, and identified 382 (24.44%) and 412 (27.6%) significantly enriched pathways, for the CRISPR and RNAi analyses, respectively (**Supplementary Table 6**). As expected, among the genes more essential in CT-CCLs were major regulators of the p53 pathway, including MDM4 (**Fig. 5c** and **Extended Data Fig. 16b,c**). In contrast, among the top 50 significantly differential essential pathways, we observed that high-CT CCLs are particularly sensitive to the knockdown and knockout of replication-related genes (**Fig. 5c,d** and **Extended Data Fig. 16b,c**). This finding aligns with the increased replication gene expression observed in CT cells, as well as their high level of DNA damage. Moreover, high-CT CCLs were also more sensitive to the genetic perturbation of proteasome-related genes (**Fig. 5c,e**, and **Extended Data Fig. 16b,d**), likely due to proteotoxic stress induced by the stoichiometric imbalance that results from CT. We note that we previously found that highly-aneuploid cells are similarly more sensitive to proteasome inhibition^62^.

Interestingly, high-CT CCLs exhibited bidirectional trends in response to DDR perturbation, being more sensitive to the perturbation of some DDR-related genes and more resistant to others (**Fig. 5c**, **Extended Data Fig. 16b**). Specifically, replication-specific DDR appears to be more essential for high-CT cells, in line with the replication-related gene expression changes (**Extended Data Fig. 17a,b**, **Supplementary Table 6**). In contrast, the high-CT CCLs are more resistant to perturbation of general (non-replication-specific) DDR mechanisms and telomere maintenance (**Fig. 5c,f,g**, **Extended Data Fig. 16b,e,f**, **Extended Data Fig. 17a,b, Supplementary Table 6**), likely due to the elevated expression of DDR mechanisms that buffer against CT-induced genomic instability.

To validate our findings, we repeated the transcriptomic and functional vulnerability analyses using an independent classification of CT status based on the WGS-based ShatterSeek^17^ calls across 325 CCLs (**Methods**). The differential gene expression analysis (**Extended Data Fig. 18a**) identified a similar number of significantly differentially expressed genes compared to our primary analysis (4,610 genes; 23.98%). GSEA^57,58^ revealed trends similar to those observed in the larger cohort, with enrichment for proteasome (**Extended Data Fig. 18b**), DNA repair (**Extended Data Fig. 18c**), and DNA replication pathways (**Extended Data Fig. 18d**), although these associations did not reach statistical significance and were notably weaker than in the main dataset (likely due to the much smaller sample size). We next compared gene essentiality profiles between high-CT and CT-CCLs using genome-wide RNAi and CRISPR screening data^23–27^ (**Extended Data Fig. 18e-f**). These differential analyses identified substantially fewer significant gene dependencies: 1,405 genes (7.8%) in the RNAi screen, and 239 genes (1.37%) in the CRISPR screen. However, GSEA^57,58^ continued to highlight increased vulnerability of high-CT cells to perturbation of replication-related genes in the RNAi screen (**Extended Data Fig. 18g**), albeit with attenuated significance. Similarly, proteasome-related genes also remained significantly enriched in the CRISPR screen (**Extended Data Fig. 18h**), consistent with the analysis of the larger cohort. Overall, while individual signals were less statistically significant in this much smaller cohort, this analysis confirmed the key transcriptomic and dependency features associated with chromothripsis.

### Chromothripsis is associated with increased sensitivity to HDAC inhibitors

To identify additional dependencies associated with CT, we integrated the differential gene expression analyses with the differential essentiality analyses, focusing on the genes that were consistently identified across multiple datasets (**Fig. 5h**; two intersections were performed, one emphasizing differential expression and the other emphasizing differential essentiality; see **Methods**). This intersection highlighted 11 genes that were differentially expressed and differentially essential in CT+ cells compared to CT-cells, including genes involved in DDR (*CHD4*^63^, *WDR70*^64^), chromosome segregation (CENPT^65^) and chromatin regulation (*CHD4*^63^, *WDR70*^64^, *BRMS1*^66,67^) (**Extended Data Fig. 19**). These analyses align well with key phenotypes observed in high-CT cells, as described above.

Chromothripsis generates extensive structural alterations that can perturb chromatin organization and regulatory interactions^7,19,40,43^. Furthermore, the formation and subsequent evolution of chromothripsis is associated with error-prone and altered DDR^6,11,12,15,60^. HDACs regulate chromatin state and also modulate DDR capacity, and previous studies have shown that HDAC inhibition can potentiate the effects of PARP inhibition, and compromise repair capacity^68–71^. We therefore hypothesized that HDAC activity might represent a selective dependency of CT+ cells. Indeed, HDAC-related processes were differentially essential in high-CT cells; although the number of available HDAC-related gene sets was limited, the overall trend indicated increased essentiality of HDAC genes in high-CT CCLs. Notably, the genes more essential in high-CT CCLs based on the CRISPR screen were significantly enriched for the ‘*PID HDAC CLASS-I PATHWAY’* (NES = -1.94, nominal p-val = 0.041; **Fig. 5i**), demonstrating that genes encoding core HDAC complex components are preferentially required for the survival of high-CT cells (**Supplementary Table 5**). This result was further supported by analysis of the CTD² drug-response screen^28–31^ (**Methods**, **Supplementary Table 6**), which revealed a consistent overrepresentation of HDAC inhibitors among compounds that were significantly more lethal to high-CT CCLs (Fisher’s Exact Test, p-value = 0.021; **Fig. 5j**). These findings suggest that HDAC-mediated chromatin remodeling may represent a potential selective vulnerability in high-CT cells, consistent with a recent report that found a synergistic interaction between HDAC inhibitors and PARP inhibitors in childhood tumors with CT^72^.

### HDAC inhibition selectively sensitizes chromothriptic cells to DNA damage-inducing agents

To experimentally dissect the functional vulnerabilities that are selectively associated with CT, we turned to a unique *in vitro* system of isogenic CCLs with and without CT^8,39^. These are engineered HCT116 cells into which an extra chromosome (either chr5 or chr18) has been introduced via micronuclei formation; in some cases, these micronuclei rupture, resulting in CT, thereby giving rise to isogenic CCLs that have either undergone CT or have not^8,39^. The use of this system therefore allows to experimentally validate CT-dependent cellular vulnerabilities, and to distinguish them from aneuploidy-dependent cellular vulnerabilities^73^. We performed drug response assays in these cells, focusing on DNA damage inducers, HDAC inhibitors and the combination of both.

First, we confirmed that CT+ cells are differentially resistant to DNA damage-inducing agents, namely the topoisomerase II inhibitor, etoposide, and the topoisomerase I inhibitor, topotecan. Both chromosome 5 and chromosome 18 CT+ cells were significantly more resistant to these drugs compared to diploid CT-cells (**Fig. 6a-h**). Interestingly, trisomy 5 and trisomy 18 CT-cells were generally resistant to these drugs as well, in line with the increased resistance of aneuploid cells to DNA damage-inducing drugs^61^;however, CT+ cells were more resistant to the drugs than their aneuploid CT-counterparts (**Fig. 6a-h**). These results confirm that CT is sufficient to induce such differential resistance, consistent with our observations from the DepMap analysis.

**Fig. 6.**
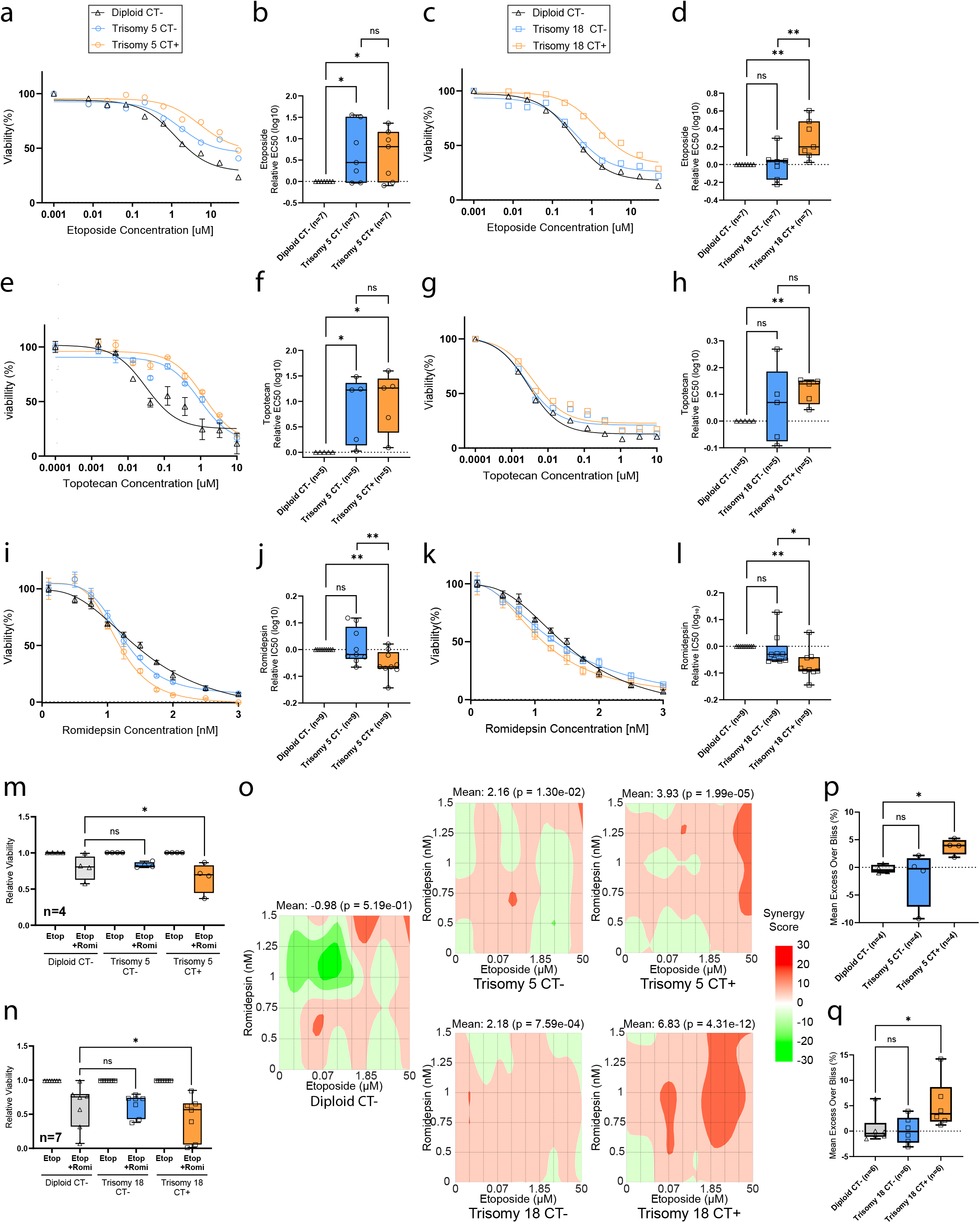
Experimental validation of functional vulnerabilities associated with chromothripsis. a-d, Sensitivity to the Topoisomerase II inhibitor, etoposide, comparing CT- vs. CT+ HCT116 clones. **(a)** Representative dose-response curve for etoposide, in the chr5 isogenic cell line system; X-axis shows the etoposide concentrations ranging from 50 µM to 0.001 µM. Y-axis shows the percentage of viability relative to the untreated controls. The curve was fitted using a non-linear regression model, Inhibitor vs. response (three parameters). **(b)** Comparison of sensitivity, determined by _10_ EC50 values, after 72 hours drug treatment with etoposide, between CT- clones (near-diploid CT- clones and trisomy 5 CT- clones) and trisomy 5 CT+ clones. _10_ EC50 fold change was calculated relative to the diploid CT- clones, per experiment. n = 7 independent experiments; One-tailed one-sample *t* test: *, P = 0.022; *, P = 0.016 for the comparisons of trisomy 5 CT+ vs. diploid CT-, trisomy 5 CT- vs. diploid CT- clones, respectively; Paired one-tailed *t* test : ns, P = 0.42 for the comparison of the trisomy 5 CT+ vs. trisomy 5 CT-clones. **(c)** Representative dose-response curve for etoposide, in the chr18 isogenic cell line system; X-axis shows the etoposide concentrations ranging from 50 µM to 0.001 µM. Y-axis shows the percentage of viability relative to the untreated controls. The curve was fitted using a non-linear regression model, Inhibitor vs. response (three parameters). **(d)** Comparison of sensitivity, determined by _10_ EC50 values, after 72 hours drug treatment with etoposide, between CT- clones (near-diploid CT- clones and trisomy 18 CT- clones) and trisomy 18 CT+ clones. _10_EC50 fold change was calculated relative to the diploid CT- clones, per experiment. n = 7 independent experiments; One-tailed one-sample *t* test: **, P = 0.007; ns, P = 0.5 for the comparisons of trisomy 18 CT+ vs. diploid CT-, trisomy 18 CT- vs. diploid CT- clones, respectively; Paired one-tailed *t* test: **, P = 0.002 for the comparison of the trisomy 18 CT+ vs. trisomy 18 CT- clones. **(e-h),** Sensitivity to the Topoisomerase I inhibitor, topotecan, comparing CT- vs. CT+ HCT116 clones. **(e)** Representative dose-response curve for topotecan, in the chr5 isogenic cell line system; X-axis shows the topotecan concentrations ranging from 10 µM to 0.0001 µM. Y-axis shows the percentage of viability relative to the untreated controls. The curve was fitted using a non-linear regression model, Inhibitor vs. response (three parameters). **(f)** Comparison of sensitivity, determined by _10_ EC50 values, after 72 hours drug treatment with topotecan, between CT- clones (near-diploid CT- clones and trisomy 5 CT- clones) and trisomy 5 CT+ clones. _10_ EC50 fold change was calculated relative to the diploid CT- clones, per experiment. n = 5 independent experiments; One-tailed one-sample *t* test: *, P = 0.011; *, P = 0.023 for the comparisons of trisomy 5 CT+ vs. diploid CT-, trisomy 5 CT- vs. diploid CT- clones, respectively; Paired one-tailed *t* test: ns, P = 0.064 for the comparison of the trisomy 5 CT+ vs. trisomy 5 CT- clones. **(g)** Representative dose-response curves for topotecan, in the chr18 isogenic cell line system; X-axis shows the topotecan concentrations ranging from 10 µM to 0.0001 µM. Y-axis shows the percentage of viability relative to the untreated controls. The curve was fitted using a non-linear regression model, Inhibitor vs. response (three parameters). **(h)** Comparison of sensitivity, determined by _10_ EC50 values, after 72 hours drug treatment with topotecan, between CT- clones (near-diploid CT- clones and trisomy 18 CT- clones) and trisomy 18 CT+ clones. _10_ EC50 fold change was calculated relative to the diploid CT- clones, per experiment. n = 5 independent experiments; One-tailed one-sample *t* test: **, P = 0.00322; ns, P = 0.207 for the comparisons of trisomy 18 CT+ vs. diploid CT-, trisomy 18 CT- vs. diploid CT- clones, respectively; Paired one-tailed *t* test: ns, P = 0.236 for the comparison of the trisomy 18 CT+ vs. trisomy 18 CT- clones. **(i-l),** Sensitivity to the HDAC inhibitor, romidepsin, comparing CT- vs. CT+ HCT116 clones. **(i)** Representative dose-response curve for romidepsin, in the chr5 isogenic cell line system; X-axis shows the romidepsin concentrations ranging from 3 nM to 0.35 nM. Y-axis shows the percentage of viability relative to the untreated controls. The curve was fitted using a non-linear regression model, Inhibitor vs. normalized response - Variable slope. **(j)** Comparison of sensitivity, determined by _10_ IC50 values, after 72 hours drug treatment with romidepsin, between CT- clones (near-diploid CT- clones and trisomy 5 CT- clones) and trisomy 5 CT+ clones. _10_ IC50 fold change was calculated relative to the diploid CT- clones, per experiment. n = 9 independent experiments; One-tailed one-sample *t* test: **, P = 0.006; ns, P = 0.3 for the comparisons of trisomy 5 CT+ vs. diploid CT-, trisomy 5 CT- vs. diploid CT- clones, respectively; Paired one-tailed *t* test: **, P = 0.005 for the comparison of the trisomy 5 CT+ vs. trisomy 5 CT- clones.**(k)** Representative dose-response curve for romidepsin, in the chr18 isogenic cell line system; X-axis shows the romidepsin concentrations ranging from 3 nM to 0.35 nM. Y-axis shows the percentage of viability relative to the untreated controls. The curve was fitted using a non-linear regression model, Inhibitor vs. normalized response - Variable slope. **(l)** Comparison of sensitivity, determined by _10_ IC50 values, after 72 hours drug treatment with romidepsin, between CT- clones (near-diploid CT- clones and trisomy 18 CT- clones) and trisomy 18 CT+ clones. _10_ IC50 fold change was calculated relative to the diploid CT- clones, per experiment. n = 9 independent experiments; One-tailed one-sample *t* test: **, P = 0.003; ns, P = 0.23 for the comparisons of trisomy 18 CT+ vs. diploid CT-, trisomy 18 CT- vs. diploid CT- clones, respectively; Paired one-tailed *t* test: *, P = 0.016 for the comparison of the trisomy 18 CT+ vs. trisomy 18 CT- clones. **(m-n)** Comparison of relative cell viability after 72 hours with a sub-lethal dose (1 nM) of the HDAC inhibitor, romidepsin, or DMSO, with the Topoisomerase II inhibitor, etoposide (1.85 µM), in the chr5 **(m)** and chr18 **(n)** isogenic HCT116 clones. Fold-change calculated relative to DMSO + etoposide treatment. **(m),** Comparison of relative cell viability following combination treatment of sub-lethal doses of romidepsin and etoposide between CT- clones (near-diploid CT- cells and trisomy 5 CT- cells) and trisomy 5 CT+ clones. Relative cell viability was calculated per experiment. n = 4 independent experiments; one-way ANOVA with Dunnett’s correction: *, P = 0.015; ns, P = 0.886 for the comparisons of trisomy 5 CT+ vs. diploid CT-, trisomy 5 CT- vs. diploid CT- clones, respectively. **(n),** Comparison of relative cell viability following combination treatment of sub-lethal doses of romidepsin and etoposide between CT- clones (near-diploid CT- cells and trisomy 18 CT- cells) and trisomy 18 CT+ clones. Relative cell viability was calculated per experiment. n = 7 independent experiments; one-way ANOVA with Dunnett’s correction: *, P = 0.022; ns, P = 0.915 for the comparisons of trisomy 18 CT+ vs. diploid CT-, trisomy 18 CT- vs. diploid CT- clones, respectively. **(o),** Representative 2D heatmaps of Bliss Independence Scores for the combination treatment of romidepsin and etoposide. Representative heatmaps and significance of Mean Excess Over Bliss (EOB) independence scores calculated using SynergyFinder^79^. 2D heatmaps are presented for the *near-diploid CT- cells(Left), Trisomy 5 CT- cells* (top middle), *Trisomy 5 CT+* (top right), *Trisomy 18 CT-* (bottom middle), and *Trisomy 18 CT+* (bottom right). **(p-q)** Comparison of the synergistic effect of the HDAC inhibitor, romidepsin, and the Topoisomerase II inhibitor, etoposide, determined by mean Excess Over Bliss (EOB), after 72 hours of the combination treatment, in the chr5 **(p)** and chr18 **(q)** isogenic HCT116 clones. **(p)** Comparison of the synergistic effect of romidepsin and etoposide between CT- clones (near-diploid CT- cells and trisomy 5 CT- cells) and trisomy 5 CT+ clones. Mean EOB was calculated per experiment. n = 4 independent experiments; one-way ANOVA with Dunnett’s correction: *, P = 0.032; ns, P = 0.768 for the comparisons of trisomy 5 CT+ vs. diploid CT-, trisomy 5 CT- vs. diploid CT- clones, respectively. **(q)** Comparison of the synergistic effect of romidepsin and etoposide between CT- clones (near-diploid CT- cells and trisomy 18 CT- cells) and trisomy 18 CT+ clones. Mean EOB was calculated per experiment. n = 6 independent experiments; Friedman test with Dunn’s correction: *, P = 0.042; ns, P > 0.999 for the comparisons of trisomy 18 CT+ vs. diploid CT-, trisomy 18 CT- vs. diploid CT- clones, respectively. Boxplots: bar = median; box = 25-75th percentile; whiskers = range.

Next, we validated that CT+ cells are differentially more sensitive to romidepsin (**Fig. 6i-l**), an HDAC inhibitor, confirming the preferential essentiality of HDAC complex genes and pathways in CT+ cells. Both trisomy 5 CT+ and trisomy 18 CT+ cells were mildly-but-significantly more sensitive to romidepsin compared to CT-clones (near-diploid CT-cells and trisomy 5/18 CT-cells). Importantly, the trisomic CT-clones did not exhibit differential sensitivity to romidepsin in comparison to diploid CT-cells, highlighting that CT uniquely increases the cellular sensitivity to HDAC inhibition.

Lastly, given that both our computational and our experimental results suggested that HDAC inhibition could be selectively effective against CT+ cells, and that HDAC inhibition was reported to increase the effects of DNA-damaging agents and compromise repair capacity^72^,, we hypothesized that HDACi could sensitize CT+ cells to DNA damage-inducing drugs. To test this, we performed drug combination treatments of HDACi and DNA damage-inducing agents in our isogenic cell system. Notably, HDAC inhibition by romidepsin sensitized CT+ cells to DNA damage-induction by etoposide (**Fig. 6m-q**; **Extended Data Fig. 20**), assessed by the relative cell viability following treatments with a sub-lethal dose of romidepsin in combination with etoposide (**Fig. 6m,n**). Furthermore, the CT-specific synergy between the drugs was confirmed by Bliss Independence analysis (**Fig. 6o,p,q**), as well as LOEWE, Highest Single Agent (HSA), and Zero Interaction Potency (ZIP) synergy analyses (**Extended Data Fig. 20**). Importantly, this sensitization did not occur in CT-cells (either parental HCT116 cells, or trisomic HCT116 cells without CT). These results highlight the importance of further exploring the potential of HDAC inhibition as a therapeutic approach specifically for CT+ tumors, both as a single agent and in combination with DNA damage-inducing agents.

## Discussion

Our study presents the first comprehensive landscape of chromothripsis across human cancer cell lines^21,22^, overcoming the limitation of sparse WGS data by developing and validating a robust copy-number-based detection pipeline. Our findings suggest that copy-number-based chromothripsis detection can complement structural-variation-based chromothripsis detection, not only when WGS data are limited but also when they are available. A key insight from our chromothripsis map is the high concordance between cell lines and primary tumors^17,18^ with regard to lineage-specific prevalence and chromosome-specific patterns of chromothripsis. While we observed a significantly higher prevalence of chromothripsis in CCLs compared to primary tumors^17,18^ – likely reflecting the intense selective pressures of *in vitro* establishment^29^ and/or the ongoing karyotypic evolution in culture^32^ – the conservation of the genomic “fingerprint” confirms that CCLs remain faithful models for studying this phenomenon. This resource allows researchers to move beyond anecdotal case studies and select appropriate *in vitro* models based on a standardized, pan-cancer chromothripsis map.

The strong associations we identified between chromothripsis and other genomic features, particularly aneuploidy, *TP53* inactivation and WGD, underscore the ‘permissive’ environment required to survive such a catastrophic event. The link to *TP53* supports the notion that checkpoint bypass is essential for cells to propagate massive genomic rearrangements rather than undergo apoptosis. Similarly, the enrichment in WGD+ tumors suggests that polyploidy may buffer the deleterious effects of the widespread copy-number losses often associated with chromothripsis^49,50^. Interestingly, 20q – the chromosome arm least frequently affected by chromothripsis in our analysis – is also the most frequently gained chromosome arm across cancers^74^. Together with the distinct associations we observed between chromosome-arm aneuploidy and chromothripsis, this inverse pattern suggests that the selection forces shaping recurrent copy-number gains are not the same as those that render chromosomes susceptible to catastrophic rearrangement .

Crucially, the tight associations between chromothripsis and other consequences of chromosomal instability pose a challenge for studying CTs in model systems (as well as in tumors): controlling for these confounders in our computational analyses, as well as applying an isogenic model^8,39^ to validate the results, were therefore critical in isolating chromothripsis-specific vulnerabilities. Future studies must carefully disentangle the specific consequences of chromothripsis from confounders such as high aneuploidy and p53 loss.

By integrating the CT genomic landscape with functional screening data^23–31^, we uncovered a specific therapeutic vulnerability. While CT+ CCLs displayed a relative resistance to standard DNA-damaging agents – potentially due to a constitutively upregulated DNA damage response (DDR) adaptation – they exhibited a distinct sensitivity to histone deacetylase inhibitors (HDACi). We hypothesize that the extensive chromosomal rearrangements characteristic of chromothripsis impose a severe demand on chromatin remodeling machinery to maintain genomic accessibility and stability. Consequently, HDAC inhibition may disrupt this fragile epigenetic homeostasis. Our validation in isogenic models confirms that HDAC inhibition acts synergistically with DNA damage, effectively re-sensitizing the resistant CT+ cells to treatment with DNA damage-inducing agents.

Our findings extend the previously observed synergy between HDAC and PARP inhibition in the context of CT^72^, suggesting that HDACi may selectively sensitize CT+ cancer cells to various types of DNA-damaging agents. This broader vulnerability of CT+ cancers to the combined disruption of chromatin homeostasis and DNA damage induction highlights a rational combination strategy for this specific genomic context. We note that the exact mechanistic underpinning of the observed synergy should be explored in future studies. As class I HDAC proteins play roles in both HR and NHEJ^75–77^, their inhibition by HDACi may impair the execution of DDR programs, which may be particularly disturbing for high-CT cells. In addition, altered chromatin composition following HDAC inhibition was reported to increase transcription-replication conflicts (TRCs) and result in RNA-DNA hybrid (R-loops) formation, further increasing the DNA-damage burden^78^. Whether the combination of HDACi and DNA damage-inducing agents acts locally (on the chromothriptic region itself) or globally (on the entire genome) also remains an intriguing question for future research. In summary, this study bridges a gap between genomic characterization and functional biology of chromothripsis. We provide the community with a detailed map of chromothripsis in the most widely used model systems, thereby enabling mechanistic studies of this important phenomenon. Moreover, by identifying the interplay between epigenetic modulation and catastrophic genomic rearrangements, we highlight a promising synthetic lethality – the targeting of CT-induced chromatin regulatory dependencies.

## Extended Data Figure Legends

**Extended Data Figure 1. Evaluation of the CT detection pipeline.** (**a**) Comparison of the detected CT-like events between our pipeline, when applied to SNP data, and visual inspection-based calling (used as the “ground truth” for high-confidence and low-confidence predictions). (**b**) Comparison of the detected CT-like events between our pipeline, when applied to WES data, and visual inspection-based calling (used as the “ground truth” for high-confidence and low-confidence predictions). (**c**) Evaluation of the performance of our pipeline: Precision, correct CT+ calls out of all predicted CT+ events; Recall, true CT+ detected out of all actual CT+ events; FP rate, CT-chromosomes misclassified as CT+; FN rate, CT+ chromosomes misclassified as CT-; F1, the harmonic mean of precision and recall, providing a single measure that balances both; CT Chr/CL, the mean number of chromosomes per cell line that were classified as CT+. (**d**) Comparison of the detected CT-like events between the standard CTLPScanner^37^ pipeline vs. our complete pipeline, when applied to SNP data. (**e**) Comparison of the detected CT-like events between the standard CTLPScanner pipeline vs. our complete pipeline, when applied to WES data. (**f**) Copy number plot from CTLPScanner^37^ output from one of the CT+ chromosomes originally described by Stephens et al. (2011)^2^. The x-axis represents genomic location in Mb, and the y-axis represents the CN (normalized to ploidy). Red and blue segments represent amplified and lost genomic fragments, respectively, and the CT event detected by CTLPScanner^37^ is marked in orange. (**g**) Copy number plot of the same chromosome from our post-CTLPscanner code output. The top section provides metadata about the cell line, including disease classification (cancer type and subtype), availability of SNP and WES data, total number of CNAs per available platform, and other CIN manifestations such as aneuploidy score and WGD status. It also details basic features of the detected CT-like event: start/end positions, number of segments and CNAs, and log likelihood ratio (comparing the likelihood of the observed copy-number pattern under a single catastrophic chromothripsis model versus a gradual accumulation model). In the copy number plot, the x-axis represents the genomic location in Mb and the y-axis represents the CN (normalized to ploidy); red, black, and blue segments represent amplification, neutral CN, or loss, respectively, and the detected CT interval is highlighted in green. The bottom section includes information regarding the genomic properties of the CT event, such as centromere/telomere involvement and event scale (whole-arm, whole-chromosome, or focal CT). In this example, CTLPScanner^37^ did not capture the entire event, which our pipeline determined to be a whole-chromosome event. For full reports on all detected CT-like events, please see the ‘Data Availability’ section.

**Extended Data Figure 2. Accurate identification of *bona fide* CT events in human CCLs.** (**a**) CT events identified by our pipeline in 14 human CCLs that were previously analyzed by Stephens et al. (2011)^2^. Each row represents a cell line; the fourth column lists the CT events reported in Stephens et al. (2011)^2^. High-confidence and low-confidence events detected by our pipeline are listed in the fifth and sixth columns, respectively. Chromosome 1 in the SW982 (ACH-001274) cell line is the only CT event in an autosomal chromosome that was missed by our pipeline. Four additional CCLs reported in Stephens et al. (2011)^2^ that had no SNP or WES data, were not included in this analysis. (**b,c**) A CT event detected in chromosome 15 in the ACH-000722 cell line that was reported in Stephens et al. (2011)^2^ and detected by our pipeline. Shown are the CN plot generated using RawCopy^41^ (**b**), and the CN plot and CT detection of our pipeline (**c**).

Extended Data Figure 3. Comparison to ShatterSeek, a WGS-based CT detection algorithm. (**a**) Venn diagram showing the overlap in CT+ chromosomes between our pipeline and the ShatterSeek^17^ algorithm across 325 human CCLs with available WGS data. Only high-confidence CT+ CCLs were considered for this analysis. (**b**) A CN-neutral rearrangement event on chromosome 5 of the HCC70 (ACH-000668) cell line that was classified as CT by ShatterSeek, but was not classified as such by our pipeline due to the lack of CNAs. **(c)** A CT-like event on chromosome 21 of the NCIH1666 (ACH-000448) cell line, lacking major clusters of interleaved SVs. This event was not classified as CT by ShatterSeek, but was classified as such by our pipeline due to its oscillating CNAs. Left: CN and SV profiles from ShatterSeek^17^. Right: outputs from our pipeline.

**Extended Data Figure 4. Example of a visual report of a human lung cancer cell line (CHAGOK1, ACH-000867) with CT-like event in chromosome 8.** (**a**) An overview of the results of all chromosomes in the cell line. (**b**) A visual report of the detected CT+ chromosome. The top section provides metadata about the cell line, including disease classification (cancer type and subtype), availability of SNP and WES data, total number of CNAs per data type, and other CIN manifestations such as aneuploidy score and WGD status. It also details basic features of the CT-like event: start/end positions, number of segments and CNAs, and log-likelihood ratio. In the copy number plot, the x-axis represents the genomic location (in Mb) and the y-axis represents CN (normalized to ploidy); red, black, and blue segments represent amplification, neutral CN, or loss, respectively, and the detected CT-like event is highlighted in green. The middle section includes information about the CN states, including a histogram of segment frequencies at each of the CN states involved in the CT event. The peaks indicate the CN states with peaks exceeding 3% of the total event length; CN states that make for >15% of the total event length are defined as ‘dominant states’. This histogram was smoothened using a Gaussian kernel with a bandwidth of 0.05 to enhance peak identification. The bottom text section includes information from CTLPScanner^37^ (the first step in our pipeline) and genomic properties of the CT event, such as centromere/telomere involvement and event scale (whole-arm, whole-chromosome, or focal CT). (**c**) CN and SV profiles from ShatterSeek visualization. (**d**) CN plot generated using RawCopy^41^.

**Extended Data Figure 5. Genomic Features of CT-like events**. **(a)** Proportions and absolute counts of human cancer cell lines (CCLs, n = 1,646), classified as chromothripsis-positive (n = 757, 46%), borderline (n = 202, 12.3%), or negative (n = 687, 41.7%). (**b**) Fraction of the genome affected (FGA) by high-confidence CT across cancer types. Cancer type is associated with the fraction of the genome affected by CT (ANOVA p-value = 2.01e-08). Bone cancer has the highest FGA (16.6%) whereas cancers of the peripheral nervous system have the lowest FGA (3.1%). (**c**) Density (number of CNAs divided by event length) of 1,419 CT events across 22 lineages derived exclusively from SNP data. Mean = 0.64, Median = 0.51. (**d**) Density (number of CNAs divided by event length) of 474 high-confidence CT events across 22 lineages derived exclusively from WES data. Mean = 0.3, Median = 0.25. Boxplots: bar = median; box = 25-75th percentile; whiskers = 1.5x interquartile range.

**Extended Data Figure 6. Distribution of copy number states in high-confidence CT events across cancer types.** (**a**) Number of copy number states (CNSs) detected in high-confidence CT events across cancer types. A CNS is defined as such if it accounts for at least 3% of the event. The mean number of CNSs is 5. Cancer type is associated with the mean number of CNSs (p-value = 3.82e-07). (**b**) Number of dominant CNSs in high-confidence CT events across cancer type. A dominant CNS is defined as such if it accounts for at least 15% of the event. The mean number of dominant CNSs is 2.5. Cancer type is associated with the mean number of dominant CNSs (p-value = 2.81e-08). (**c**) Comparison of CT prevalence between CCLs derived from primary tumors and those derived from metastases across cancer types. Only a slight increase is observed in metastasis-derived vs. primary-derived CCLs (54.2% vs. 48.7, Chi-square, p-value = 0.233) (**d**) Comparison of CT prevalence between CCLs derived from primary tumors and those derived from metastases across cancer types. No significant association is observed, except for bone cancer (Chi-square test).

**Extended Data Figure 7. Chromothripsis prevalence and co-occurence across chromosomes and chromosome-arms.** (**a**) CT prevalence across chromosome-arms. CT prevalence varies significantly across chromosome-arms (p-value < 2.2e-16). The most chromothriptic chromosome-arm is 17q (10.1%), whereas the least chromothriptic chromosome-arm is 21q (1.94%). (**b**) Comparison of CT prevalence between acrocentric and non-acrocentric chromosomes. Acrocentric chromosomes (13, 14, 15, 21 & 22) are significantly less chromothriptic than non-acrocentric chromosomes (3.7% vs. 5.7% respectively, p-value = 2.53e-12). (**c**) Genomic landscape of 12 chromosomes affected by high-confidence CT in a single ovarian cancer cell line (ONCO-DG-1). (**d**) Distribution of the number of chromosomes affected by CT in the WGS subset of CCLs (n=201), closely resembling the full cohort shown in Fig. **2d** (means: 2.75 vs. 2.52, Mann-Whitney p-value = 0.47). (**e**) Distribution of the number of chromosomes affected by high-confidence CT across cancer types. Cancer type is significantly associated with the number of chromosomes involved in high-confidence CT (One-Way ANOVA, p-value = 8.36e-08).

**Extended Data Figure 8. Correlations between measurements of CT.** (**a**) Prevalence of CT+ CCLs vs. the mean number of chromosomes involved in high-confidence CT, across cancer types. CT prevalence is strongly correlated with the number of involved chromosomes (Pearson’s R = 0.86, p-value = 1.11e-08). (**b**) Prevalence of CT+ CCLs vs. the fraction of the genome affected (FGA) by CT across cancer types. CT prevalence is strongly correlated with the FGA affected by CT (Pearson’s R = 0.98, p-value < 0.0001).

**Extended Data Figure 9. Prevalence of whole-chromosome, chromosome-arm and focal high-confidence CT events across cancer types and chromosomes.** (**a**) Prevalence of whole-chromosome, chromosome-arm and focal CT events across cancer types. Cancer type is significantly associated with the relative prevalence of whole-chromosome events (p-value = 4.1e-07). (**b**) Prevalence of whole-chromosome, chromosome-arm and focal CT events across chromosomes. Chromosome identity is significantly associated with the relative prevalence of whole-chromosome events (p-value < 2.2e-16). For acrocentric chromosomes, whole q-arm events were considered as whole-arm (**c**) Prevalence of chromosome-arm and focal CT events across chromosome-arms. Chromosome-arm identity is significantly associated with the relative prevalence of chromosome-arm events (p-value < 2.2e-16).

**Extended Data Figure 10. Correlations between CT length and chromosome length to the prevalence of whole-chromosome events across cancer types and chromosomes.** (**a**) Correlation between mean CT length and the fraction of whole-chromosome events (out of all CT events) in high-confidence chromothriptic CCLs. Pearson’s R = 0.45, p-value = 0.0255. (**b**) Correlation between chromosome length and the fraction of whole-chromosome events (out of all CT events) in high-confidence chromothriptic CCLs. Pearson’s R = -0.63, p-value = 0.0017.

**Extended Data Figure 11. Chromothripsis association with aneuploidy.** (**a**) Comparison of aneuploidy scores between CT+ and CT-CCLs across cancer types. The aneuploidy score is significantly higher in high-confidence CT+ CCLs in 10 cancer types, and shows a similar trend for almost all cancer types (unpaired t-tests). (**b**) Correlation between chromosome-specific aneuploidy and CT prevalence across all autosomes (Pearson’s R = 0.302, p-value = 0.172). (**c**) Comparison of CT prevalence between aneuploid and non-aneuploid chromosome-arms. CT is significantly more frequent within aneuploid chromosome-arms compared to their non-aneuploid counterparts (paired t-test, p = 1.06e-10). (**d**) An example for a chromosome-arm-specific increase in CT prevalence when that chromosome-arm is aneuploid. Shown is the comparison of CT prevalence on Chr1p when the chromosome-arm is aneuploid vs. when it is not (chi-square, q-value = 2.92e-6). Boxplots: bar = median; box = 25-75th percentile; whiskers = 1.5x interquartile range.

**Extended Data Figure 12. Chromothripsis associations with *TP53* status and with metastasis.** (**a**) Comparison of CT prevalence between *TP53*-WT and *TP53*-perturbed CCLs across cancer types. A trend towards increased CT prevalence in TP53-perturbed CCLs is observed in nearly all cancer types, but reaches statistical significance only in 3 of them (leukemia, lung and gastric cancers; Chi-square test). (**b**) Comparison of the expected vs. observed prevalence of structural variants (SVs) within CT events. CT events are strongly enriched for the presence of SVs compared to the expected prevalence if the two phenomena were independent (Chi-square test, p-value < 0.0001).

**Extended Data Figure 13. Comparison of CT-like patterns between human tumors and CCLs.** (**a**) Comparison of CT prevalence between tumors (Voronina et al.) and CCLs across 15 matched cancer types. CT prevalence is significantly higher in CCLs than in tumors (56.8% vs. 43.0%, p-value < 0.001). (**b**) Correlation of the mean CT prevalence in CCLs vs. tumors across cancer types (Pearson’s R = 0.604, p-value = 0.001). (**c**) Comparison of the number of chromosomes involved in CT-like events in CCLs vs. tumors. The number of chromosomes affected by CT-like events in CT+ CCLs is similar to that of CT+ tumors. (**d**) Comparison of the CT prevalence across chromosomes between CCLs and tumors. The relative prevalence of CT+ chromosomes is not significantly correlated (Pearson’s R = 0.16, p-value = 0.481).

**Extended Data Figure 14. Evidence for *in vitro* evolution of CT-like events in CCLs.** (**a,b**) Comparison between CT-like events detected by our pipeline using Broad^21^ and Sanger^38^ WES-based segmented data, at the chromosome (a) and cell line (b) levels. CT-like events were detected in 347 CCLs spanning 493 chromosomes using the Broad data, versus 143 CCLs spanning 164 chromosomes using the Sanger dataset, with 126 overlapping chromosomes and 132 overlapping CCLs. (**c**) Comparison of the number of CN segments detected per cell line between the datasets, showing a significantly higher number of segments in the Broad data (paired Wilcoxon test, p = 1.69e-131). Boxplots–bar, median; box, 25th–75th percentile; whiskers, range. (**d**) Comparison of the number of CN alterations of adjacent segments per CCL between the datasets, indicating a significantly higher number of alterations in the Broad dataset (paired Wilcoxon test, p = 1.68e-161). Boxplots–bar, median; box, 25th–75th percentile; whiskers, range. (**e-h**) Comparison of CN landscapes between Broad^21^ (left) and Sanger^38^ (right) WES-based segmented data. Panels show (**e**) chromosome 2 of SK-N-DZ (ACH-000366) CCL, (**f**) chromosome 18 of MCF7 (ACH-000019) CCL, (**g**) chromosome 21 of NCIH-1666 (ACH-000448) CCL, and (**h**) chromosome 2 of GOTO (ACH-001716) CCL. Log2 CN ratios are displayed, with segments colored by ploidy (baseline ploidy shown in black, gains in red, and losses in blue). Clustered oscillating CN states consistent with CT-like events are observed in one dataset but not the other, suggesting continued genomic evolution of CT landscapes during *in vitro* propagation.

**Extended Data Figure 15. mRNA expression patterns associated with CT.** (**a**) Volcano plot of the differential gene expression between high-CT cells and non-CT-cells. (**b-d**) Examples of pre-ranked GSEA plots based on the gene list ordered according to the fold change in mRNA expression; (**b**) ‘Protesome’ gene set (NES = 2.227, FDR q-value < 0.001), (**c**) ‘DNA replication’ gene set (NES = 2.097, FDR q-value < 0.001), (**d**) ‘Base excision repair’ gene set (NES = 2.080, FDR q-value < 0.001).

**Extended Data Figure 16. CT-associated responses to RNAi perturbations.** (**a**) Volcano plot of the differential sensitivity to RNAi-mediated gene knockdown. (**b**) GSEA of differential response to knockdown in high-CT vs. CT-CCLs, highlighting pathways among the top 50 most significantly enriched and depleted. Enriched pathways include those associated with *TP53*, DNA damage response (DDR), proteasome, and ribosome functions. The x-axis represents the normalized enrichment score (NES), with significance indicated by dot color. (**c-f**) Examples of pre-ranked GSEA plots based on the gene lit ordered by RNAi essentiality score; (**c**) ‘DNA synthesis’ gene set (NES = -2.912, FDR q-value < 0.001), (**d**) ‘Proteasome’ gene set (NES = -2.782, FDR q-value < 0.001), (**e**) ‘Base excision repair’ gene set (NES = 2.227, FDR q-value < 0.001), (**f**) ‘Nonhomologous end joining repair’ gene set (NES = 2.055, FDR q-value < 0.001).

**Extended Data Figure 17. Stratified analysis of DDR dependencies in high-CT vs. non-CT CCLs.** (**a**) Volcano plot of the differential sensitivity to CRISPR-mediated knockout. (**b**) Volcano plot of RNAi-mediated knockdown. High-CT CCLs are more sensitive than non-CT CCLs to replication-associated DDR perturbations, but are more resistant to general DDR mechanisms. See DDR signature classification in **Supplementary Table 6**.

**Extended Data Figure 18. Transcriptomic features and cellular vulnerabilities associated with CT using WGS-based CT calling (ShatterSeek) across 325 CCLs.** (**a**) Volcano plot of the differential gene expression of high-CT vs. CT-CCLs. (**b-d**) Examples of pre-ranked GSEA on the gene list ordered according to the fold change in expression; (**b**) ‘Proteasome’ gene set (NES = 1.63, FDR q-value = 0.174), (**c**) ‘DNA repair’ gene set (NES = 0.948, FDR q-value = 0.813), (**d**) ‘DNA replication’ gene set (NES = 1.16, FDR q-value = 0.607). (**e,f**) Volcano plot of the differential sensitivity of high-CT vs. CT-CCLS to RNAi-mediated gene knockdown (**e**), and to CRISPR-mediated gene knockout (**f**). (**g,h**) Examples of pre-ranked GSEA on the gene list ordered according to the fold change in essentiality score, in the RNAi screen (**g**) and the CRISPR screen (**h**); (**g**) ‘DNA synthesis’ gene set (NES = -1.468, FDR q-value = 0.165), (**h**) ‘Proteasome’ gene set (NES = -1.800, FDR q-value = 0.040).

**Extended Data Figure 19. CT-specific gene dependencies.** Comparison of high-CT vs. non-CT CCLs for genes that belong to differential pathways emerging from mRNA, CRISPR, and RNAi analyses. Violin plots of mRNA expression (left), CRISPR knockout response (middle), and RNAi knockdown response (right) in high-CT vs. non-CT CCLs for (**a**) *BRMS1,* (**b**) *CENPT,* (**c**) *CHD4* (**d**) *WDR70.* Boxplots: bar = median; box = 25-75th percentile; whiskers = range.

**Extended Data Figure 20. CT-specific synergy between HDACi and DNA damage induction. (a-c)** Comparison of the synergistic effect of the HDAC inhibitor, romidepsin, and the Topoisomerase II inhibitor, etoposide, as calculated by mean LOEWE score (**a**) mean Highest Single Agent score (HSA) (**b**), and mean Zero Interaction Potency (ZIP) (**c**), after 72 hours of the combination treatment, in the chr5 isogenic HCT116 clones. **(a)** Comparison of synergistic effect of romidepsin and etoposide between CT-clones (near-diploid CT-cells and trisomy 5 CT-cells) and trisomy 5 CT+ clones. Mean LOEWE scores were calculated per experiment. n = 4 independent experiments; one-way ANOVA with Dunnett’s correction: *, P = 0.019; ns, P = 0.608 for the comparisons of trisomy 5 CT+ vs. diploid CT-, trisomy 5 CT- vs. diploid CT- clones, respectively. **(b)** Same as in **(a)** but with Mean HSA scores. One-way ANOVA with Dunnett’s correction: ns, P = 0.104; ns, P = 0.569 for the comparisons of trisomy 5 CT+ vs. diploid CT-, trisomy 5 CT- vs. diploid CT- clones, respectively. (**c**), Same as in **(a)** but with Mean ZIP scores. Friedman test with Dunn’s correction: ns, P = 0.068; ns, P > 0.999 for the comparisons of trisomy 5 CT+ vs. diploid CT-, trisomy 5 CT- vs. diploid CT- clones, respectively. **(d-f)** Comparison of the synergistic effect of the HDAC inhibitor, romidepsin, and the Topoisomerase II inhibitor, etoposide, as calculated by mean LOEWE score (**d**) mean Highest Single Agent score (HSA) (**e**), and mean Zero Interaction Potency (ZIP) (**f**), after 72 hours of the combination treatment, in the chr18 isogenic HCT116 clones. **(d)** Comparison of synergistic effect of romidepsin and etoposide between CT- clones (near-diploid CT- cells and trisomy 18 CT- cells) and trisomy 18 CT+ clones. Mean LOEWE scores were calculated per experiment. n = 6 independent experiments; one-way ANOVA with Dunnett’s correction: ns, P = 0.062; ns, P = 0.378 for the comparisons of trisomy 18 CT+ vs. diploid CT-, trisomy 18 CT- vs. diploid CT- clones, respectively. **(e)** Same as in **(d)** but with Mean HSA scores. Friedman test with Dunn’s correction: ns, P = 0.167; ns, P = 0.773 for the comparisons of trisomy 18 CT+ vs. diploid CT-, trisomy 18 CT- vs. diploid CT- clones, respectively. (**f**), Same as in **(d)** but with Mean ZIP scores. Friedman test with Dunn’s correction: *, P = 0.042; ns, P > 0.999 for the comparisons of trisomy 18 CT+ vs. diploid CT-, trisomy 18 CT- vs. diploid CT- clones, respectively.

## Methods

### Ethics

This study utilized established human cancer cell lines and publicly available, de-identified genomic data. No new human biological samples were collected, and no animal experiments were performed in this study. According to the guidelines of Tel Aviv University Institutional Review Board, a specific ethical approval was not required.

### Copy number data assembly

Copy number data files containing 1,745 unique cell lines were downloaded directly from the DepMap portal^23^. One segmented copy number data (.seg) file of 1,040 cell lines originated from Single Nucleotide Polymorphism (SNP) array data, CCLE_copynumber_2013-12-03.seg.txt, as available and described in the Cancer Cell Line Encyclopedia (CCLE)^22^. The second .seg file of 1,604 cell lines originated from Whole Exome Sequencing (WES) CCLE^21^, CCLE_segment_cn.csv 20q2 release. 12 engineered cell lines were omitted from the analysis, resulting in 1,733 cell lines, which included 1,646 CCLs and 87 non-cancerous cell lines. 899 of these cell lines had both SNP and WES CN data available. The segmented CN files underwent the following transformations: the Segment_mean column was log2 transformed and capped at ≤ |3|. Additionally, a Segment_Length column, indicating the length of each segment, was added to meet the input format requirements of the CTLPScanner detector^37^. The segmented CN data file of the Sanger Institute, used for the *in vitro* evolution analysis, (WES_pureCN_CNV_segments_20221214.csv) was obtained from the Sanger Institute Cell Model Passports portal^38^.

### Detection of chromothripsis-like events

CTLPScanner^37^ was run on both SNP & WES .seg files, using the matching human genome builds and the following parameters: number of CNAs in the event ≥ 10, log10 of likelihood ratio ≥ 0, minimum segment size ≥ 10 Kb, and difference in CN called in adjacent segments ≥ 0.3. CTLPScanner output files were then placed into our CT detection post-CTLPscanner code, as extensively described in the Supplementary Methods. Briefly, we reassessed all the positive events provided by CTLPScanner as follows: potential CT+ events were filtered by CN alterations thresholds and a Log10 likelihood ratio score was calculated for assessing the confidence level of the CT. If deemed positive or borderline (“low confidence”), the original CT event coordinates provided by the CTLPScanner were then elongated based on the CNA density observed in proximity to the called event. Importantly, all positive (high-confidence) and borderline (low-confidence) events were subject to visual inspection and corrected as needed. Sex chromosomes were excluded from the analysis.

### Detection of chromothripsis based on whole genome sequencing data

Structural variance (SV) and CN data from 325 CCLs that were previously characterized using WGS^21^ were used as inputs for ShatterSeek^17^. The thresholds for the classification of the events were the default thresholds of the program. P-values associated with ShatterSeek’s statistical tests for CT features (including breakpoint clustering and CN oscillation) were adjusted for multiple testing using the Benjamini-Hochberg false discovery rate (FDR) correction across all chromosomes analyzed. For the analysis of multiple chromosomes in one chromothripsis event, two chromosomes were considered to be part of the same event if they shared at least two SVs within CT regions (at most 10 kb apart), with transitivity applied.

### Aneuploidy, ploidy and omics datasets assembly

Aneuploidy scores, chromosome-arm CNAs, and ploidy for CCLs were obtained from Zerbib et al. (2024)^61^. mRNA gene expression values, CRISPR and RNAi dependency scores (Chronos scores) were obtained from DepMap^23–27^ 22Q2 release. CTD² chemical perturbation screening data^28–31^ were obtained from the DepMap portal^23^ 25Q2 release (Drug_sensitivity_AUC_(CTD^2).csv).

### Matching tumor and cell line cancer types

CCLE lineages were matched to lineages annotated by Cortés-Ciriano et al.^17^ and Voronia et al.^18^ based on biological and histological similarities. Specifically, Plasma lineage was changed to Multiple myeloma, Blood lineage was changed to Leukemia, and Lymphocyte lineage was changed to Lymphoma. CCLs without a match were assigned NA and were excluded from the comparison between tumors and CCLs. The full matching is reported in Supplementary Table 7.

### Chromothripsis associations with focal amplification, cancer genes, and structural variants

To evaluate the association between CT and high-level focal amplifications, high-level amplifications were defined as genomic segments with CN >10 with immediately adjacent flanking regions at baseline chromosomal ploidy, as in Cortés-Ciriano et al. (2020)^17^. The detected high-level focal amplifications are listed in Supplementary Table 4. List of oncogenes (OGs) and tumor suppressor genes (TSGs) were retrieved from COSMIC Cancer Gene Census (CGC)^80^, gene-level CNAs were determined based on the segmented CN data described above. Structural variants (SVs), including translocations and inversions were retrieved from CCLE^21^. For each focal amplification, gained OG, lost TSG, and SV, its location within/outside the CT region was determined. Observed frequencies were compared to genome-wide expectations assuming random genomic distribution, with statistical significance assessed using two-sided Chi-square tests.

### Chromothripsis associations with gene expression and cellular vulnerabilities

CCLs were divided into two groups corresponding to the top and bottom quartiles of the chromothripsis score, defined by the fraction of the genome affected by high-confidence chromothripsis events. For each gene or chemical compound, the log fold-change was estimated in expression, genetic and chemical perturbation between the two groups while adjusting for potential confounders, including cancer type, *TP53* inactivation status, aneuploidy score, ecDNA-like amplification status, and overall ploidy. These confounders were accounted for using inverse probability of treatment weighting (IPTW)^81^, in which the IPTW weights were derived from the samples’ covariates. First, the probability that a sample belongs to the high-CT group (P(high-CT)), defined by scores in the top 25% (gene expression and dependency analyses) and top 5% (chemical compound analysis), was computed. Next, a logistic regression model 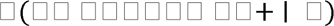 was trained to estimate the probability that a cell line is high-CT given the full confounder set ={cancer type, TP53 inactivation status, aneuploidy score, ploidy, ecDNA-like amplification status}. Each sample’s stabilized weight was then computed as: 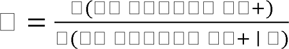. Weighted estimates of gene- or compound-level differences were then obtained using a modified version of Cohen’s d applied to 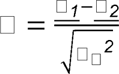, where 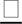 denotes the weighted group mean and 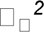 is the weighted pooled variance. P-values of the weighted statistical tests were adjusted for multiple testing using the Benjamini-Hochberg false discovery rate (FDR) correction. The resultant lists of genes, and chemical compounds are available in Supplementary Table 5.

Each type of differential analysis was followed by GSEA, using the GSEA software (version 4.4.0)^57,58^. Four collections of MSigDB^82,83^ gene sets were queried in each analysis: ’Hallmark’^82^, ‘Reactome’^84^, ‘Kegg’^85^, and ‘Biocarta’^82^. The resultant lists of pathways are available in Supplementary Table 6. For the analysis of the HDAC pathway, the following gene sets from the PID database^86^ were used: : ‘PID_HDAC_CLASSI_PATHWAY’, ‘PID_HDAC_CLASSII_PATHWAY’, ‘PID_HDAC_CLASSIII_PATHWAY’ (Supplementary Table 6). The list of the chemical compounds classified as HDACi, appears in Supplementary Table 5.

### Classification of DNA damage response pathways by functional context

DNA damage response (DDR) gene sets were manually grouped into two functional categories based on the biological context in which each repair process is primarily activated. Pathways involved in broad damage sensing, checkpoint activation, double-strand break signaling/repair, nucleotide or base excision repair, and general repair-associated protein regulation were classified as ‘General DDR’, reflecting mechanisms that operate across multiple cell-cycle phases and damage contexts. In contrast, pathways whose activity is tightly coupled to DNA replication, including mismatch repair, DNA damage bypass, repair synthesis/ligation, long-patch base excision repair, and selected homologous recombination-related processes, were classified as ‘Replication-associated DDR’. This categorization was used to distinguish canonical genome-maintenance programs from repair processes more directly linked to replication stress or post-replicative lesion processing.

### Gene-specific analysis

Gene-specific vulnerabilities in high-CT cells were identified using two prioritization strategies integrating differential gene expression^21^ with functional dependency data from CRISPR and RNAi screens^23–27^. In the first strategy, genes were required to be significantly overexpressed and significantly essential, ranking within the top 5% of genes in differential expression modality and at least one of two functional modalities: CRISPR dependency, and RNAi dependency. In the second strategy, genes were required to be significantly overexpressed and significantly essential, ranking among the top 15 most essential genes (approximately the top 1%) in both CRISPR and RNAi screens, independent of expression rank. Statistical significance was defined as an FDR-adjusted q-value < 0.25.

### Cell culture

HCT116 cells (ATCC, RRID: CVCL_0291), and all of their derivative GFP, trisomic and chromothriptic clones were obtained from Kneissig et al.^8,39^. Cells were cultured in DMEM/F-12 (Sartorius) with 15% fetal bovine serum (Sigma-Aldrich), 1% sodium pyruvate, 4mM glutamine, 100 U/ml penicillin and 100 μg/ml streptomycin.

### Drug treatments

Cells were seeded in a 96w plate using Multidrop™ Combi Reagent Dispenser (ThermoFisher). 24h later, cells were treated with drugs of interest. Cell viability was measured after 72h using the MTT assay (Sigma M2128), with 500 μg/mL salt diluted in complete medium and incubated at 37°C for 3h. Formazan crystals were extracted using 10% Triton X-100 and 0.1N HCl in isopropanol, and color absorption was quantified at 570 nm and 630 nm using BioTek Synergy H1 Hybrid Multi-Mode Reader. Absolute IC50 values, and EC50 values were calculated using GraphPad PRISM 9.1, inhibitor vs. normalized response (variable slope) equation. EC50 values were calculated using GraphPad PRISM 9.1, Asymmetric (five parameter), X is concentration equation or inhibitor vs. response (variable slope). Combination treatment interpretation was done using SynergyFinder^79^, calculating mean Excess Over Bliss (EOB), ZIP, LOEWE, and HSA scores. All drugs details are available in Supplementary Table 9.

## Data availability

All the datasets used in the study are publicly available or supplied in the Supplementary Data.

## Code availability

All codes used in this paper are available upon request.

## Acknowledgements

The authors thank Zuzana Storchova for providing the HCT116 euploid, trisomic and chromothriptic CCLs; and the members of the Ben-David and Shamir labs for helpful discussions. Work in the Ben-David lab was supported by the European Research Council Starting Grant (grant #945674 to U.B.-D.), the Israel Cancer Research Fund Project Award (to U.B.-D.), the Israel Science Foundation Project Grant (grant #1805/21 to U.B.-D.), the BSF project grant (grant #2023236 to U.B.-D.) and the Schmidt Science Polymaths Award (to U.B.-D.). Work in the Shamir lab was supported by the Israel Science Foundation (grant No. 2206/22), Len Blavatnik and the Blavatnik Family foundation, and Koret - UC Berkeley - Tel Aviv University Initiative in Computational Biology and Bioinformatics. T.B.-Y. is supported by a fellowship from the Edmond J. Safra Center for Bioinformatics at Tel Aviv University. G.L., A.T., Y.E. and T.W. are supported by fellowships from the Gray Faculty of Medical and Health Sciences at Tel Aviv University. A.T is also supported by a fellowship from the Naomi Prawer Kadar Foundation.

## Author contributions

U.B.-D. conceived the study. T.B.-Y. and G.L. led and performed most of the computational analysis. A.T. led and performed most of the *in vitro* experiments and the data analysis of these experiments. A.T., Y.E., T.W., M.G., L.R., and E.R. assisted with the computational analysis and/or with the manual inspection of CT events. G.L. and Y.E. assisted with the *in vitro* experiments. R.S. contributed to data analysis and supervision. U.B.-D. contributed to experimental design and data analysis, acquired funding, and directed and supervised the overall study. G.L., T.B.-Y., A.T. and U.B.-D. wrote the manuscript with input from all co-authors.

## Additional information

### Supplementary information

#### Supplementary Methods Running CTLPScanner1

CTLPScanner^1^ utilizes a .seg file and a genomic reference build to generate a table of potential CT+ chromosomes. CTLPScanner was executed with maximum sensitivity parameters: number of CNAs in the event ≥ 10, log10 of likelihood ratio ≥ 0, minimum segment size ≥ 10 Kb, and difference in CN called in adjacent segments ≥ 0.3. Segments shorter than 10 Kb and differences between adjacent segments lower than 0.3 are ignored by the statistical analysis performed by the tool. To evaluate the false-negative rate of the results, negative samples were manually assessed and inspected using the CN alteration graphs. 100 samples that were classified as negative by CTLPScanner were examined, and a single sample that could be considered borderline was found. The other samples were true negatives. This indicated that CTLPScanner had a false-negative rate of less than 1% under the lenient parameters we employed. Consequently, any further manual curation was done to samples identified as positive by CTLPScanner. These positive samples were within the CTLPScanner output file, which was then used as an input for our CT detection pipeline.

#### Pipeline for the detection of chromothripsis-like events

##### Pipeline overview

(Supplementary Methods Fig. 1a). CTLPScanner^1^ was run using the sensitive parameters described above, and all intervals identified as CT positive were reassessed. Intervals shorter than 50 Mb or having less than 10 CNAs were re-classified as negative. Subsequently, log likelihood ratio score was used to assess the confidence level of each remaining interval and defined it as positive or borderline (“low confidence”). The original CT interval coordinates provided by CTLPScanner were extended if a sufficient density of CNAs was observed near the interval. After the extension step, all positive and borderline intervals underwent visual inspection and corrections as necessary. The following paragraphs provide a detailed description of our detection method. Throughout this process, we focused on maintaining a very low false positive (FP) rate, at the expense of a higher false negative rate, ensuring that detected CT events were reliable, in order to have higher confidence in downstream analyses .

The input for our detection pipeline consists of (1) segmented CN data (.seg file), (2) the reference human genome build file, and (3) the output table from CTLPScanner, containing in particular the genomic intervals identified as CT+ in each chromosome and for each such interval a partition into segments of constant CN along with their called CN. GRCH37/hg19 was used for the SNP data and GRCH38/hg38 was used for the WES data. Genome builds were imported from the R package RCGH^2^. A preprocessing step is applied to the .seg file to enhance data quality, which involves filtering out segments located within the centromeric regions or the p-arm of acrocentric chromosomes, as these regions cannot be sequenced with high confidence. Additionally, short CN segments were excluded to reduce noise and improve classification accuracy. Specifically, segments shorter than 15,000 nucleotides were filtered out from SNP array data, while segments shorter than 75,000 nucleotides were removed from WES data. These thresholds were empirically determined to suit the characteristics of WES and SNP data segmentation and the different approaches of CN calling algorithms.

Once the data was processed, each chromosome identified as CT-positive by CTLPScanner was analyzed. For the identification of meaningful CNAs in a chromosome, the output of CTLPScanner was used and minor changes in the CN between consecutive segments were disregarded. Specifically, two adjacent segments were considered to have a CNA between them if the difference between their CNs is ≥ |0.3|. The chromosome from the start of every new segment was then scanned and examined whether the window of 50Mb starting at that position contains at least ten CNAs (the threshold suggested by Voronina et al.^3^). The overlapping windows that passed this criterion were then merged and a new larger window called the maximum possible CT interval. In subsequent steps, an attempt was made to elongate the detected CTLPScanner^1^ event within the boundaries of the maximum possible CT interval.

##### Refinement and extension of event intervals

(Supplementary Methods Fig. 1b). Two consecutive steps were used to refine CT event calling. Both rely on considering the density of CNAs in genomic windows. A window’s CNA density (or simply density) was defined as the number of CNAs divided by the window length in 10 MBs units. Hence, for example, a window of 20Mb with 3 CNAs has density 1.5.

Let d denote the density of the CTLPScanner interval. In the first, ‘sliding window’ method, an attempt was made to repeatedly extend the current window by a fixed amount (Supplementary Methods Fig. 2b, top). Two borders were initialized, left and right, at the CTLPScanner event’s boundaries. For each direction, we examine a window of 5% of the chromosome’s length, starting at the current border, and calculate the number of CNAs within that window. If the current window’s density exceeded 0.2d the border to the end of that window was extended. The elongation process stopped when a low density window failed to meet the threshold. This method aimed to address the issue of prematurely terminating an event due to a mildly decreasing number of CNAs in a certain subinterval of the event.

The second, ‘anchored windows’ method, took the output of the sliding window method as its base event. Windows in the size of 10% of the chromosome’s length were appended from each boundary of the base event (Supplementary Method Fig. 2b, bottom). Such windows were repeatedly appended until the border of the maximum possible CT event was reached. Let α be the base event’s density. Windows of density that are ≥0.2α and also ≥4 were searched for. The base interval was extended to include all the range from the leftmost to the rightmost windows that passed this criterion. This method was aimed to overcome events with CNAs distributed bimodally across the chromosome, i.e., two clusters with high CNA density that were separated by low CNA density chromosomal interval. In this scenario, the sliding window method did not elongate the event and did not combine the two far apart clusters. However, the anchored windows method solved this limitation by searching for non-adjacent regions to the primary interval and incorporating them when the CNA density in between met the required threshold.

##### Calling of CN states in the detected event

After the event borders were determined, CN states that reflect current CN levels in the CT landscape were calculated. To determine these states, a histogram of segment frequencies at each CN level was created. The histogram was then smoothed using a Gaussian kernel (using bandwidth of b=0.05 as the normalization parameter), and its peaks identified. Peaks that span throughout >3% of the length of the event (in total) were considered CN states, while those that span >15% of the event were reclassified as dominant CN states.

##### Assessment of chromosomal features at a CT event

Chromosome-arm involvement was determined by setting a threshold of at least 20% of the arm length. The entire chromosome or the whole arm was considered affected by CT if at least 80% of the chromosome or the arm’s length was involved in CT. Additionally, telomeres were determined to be involved if the event’s distance from at least one end of the chromosome was less than 10% of the length of the entire chromosome arm. The centromere was considered involved if the event encompassed the centromere, or if the event’s distance from the centromere was less than 10% of the length of that chromosome arm.

##### Classification of events

After the conclusion of the process, each chromosome with a detected CT event was classified into one of three categories based on our confidence in the detected CT. Events with high certainty were called positives, those with moderate confidence were called borderline, and events highly likely to be negative were classified as negative. Classification was based on several event properties and empirically tested thresholds, which are described below.

A key property considered was the log-likelihood ratio (LLR), which measures the likelihood that a specific genomic interval had undergone CT. This ratio indicates how deviant the alterations frequency in that genomic interval is from a uniform distribution of CNAs across the entire genome of the cell. The LLR was calculated by the formula used by Cai et al. (2014)^4^, suggested originally by Kulldorff (1997)^5^:

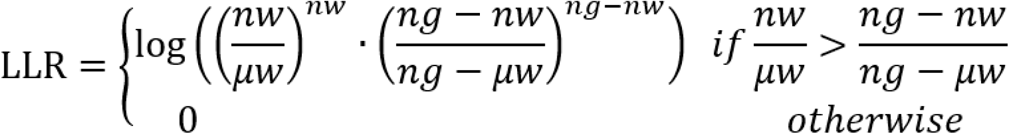

Here □□ is the number of alterations observed in the entire genome, □□ is the number of alterations observed in the specific window, and *μ*□ is the number of alterations expected in that window. *μ*□ is calculated by the formula: 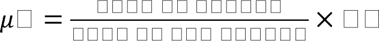.

To classify a chromosome as CT+, the first criterion was that it must contain at least 10 CNAs. Additionally, to avoid capturing large CN changes that represent single gains or losses, with minimal deviation around a single CN state, a requirement from the CT event was to have at least two dominant CN states. Genomic intervals that did not meet these conditions were automatically classified as borderline or negative. Those that did meet the criteria proceeded to the next level of classification. Specifically, events from SNP data that were shorter than 30MB and have 6 CNAs or fewer and a log-likelihood ratio below 0.8 were directly classified as negative.

##### Positive criteria

Chromosomes analyzed based on SNP data were defined as positive (high-confidence CT+) if they met at least one of the following conditions:

1.1. LLR ≥ 8.

1.2. 10Mb ≤ length ≤ 50Mb, and 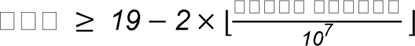

(e.g., LLR≥11 for 40Mb chromosomal-regions and ≥17 for 10Mb regions).

1.3. 30Mb ≤ length ≤ 100Mb, density ≥ 4 (short events with high density).

1.4. Length > 100Mb, density ≥ 3 (long events with medium density).

Chromosomes analyzed based on WES data were defined as positive if they met at least one of these conditions:

2.1. LLR ≥ 4.

2.2 Length ≥ 100Mb, density ≥ 1.6 and LLR ≥ 3 (long events with medium density).

As there were inherent differences in the data characteristics between SNP and WES data, distinct conditions and thresholds were applied for each platform. SNP data contained more segments than WES data, leading to lower CNA density detection in WES data. Therefore, the thresholds on LLR and density were higher for SNP data. Conditions 1.1, 1.2, 2.1, and 2.2 were used for filtering out suspected events that occurred in a genome with a high rate of CNAs, as the LLR value depended on the total number of CNAs in the genome. As long events typically did not maintain high density across their entire length, conditions 1.3, 1.4, and 2.2 provided more lenient LLR and density thresholds for longer intervals.

##### Borderline criteria

Intervals that failed the positive criteria were called borderline (low-confidence CT+) when their CT status was uncertain. These intervals were assessed by manual inspection of CNA plots. Chromosomes analyzed based on SNP data were classified as borderline if they met at least one of the following conditions:

3.1. LLR≥6.

3.2. The original chromosomal region detected by CTLPScanner had LLR ≥ 6.

3.3. The total number of CNAs in the genome was more than 400.

3.4. Before filtering out the noisy segments (short segments), the interval contained > 100 segments.

3.5. The number of CNAs in the event detected by CTLPScanner was at least 1.4 times higher than the number of CNAs in our called interval.

For WES data, samples were defined as borderline if they had a density ≥1.6, and they met at least one of the following conditions:

4.1. LLR ≥ 2.

4.2. The original chromosomal region detected by CTLPScanner had LLR ≥ 4.

4.3. The number of CNAs all over the genome was more than 150.

The conditions outlined above were aimed to strike a balance in our decision regarding the presence of CT in a chromosome. A moderate LLR suggests an event with intermediate confidence of CT (conditions 3.1, 3.2, 4.1, and 4.2). A high rate of CNAs across the entire genome may indicate multiple CT+ chromosomes within a cell. According to conditions 3.3 and 4.3, intervals with at least one 50Mb window with ≥ 10 CNAs were classified as borderline CT if the overall rate of CNAs in the genome was not high enough. An interval that is highly segmented before the pre-processing step with at least one 50Mb window with ≥10 CNAs could not be a confident CT event even if the segments were short, and was thus classified as borderline (condition 3.4). Lastly, condition 3.5 aimed to mark potential CT events for manual examination in which our post-CTLPscanner code’s results and those produced by CTLPScanner differed markedly.

<u>Output</u>. The algorithm generates two outputs: an events table (in .csv format) and a visual report (in .pdf format, see example in Extended Data Fig. 4). The events table contains details about the CT event, such as event coordinates, total number of alterations, and centromere and telomere involvement, alongside other features.

In the visual report, each page presents a visualization corresponding to an event or to a row in the event table. The report includes the data from the table and the histogram used to determine the CN states. The manual curation of events was performed using these visualizations.

Additionally, we execute a supplementary post-processing script to generate a cell line-centric table (.csv). This table provides extensive information about each cell line’s CT status, metadata, and other chromosomal instability (CIN) properties, such as cell type, chromosomes involved in CT, aneuploidy score, type of data used (SNP/WES), and total CNAs in the genome.

##### Final manual modifications

To minimize classification errors and ensure accurate characterization of events, 1,484 positive and 4,352 borderline chromosomes identified by our post-CTLPscanner code were manually curated. Additionally, All 204 events shorter than 15MB were also manually reviewed, including those classified as negative.

As a result of this process, 247 chromosomes (16.6%) were reclassified from positive to borderline and 52 chromosomes (3.5%) were reclassified from positive to negative. In addition, 725 of the borderline chromosomes (16.7%) were reclassified to positive, and 2,289 chromosomes (52.6%) were reclassified to negative. Furthermore, Event boundaries were adjusted for 704 positive events (47.4%), as manual inspection revealed the post-CTLPscanner code was not precise in predicting these boundaries. These modifications were based on visual inspection and applied exclusively to positively called chromosomes.

##### Estimation of error rates

Our pipeline categorized certain CT events as borderline when there was low certainty regarding their CT nature, yet they could not be confidently classified as negative. These events were manually curated after the computational procedure was executed. Performance evaluation was then conducted for three scenarios in which borderline events were considered (1) as a separate category, (2) as part of the positive group, and (3) as part of the negative group.

Our results were based on CN data from both WES and SNP arrays. In a 1Mb genomic interval, it is expected to have approximately 1,000 SNPs and around 50 exons. SNPs tend to be more evenly spread along the genome while exons tend to appear in clusters. Therefore, the results based on SNP arrays were usually of higher resolution and contained more segments from those of WES. Consequently, performance evaluations for the results on SNP and WES data are presented separately.

Chromosomes that were not initially flagged as potential CT cases when CTLPScanner^1^ was run with sensitive parameters were classified as negative and excluded from further analysis by our pipeline. To assess the prevalence of false negatives due to this decision, 200 random chromosomes that CTLPScanner^1^ had classified as negative were visually inspected. This inspection was conducted separately for SNP and WES data, and in each test a single positive CT event was found. Chromosomes called negative by CTLPScanner^1^ were not changed by our pipeline, with a false negative rate of 0.5% based on the visual inspection.

The prevalence of chromosomes that were detected as CT positive by CTLPScanner^1^ and also classified as negative by our post-CTLPscanner code were assessed. For results based on SNP data, 200 random chromosomes were selected for manual inspection. One positive and no borderline chromosomes were identified matching these criteria, for a false negative rate of 0.5% due to such errors.

To estimate error rates of chromosomes that were detected as positive by CTLPScanner^1^ but were classified as negative by our post-CTLPscanner code in WES data, the SNP data was used as a benchmark as it has higher resolution and thus higher sensitivity of CT detection. This was done for both the computational procedure and in visual inspection. 200 negative WES chromosomes that had matching SNP data were randomly selected and visually inspected against the results of the SNP data. Using the results of that visual inspection as the golden standard, this analysis produced six false positive (3%) and 12 borderline events (6%) in the WES data. These numbers were used to extrapolate false negative rates for the entire WES cohort, including samples without matching SNP data.

In summary, based on our sampling, a false negative rate of 0.5% was estimated for all chromosomes classified as negative using SNP data in the entire pipeline. On WES data, for the chromosomes labeled negative, a false negative rate of 0.5% was estimated. For chromosomes labeled positive by CTLPScanner^1^ but classified as negative by our pipeline, A rate of 3% false negatives that were positives and 6% of the negative chromosomes were borderlines was estimated. Based on these calculations, the false negative rate of the entire procedure using WES data was estimated to be 0.55% for chromosomes classified as negative but were actually positive, and 0.12% for those classified as negative but were actually borderline. In all subsequent analyses, these error estimates were relied upon and all the chromosomes classified as negative were not manually reviewed.

##### Performance on SNP data

Extended Data Fig. 1a describes the performance of the pipeline applied to the SNP data compared to the gold standard visual inspection. The visual inspection was performed on all chromosomes classified as positive or borderline by the pipeline. For chromosomes classified as negative, the positive and negative numbers are estimates based on the error rates estimation analyses described above. The performance was measured using precision and recall for each class. For the positive class, the pipeline achieves a precision of 0.8, meaning that most of the samples it predicts as positive are correct, but its recall is lower at 0.55, indicating that it misses many truly positive samples. For the borderline class, precision is much lower at 0.31, but recall is high at 0.87, suggesting the pipeline identifies most borderline cases, though it often misclassifies other classes as borderline.

When the borderline class is considered part of the positive group (combining the first two rows and the first two columns of Extended Data Fig. 1a), our post-CTLPscanner code has precision of 0.56, recall of 0.97, false positive rate of 0.12, false negative rate of 0.03 and F1 score of 0.71. When the borderline is considered part of the negative group (combining the last two rows and the last two columns of Extended Data Figure. 1a), our post-CTLPscanner code has precision of 0.8, recall of 0.55, false positive rate of 0.01, false negative rate of 0.45 and a F1 score of 0.65.

##### Performance on WES data

Extended Data Figure. 1b describes the performance of the pipeline applied for the WES data compared to the gold standard. For the positive class, the pipeline achieves precision of 0.78, indicating that most of the samples classified as positive are correct, and the recall of 0.64. For the borderline class, both precision and recall are very low at 0.32 and 0.38, respectively, indicating substantial difficulty in correctly identifying borderline cases.

When the positives and borderlines are considered one class, our post-CTLPscanner code has precision of 0.91, recall of 0.82, false positive rate of 0.003 and false negative rate of 0.18 and F1 score of 0.86. When the borderline and the negative are considered one group, our post-CTLPscanner code has precision of 0.78, recall of 0.64, false positive rate of 0.01, and false negative rate of 0.36 and F1 score of 0.7.

## Supplementary Methods Figure Legends

**Supplementary Methods Figure 1: Visual description of our pipeline. (a)** Main steps of our CT detection pipeline. Our CT computational pipeline consists of four main steps: (1) filtering based on CNAs, (2) an elongation process, (3) calculation of genomic properties, and (4) event classification. **(b)** Description of the elongation approach applied to each potential CT event. The maximum possible CT interval comprises the union of all the 50Mb genomic intervals along the chromosomes that contain>= 10 CNAs. (top) The sliding window phase: the interval is extended in steps equal to 5% of the chromosome’s length. Extension stops when the density of CNAs gets below the threshold or when the boundary of the maximum possible CT interval is reached. (bottom) The anchored windows phase:regions of 10% of the chromosome length between the ends of the interval output in step A and the maximum possible boundaries, are considered for event inclusion. A region is included in the event if it has sufficient density. The event is then extended throughout the window up to (and including) the furthest positive region on each side. This way, intermediate regions with lower density – which are missed by CTLPScanner – get included in the detected CT-like event.

