## Extended Data Figures for "Mapping chromothripsis across human cancer cell lines reveals genome evolution and therapeutic vulnerabilities"

Extended Data Fig. 1

a

| 1033 non-engineered cancer cell lines |  | SNP pipeline prediction |  |  |  |
| --- | --- | --- | --- | --- | --- |
|  |  | Positive | Borderline | Negative |  |
| SNP visual inspection | Positive | 854 | 602 | 88 | 1544 |
|  | Borderline | 183 | 1274 | 0 | 1457 |
|  | Negative | 33 | 2279 | 17413 | 19725 |
|  |  | 1070 | 4155 | 17501 | 22726 |

b

| 1599 non-engineered cancer cell lines |  | WES pipeline prediction |  |  |  |
| --- | --- | --- | --- | --- | --- |
|  |  | Positive | Borderline | Negative |  |
| WES visual inspection | Positive | 641 | 168 | 187 | 996 |
|  | Borderline | 133 | 106 | 40 | 279 |
|  | Negative | 51 | 56 | 33796 | 33903 |
|  |  | 825 | 330 | 34023 | 35178 |

c

| Data | Classified as positive | Precision | Recall | FP rate | FN rate | F1 | CT Chr / cell line |
| --- | --- | --- | --- | --- | --- | --- | --- |
| SNP | Positive & Borderline | 0.56 | 0.97 | 0.12 | 0.03 | 0.71 | 2.91 |
| SNP | Positive only | 0.8 | 0.55 | 0.01 | 0.45 | 0.65 | 1.49 |
| WES | Positive & Borderline | 0.91 | 0.82 | 0 | 0.18 | 0.86 | 0.8 |
| WES | Positive only | 0.78 | 0.64 | 0.01 | 0.36 | 0.7 | 0.62 |

d

| 1033 non-engineered cancer cell lines |  | CTLPSscanner prediction |  |  |
| --- | --- | --- | --- | --- |
|  |  | Positive | Negative |  |
| SNP visual inspection | Positive | 547 | 909 | 1456 |
|  | Negative+ Borderline | 146 | 21124 | 21270 |
|  |  | 693 | 22033 | 22726 |

e

| 1599 non-engineered cancer cell lines |  | CTLPSscanner prediction |  |  |
| --- | --- | --- | --- | --- |
|  |  | Positive | Negative |  |
| WES visual inspection | Positive | 214 | 595 | 809 |
|  | Negative+ Borderline | 19 | 34350 | 34369 |
|  |  | 233 | 34945 | 35178 |

f

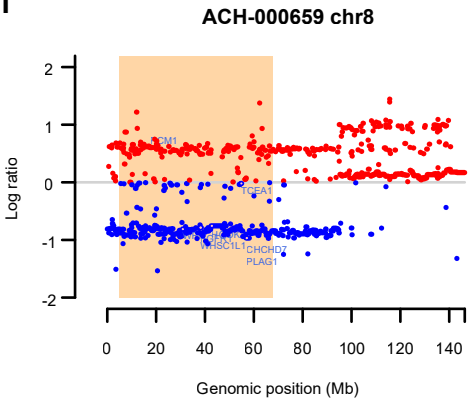

g

Cell Line: ACH-000659 | SCLC21H Lineage - CCLE: lung | PCAWG (matched): Lung-SCC
Data availability - SNP: True WES: False | Event Data Type & Overall Genome CNAs: SNP 681
Aneuploidy Score: 9 Ploidy: 1.93 | WGD Status: NA | Chromothripsis status: Positive
Chromosome: 8 Chromosome Size: 146,364,021 P Arm: 1 --- 43,838,887 | Q Arm: 46,838,887 --- 146,364,022
Number Of Segments: 590 Number Of Segments Filtered Out: 77 (13.05%)
Event Start Position: 31,254 | Event End Position: 146,298,155 (Event Length: 146,266,901) Number of Alterations: 450
New Log10LR: inf

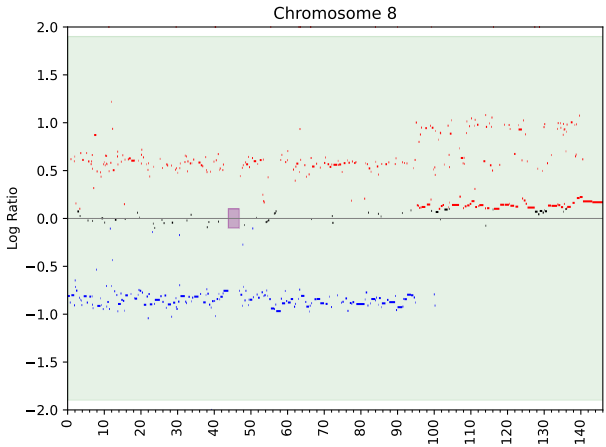

Extended Data Fig. 2

a

| DepMap ID | Cell Line Name | Cell type | Positive chromosome according to Stephens et al. (2011) | Positive chromosomes according to our method | Borderline chromosomes according to our method |
| --- | --- | --- | --- | --- | --- |
| ACH-000367 | NCIH226 | Squamous Cell Lung | 3 | 3 | 1 |
| ACH-000394 | NCIH2081 | Small Cell Lung | 3 | 1, 3 |  |
| ACH-000558 | A172 | Glioblastoma | 16 | 9, 16, 17 | 1, 5, 14, 19 |
| ACH-000580 | C32 | Melanoma | 11 | 11, 20 | 7 |
| ACH-000614 | RVH421 | Melanoma | X | 2 | 1 |
| ACH-000659 | SCLC21H | Small Cell Lung | 8 | 8 |  |
| ACH-000722 | SNUC1 | Colorectal | 15 | 15 |  |
| ACH-000790 | SHP-77 | Small Cell Lung | 4 | 4, 12, 16 | 1, 2, 7, 11 |
| ACH-000865 | KYSE450 | Esophageal | 4 | 4, 6 | 1, 5, 7, 8, 9, 11, 16, 22 |
| ACH-001208 | TK10 | Renal Cell | 5 | 5 | 17 |
| ACH-001274 | SW982 | Synovial Sarcoma | 1 |  |  |
| ACH-001307 | 8505C | Thyroid | 9 | 9 |  |
| ACH-002040 | HMVII | Melanoma | 3 | 3, 16 |  |
| ACH-002098 | CP67MEL | Melanoma | 1 | 1 |  |

b

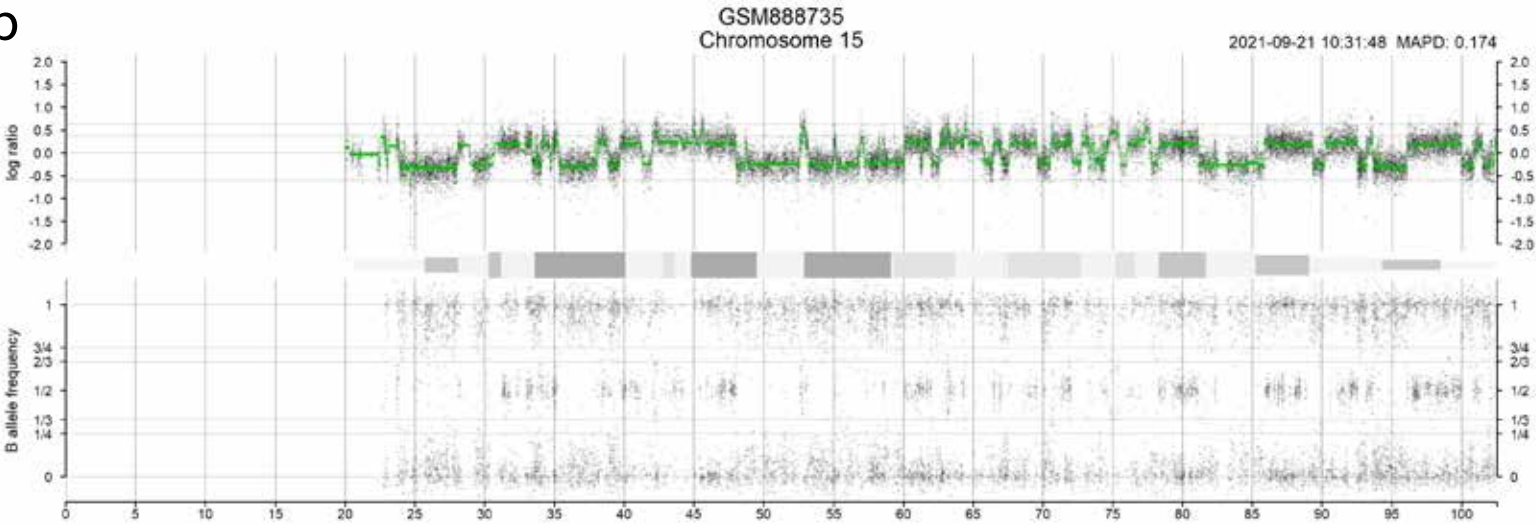

c

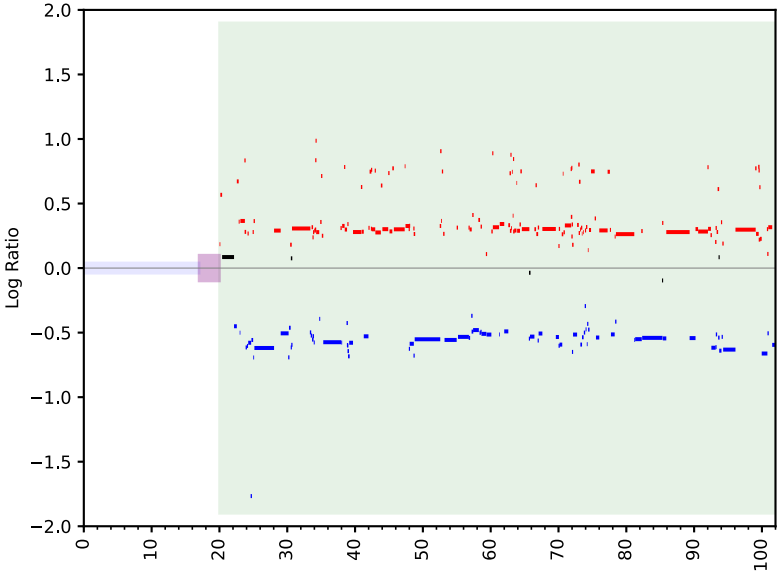

Extended Data Fig. 3

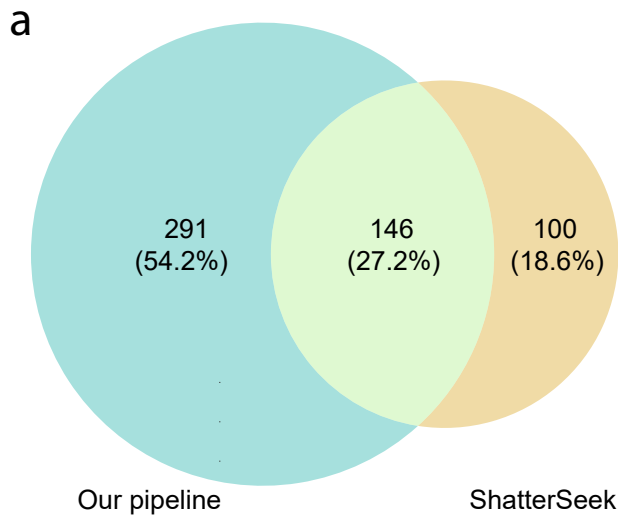

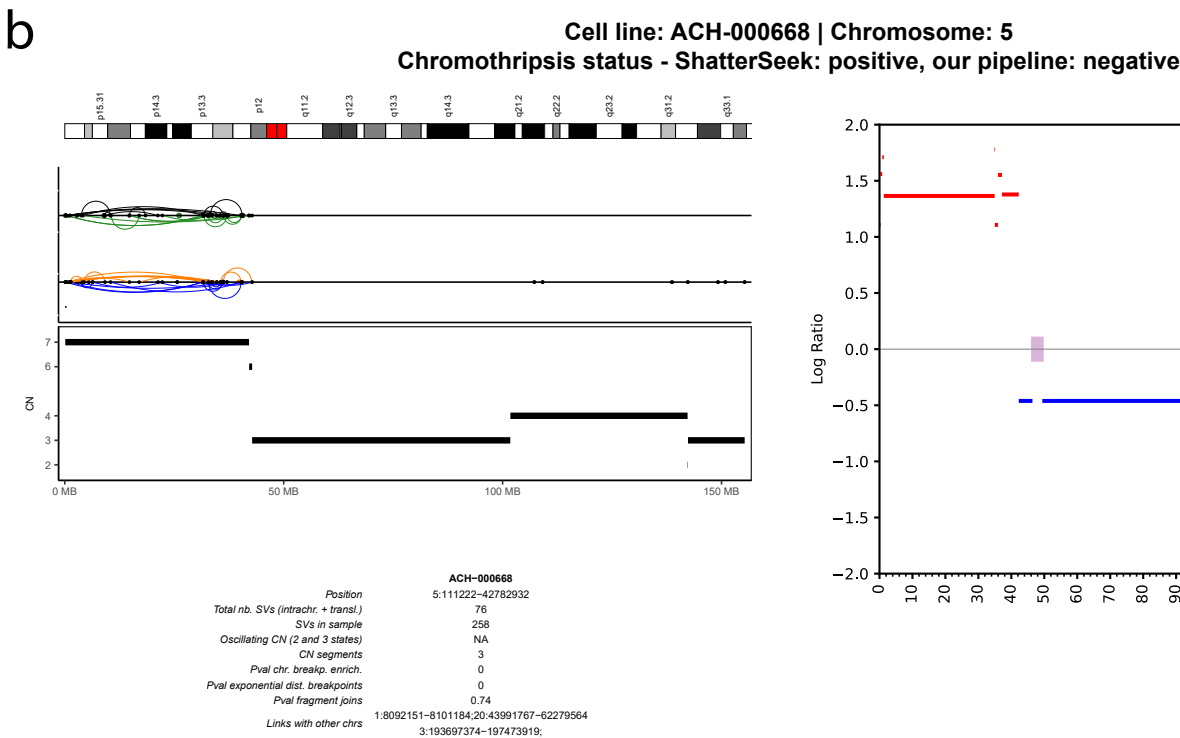

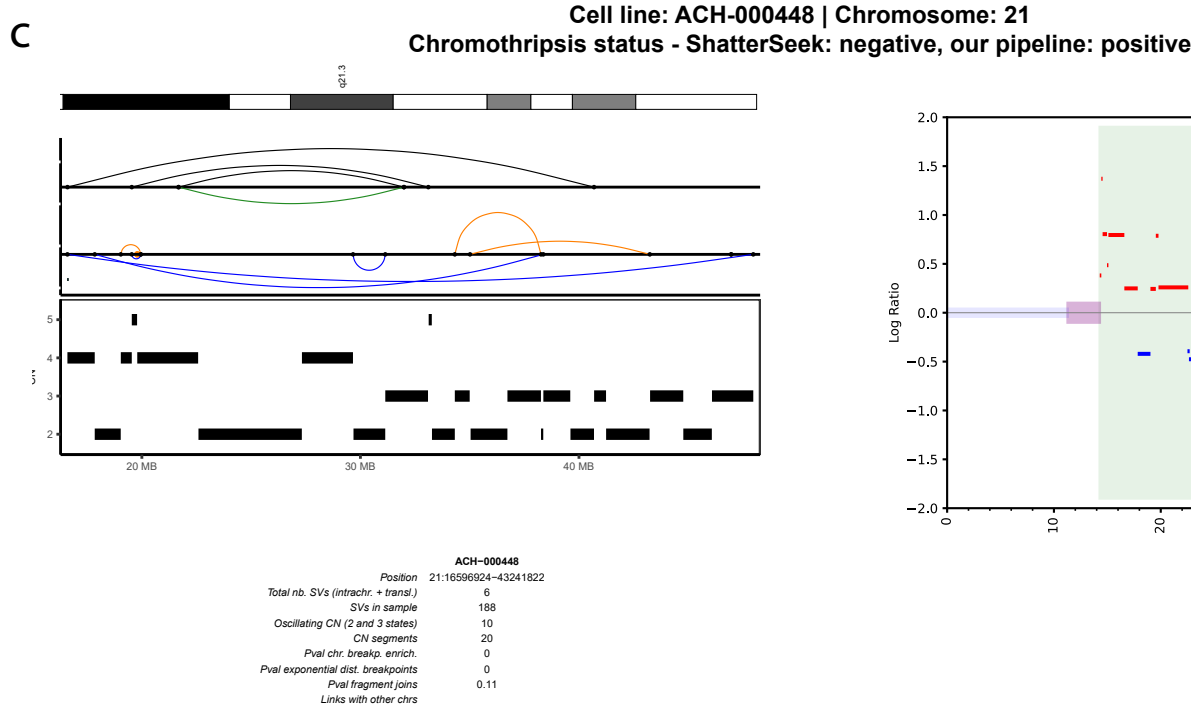

Extended Data Fig. 4

**a** Cell Line: ACH-000659 | SCLC21H CCLE Lineage: lung | Aneuploidy Score: 9, Ploidy: 1.93, WGD Status: nan
Positive Chromosomes: [8] Borderline Chromosomes: []

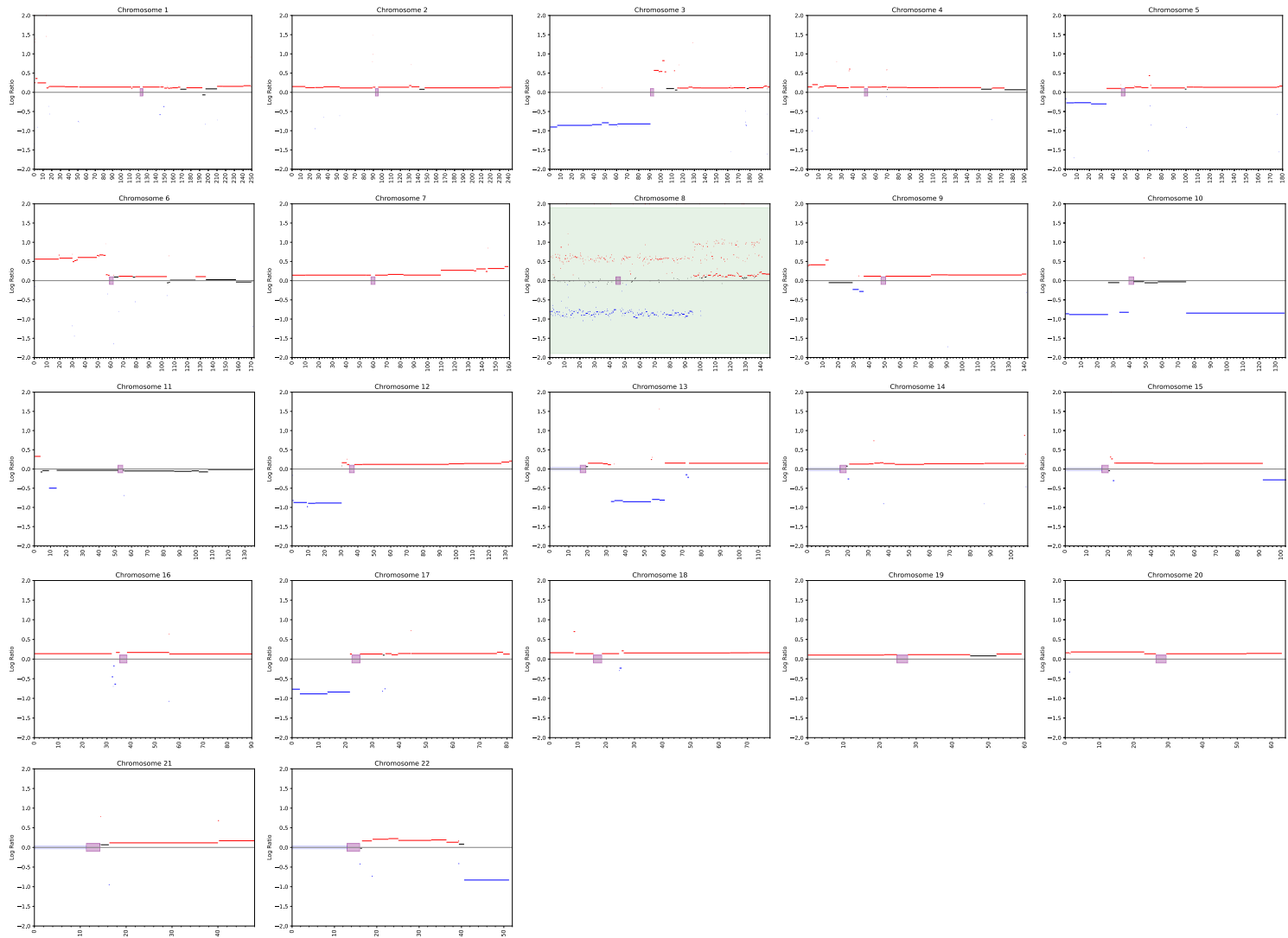

**b** Cell Line: ACH-000659 | SCLC21H Lineage: CCLE | PCAWG (matched): Lung-SCC
Data availability - SNP: True WES: False | Event Data Type & Overall Genome CNAs: SNP 681
Aneuploidy Score: 9 Ploidy: 1.93 | WGD Status: NA | Chromothripsis status: Positive
Chromosome: 8 Chromosome Size: 146,364,021 P Arm: 1 --- 43,838,887 | Q Arm: 46,838,887 --- 146,364,022
Number Of Segments: 590 Number Of Segments Filtered Out: 77 (13.05%)
Event Start Position: 31,254 | Event End Position: 146,298,155 (Event Length: 146,266,901) Number of Alterations: 450
New Log10LR: inf

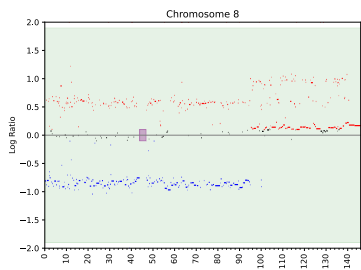

Is Whole Chromosome: True Is Whole P-Arm: True Is Whole Q-Arm: True Is Centromere: True
Distance Centromere: 0 Distance Telomere-P: 31,253 Distance Telomere-Q: 65,867
CN States: [-0.86, 0.12, 0.61, 0.86] | CN Dominant States: [-0.86, 0.12, 0.61]

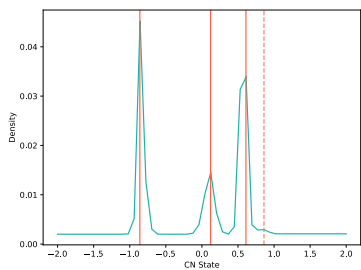

CTLPScanner Details: Length (Mb): 63.03 Start: 5,000,001 End: 68,025,520 Alterations: 262 Log10LR: inf

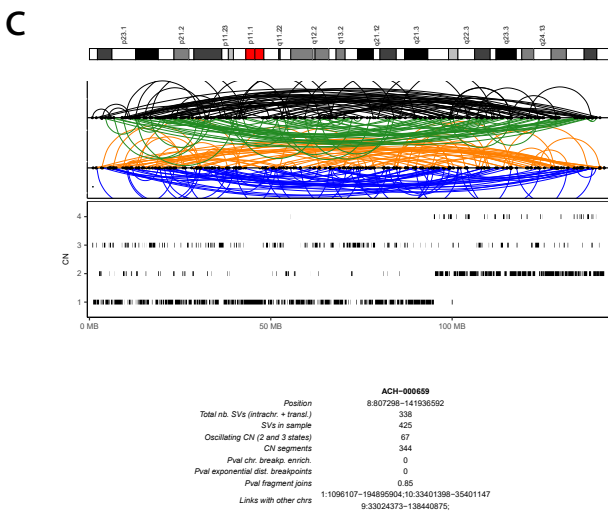

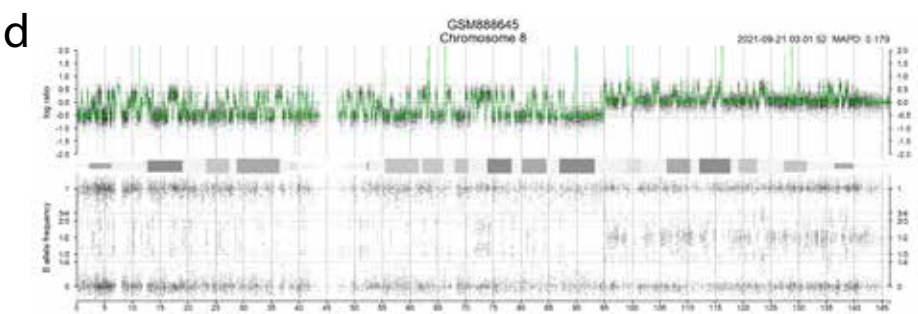

Extended Data Fig. 5

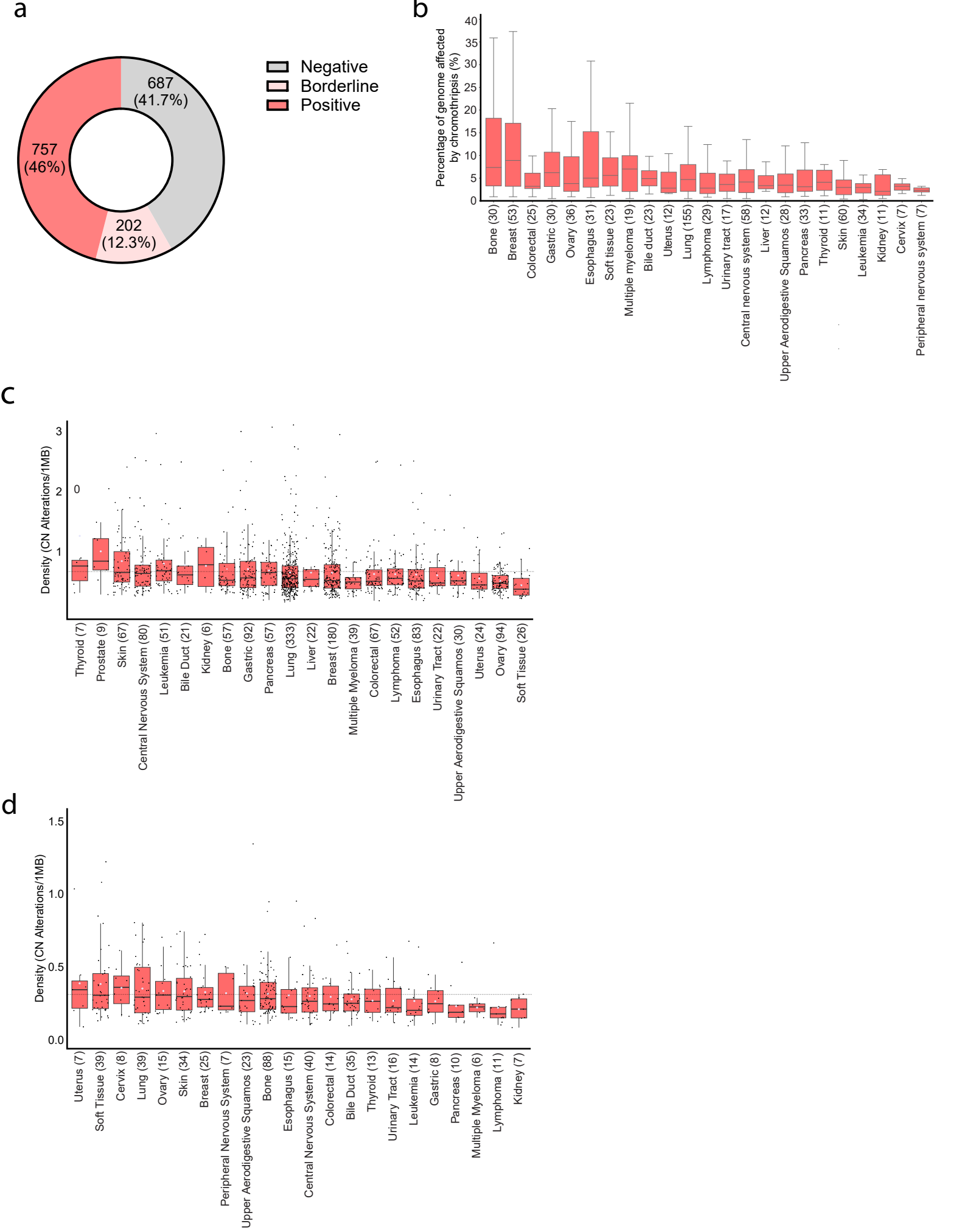

### Extended Data Fig. 6

**a**

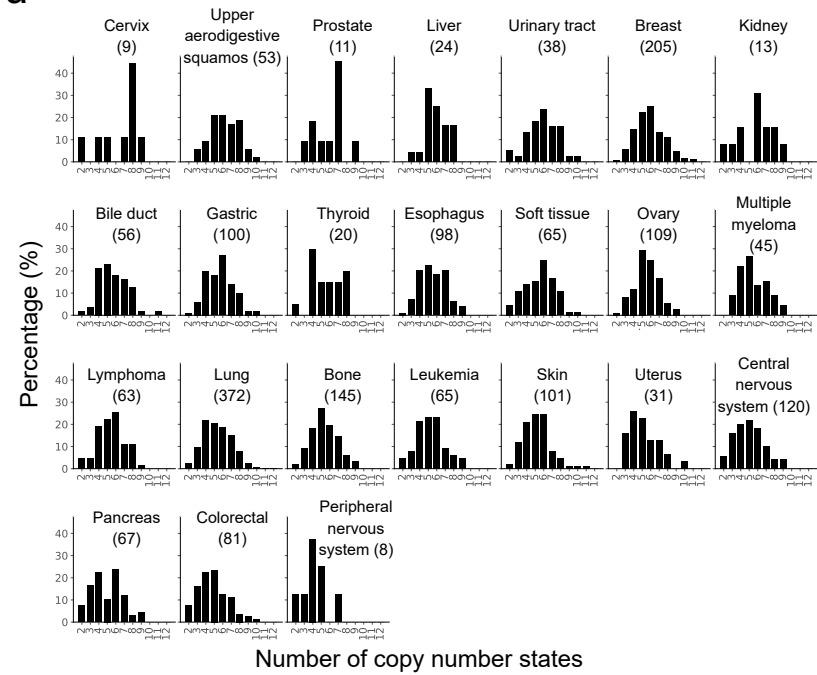

**b**

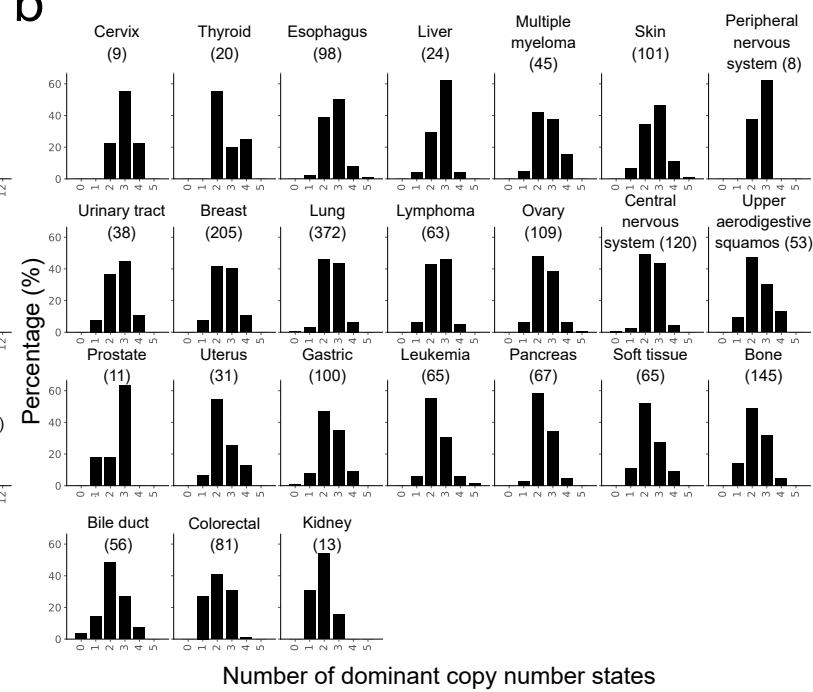

**c**

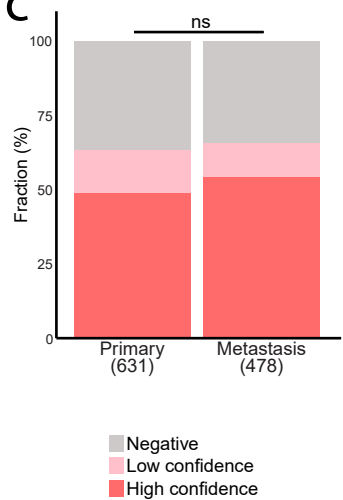

**d**

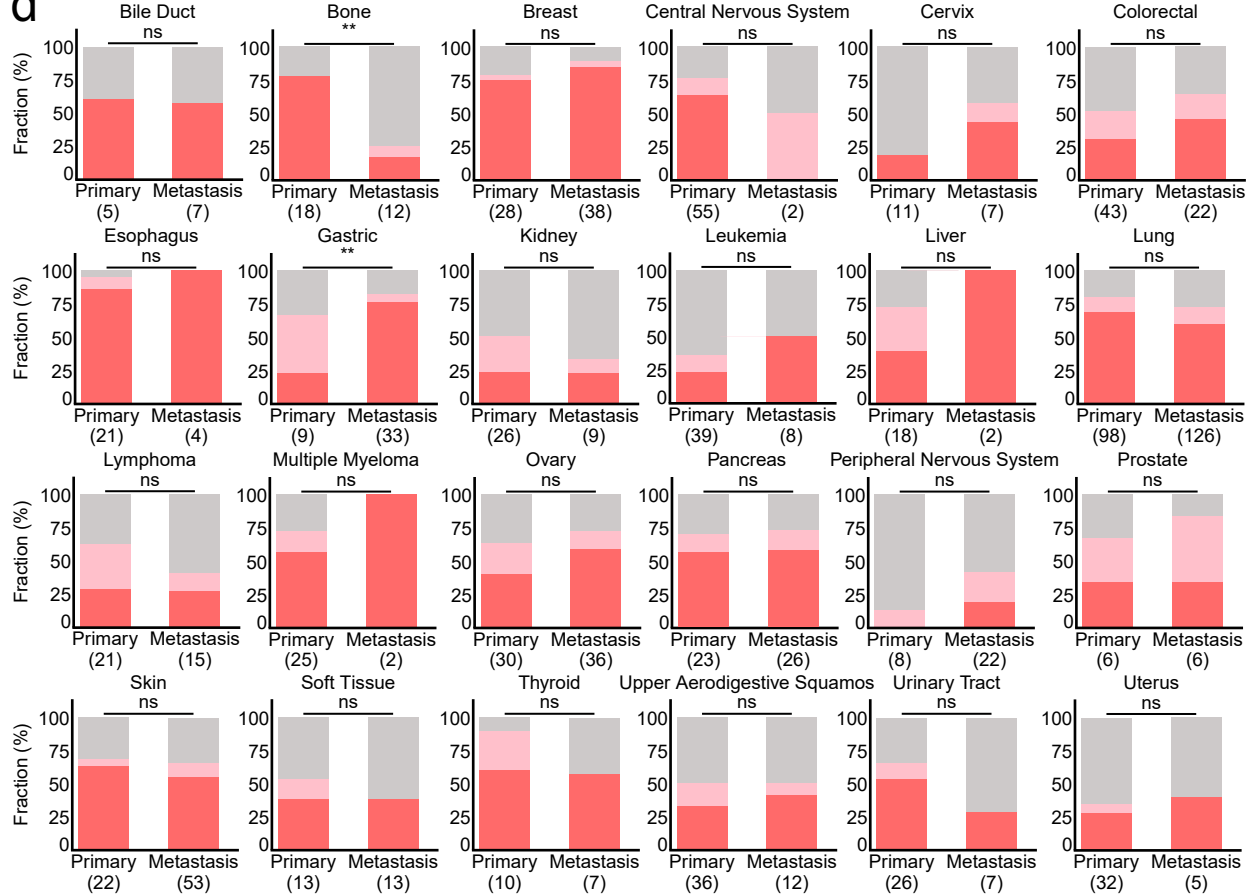

### Extended Data Fig. 7

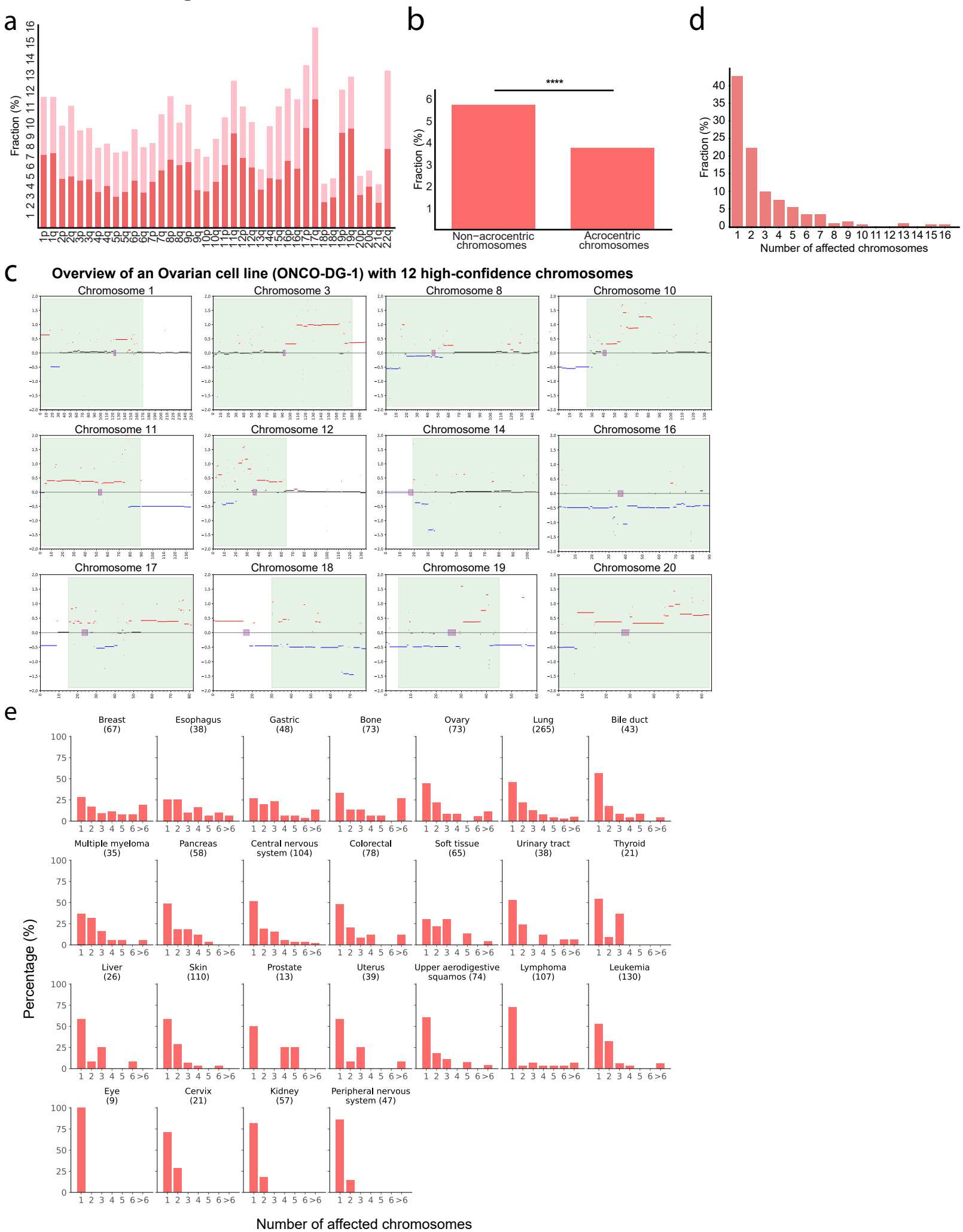

Extended Data Fig. 8

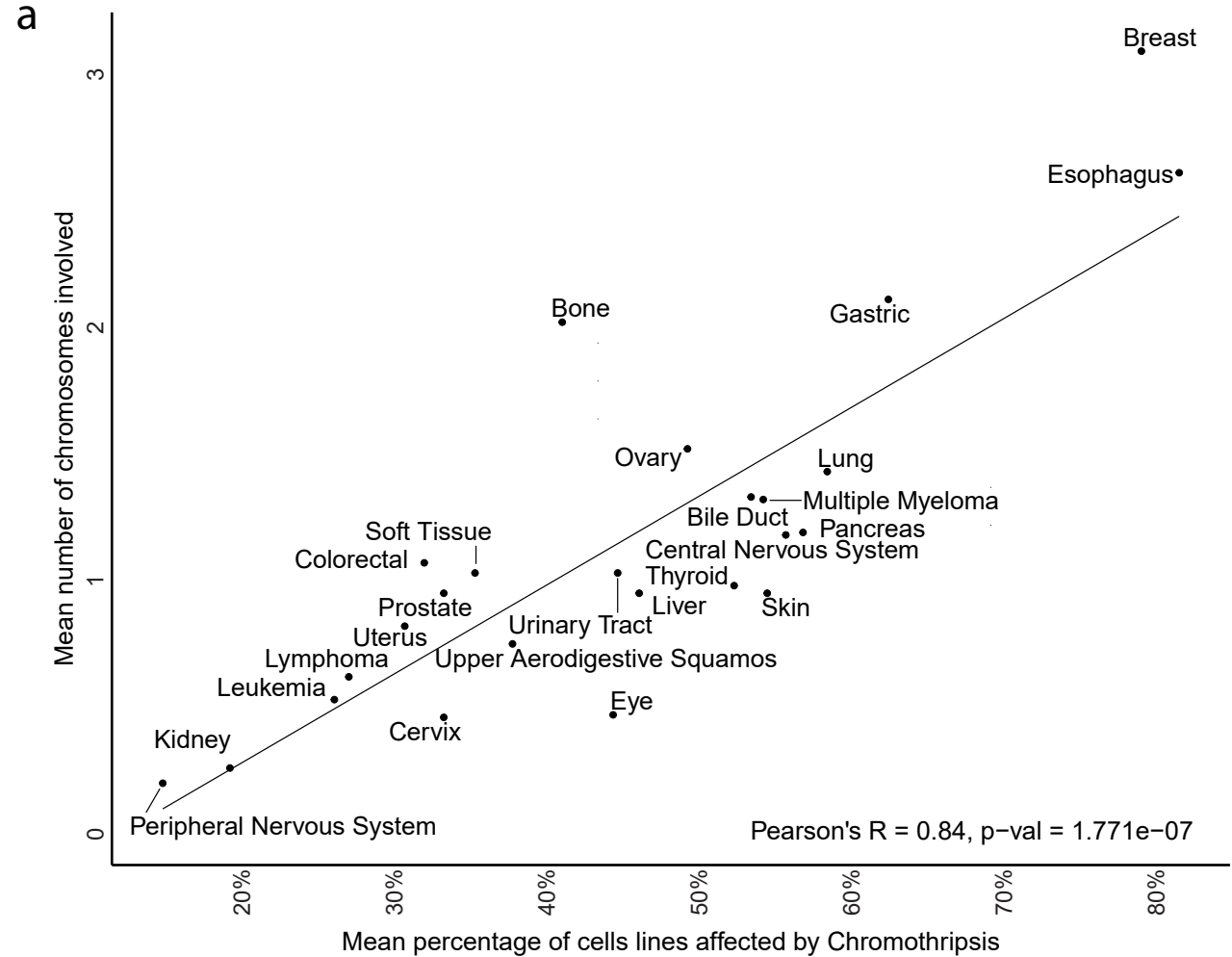

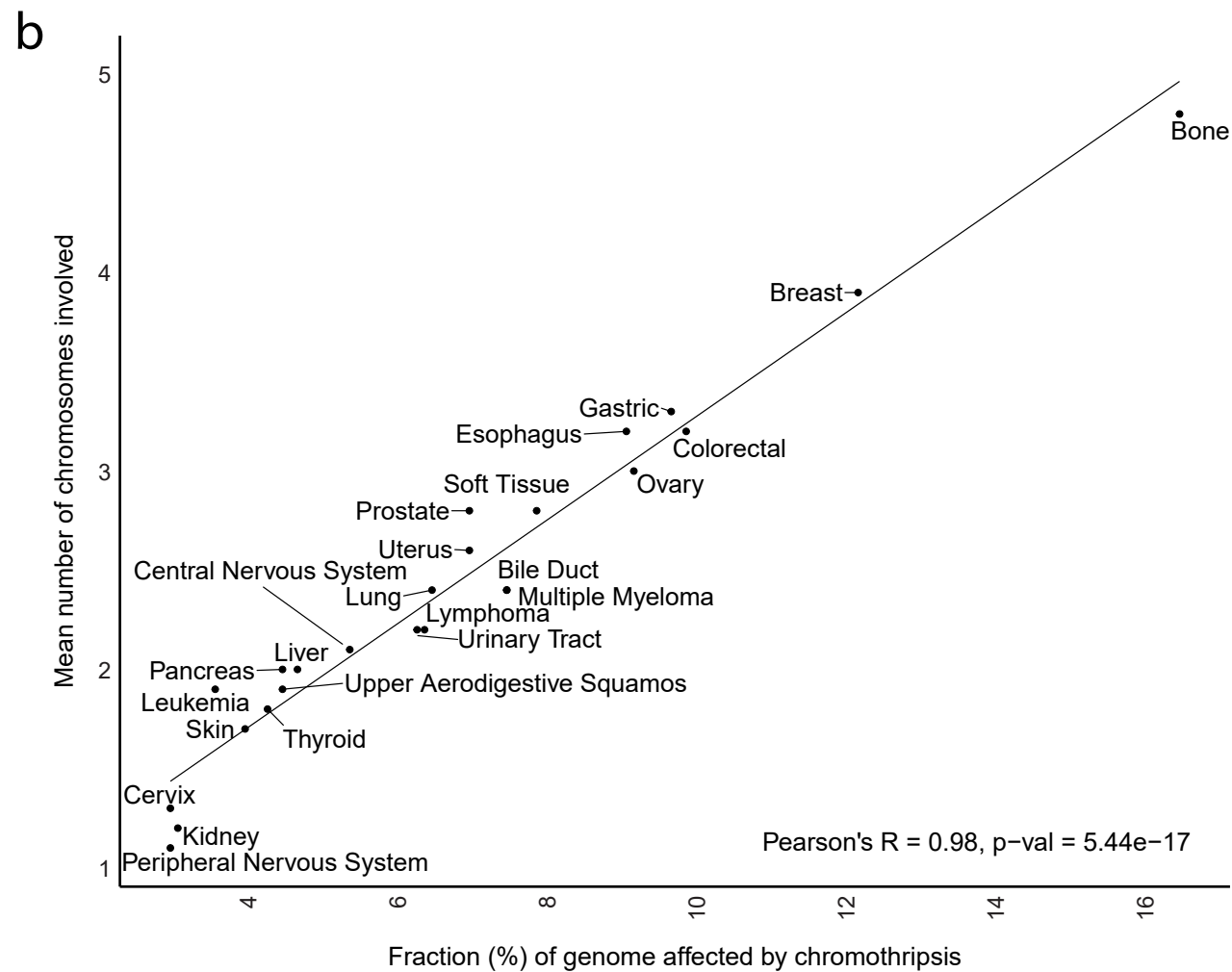

Extended Data Fig. 9

a

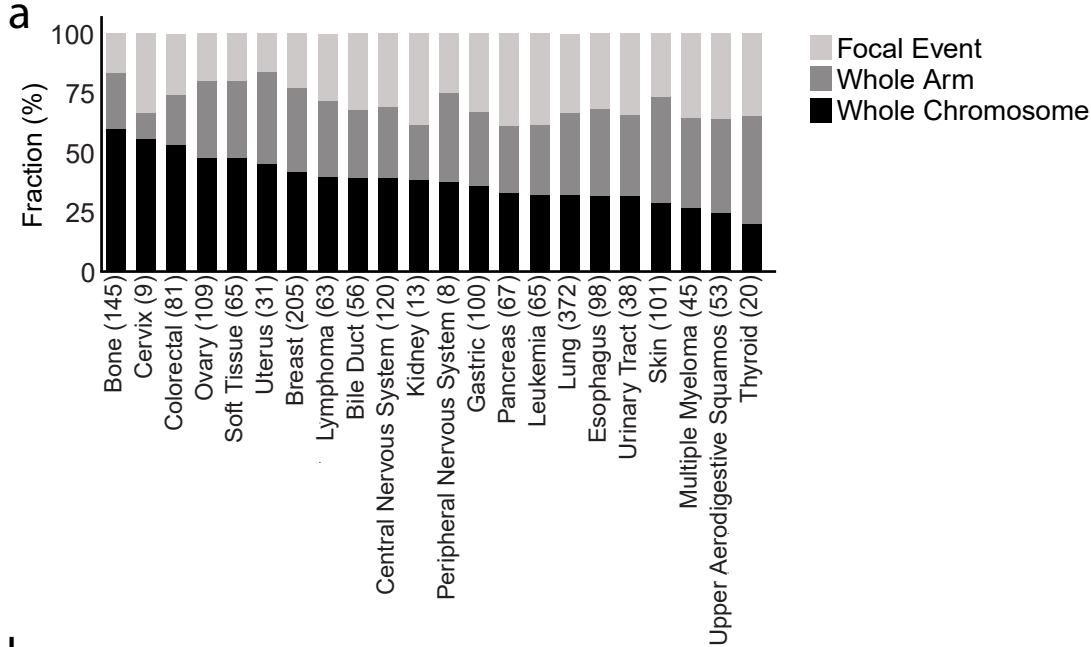

b

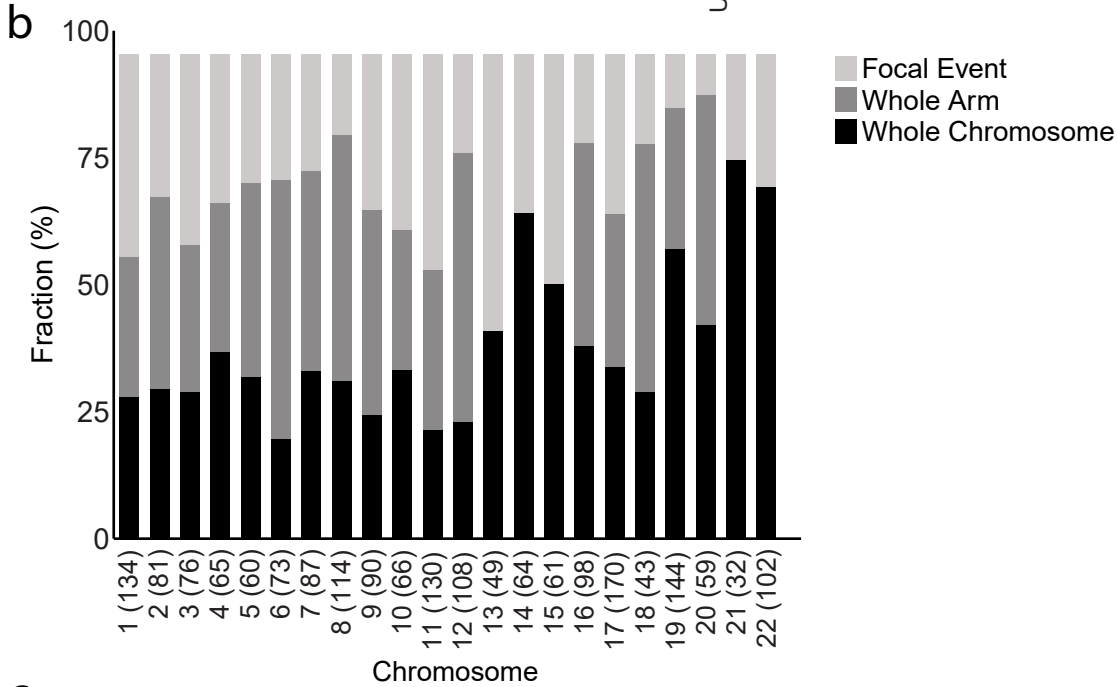

c

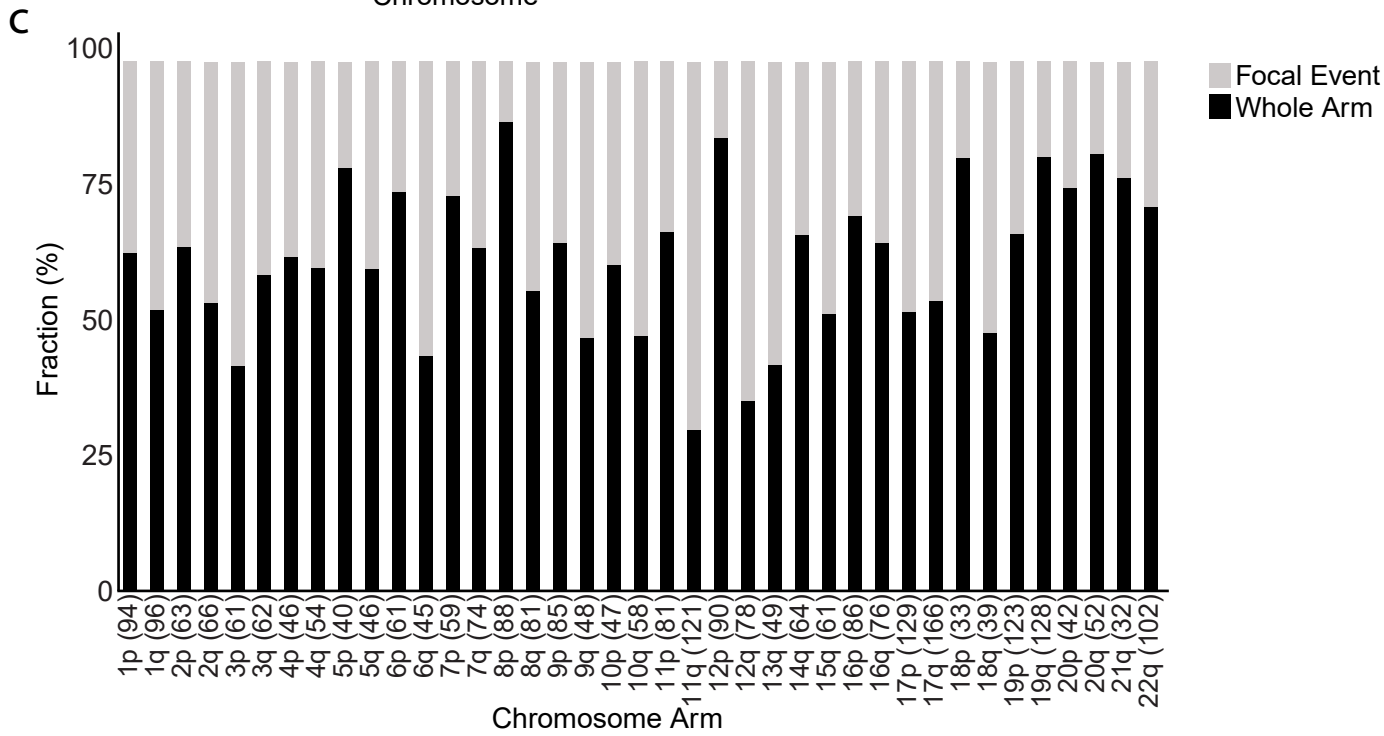

Extended Data Fig. 10

a

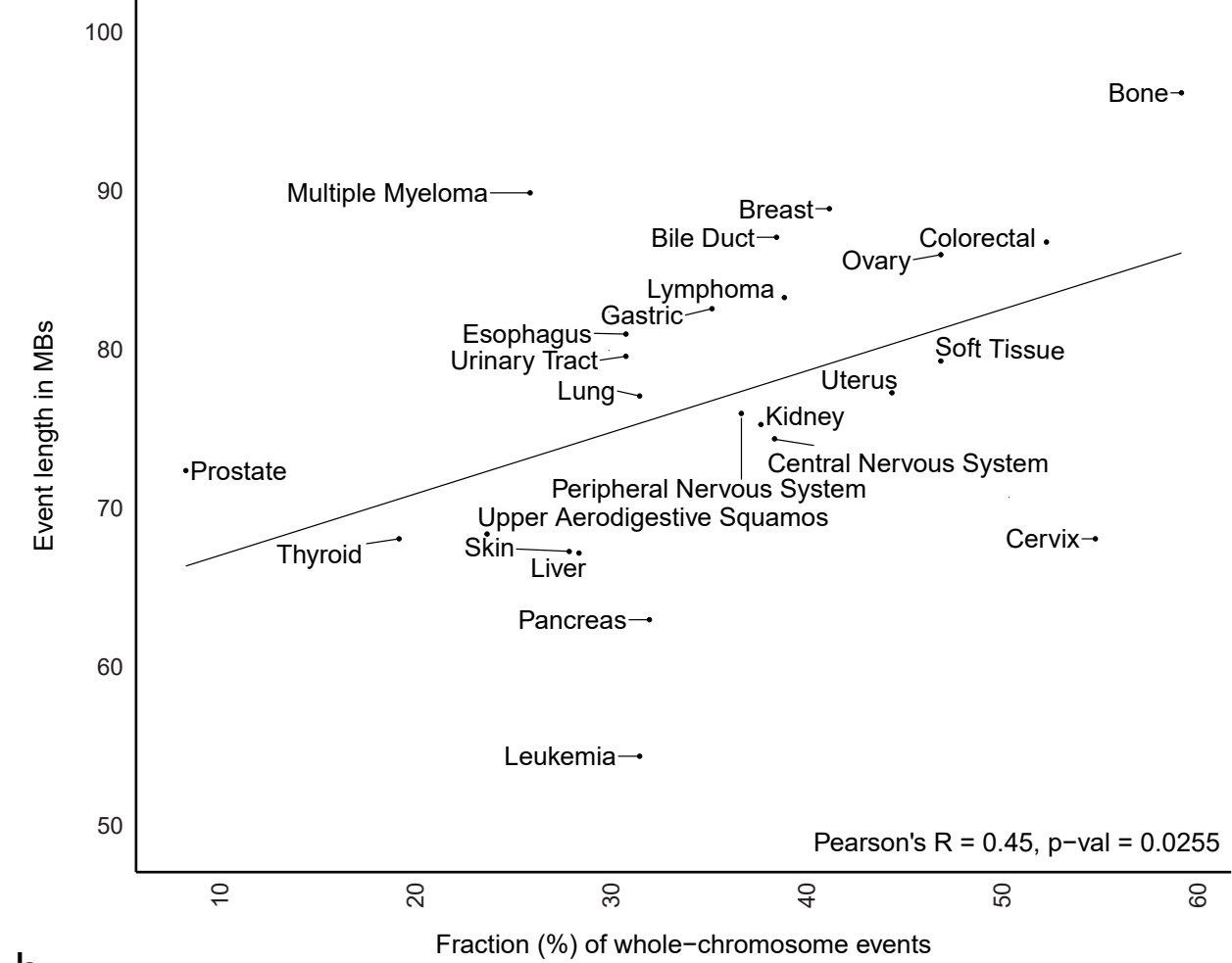

b

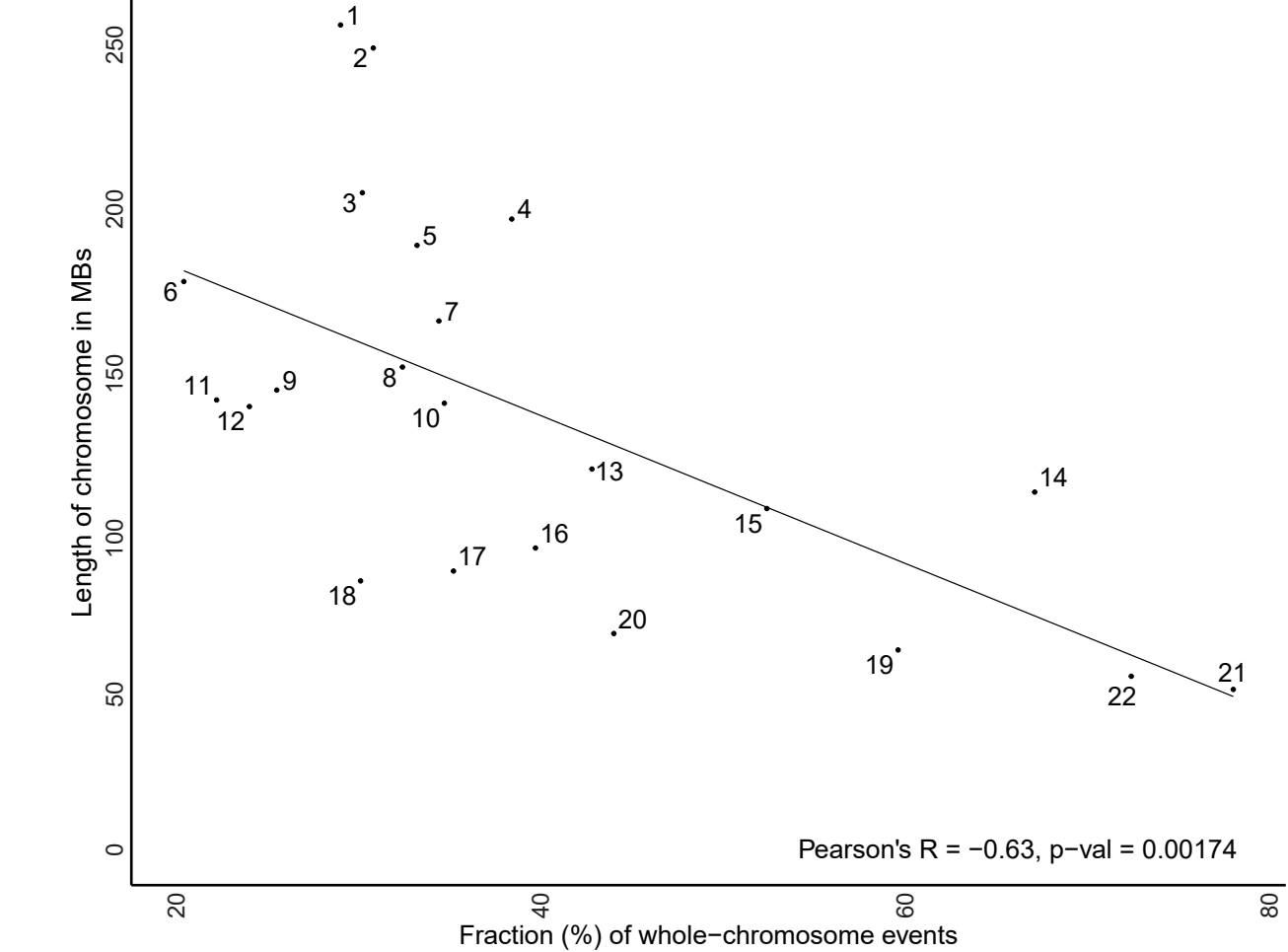

Extended Data Fig. 11

a

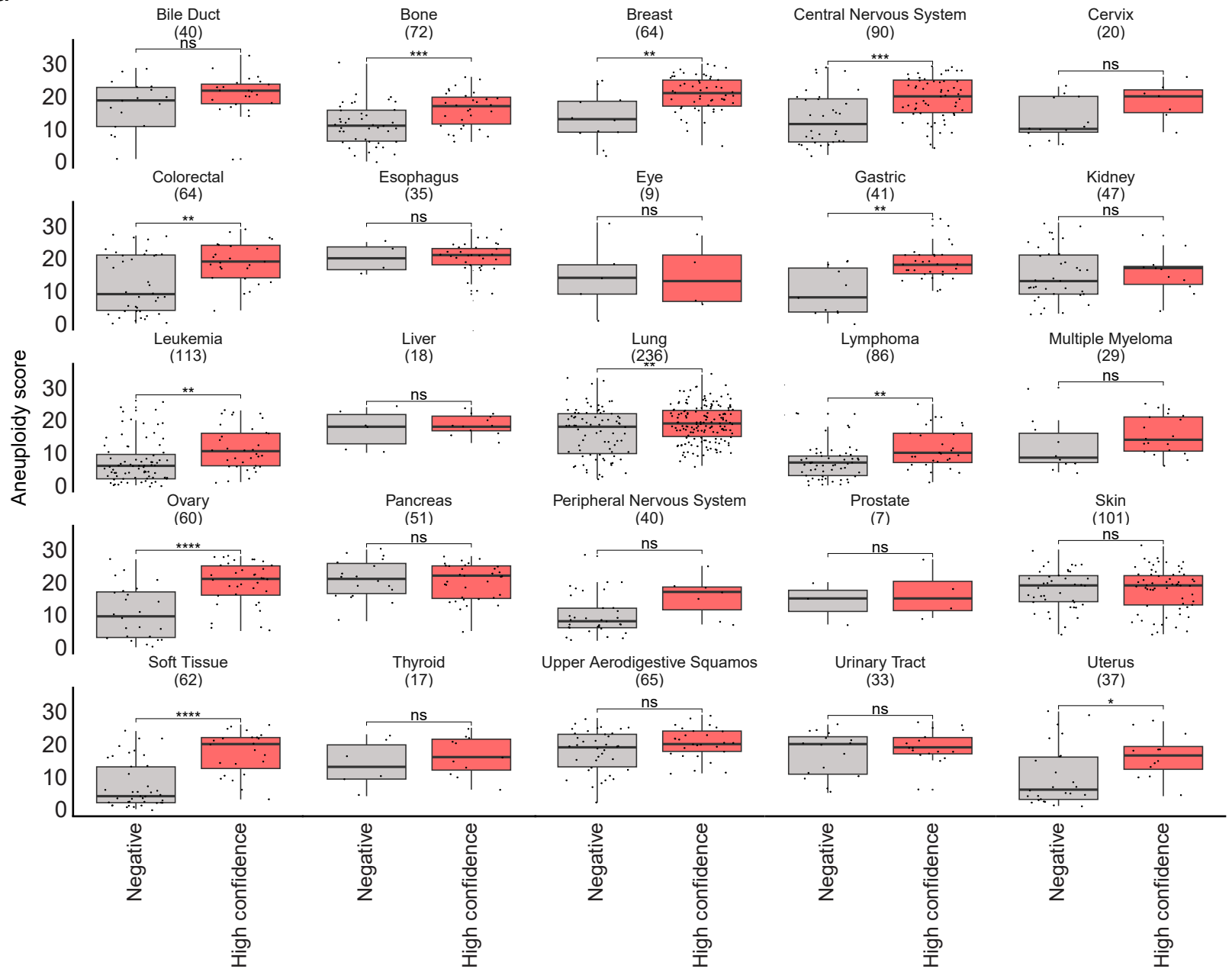

b

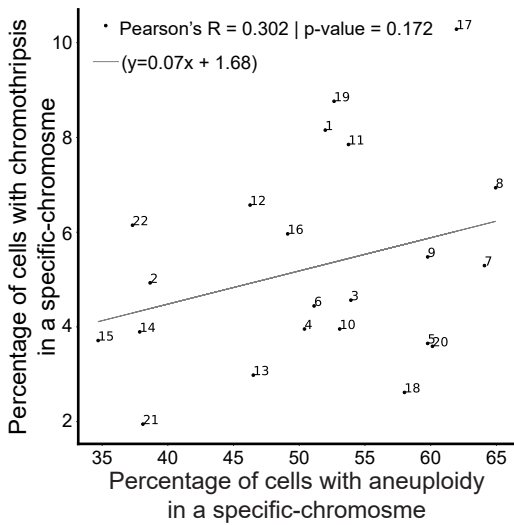

c

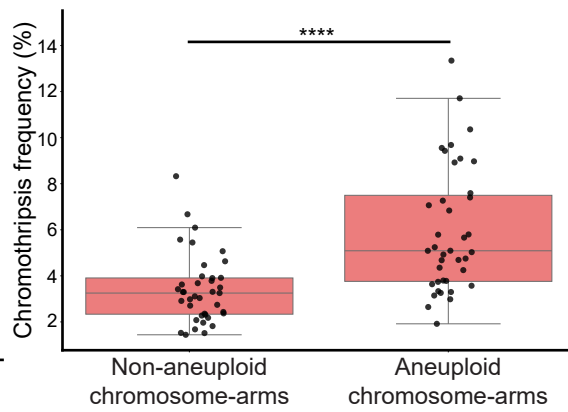

d

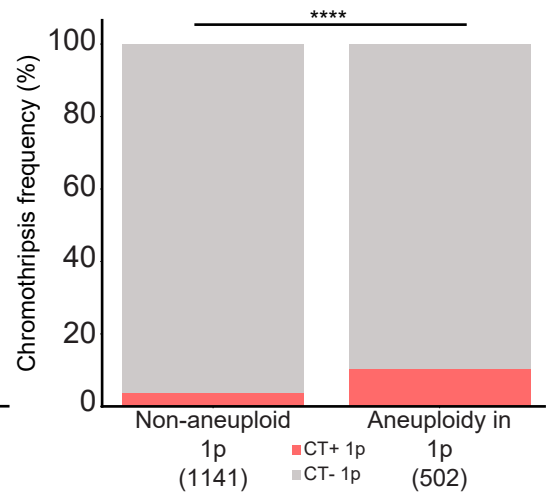

Extended Data Fig. 12

a

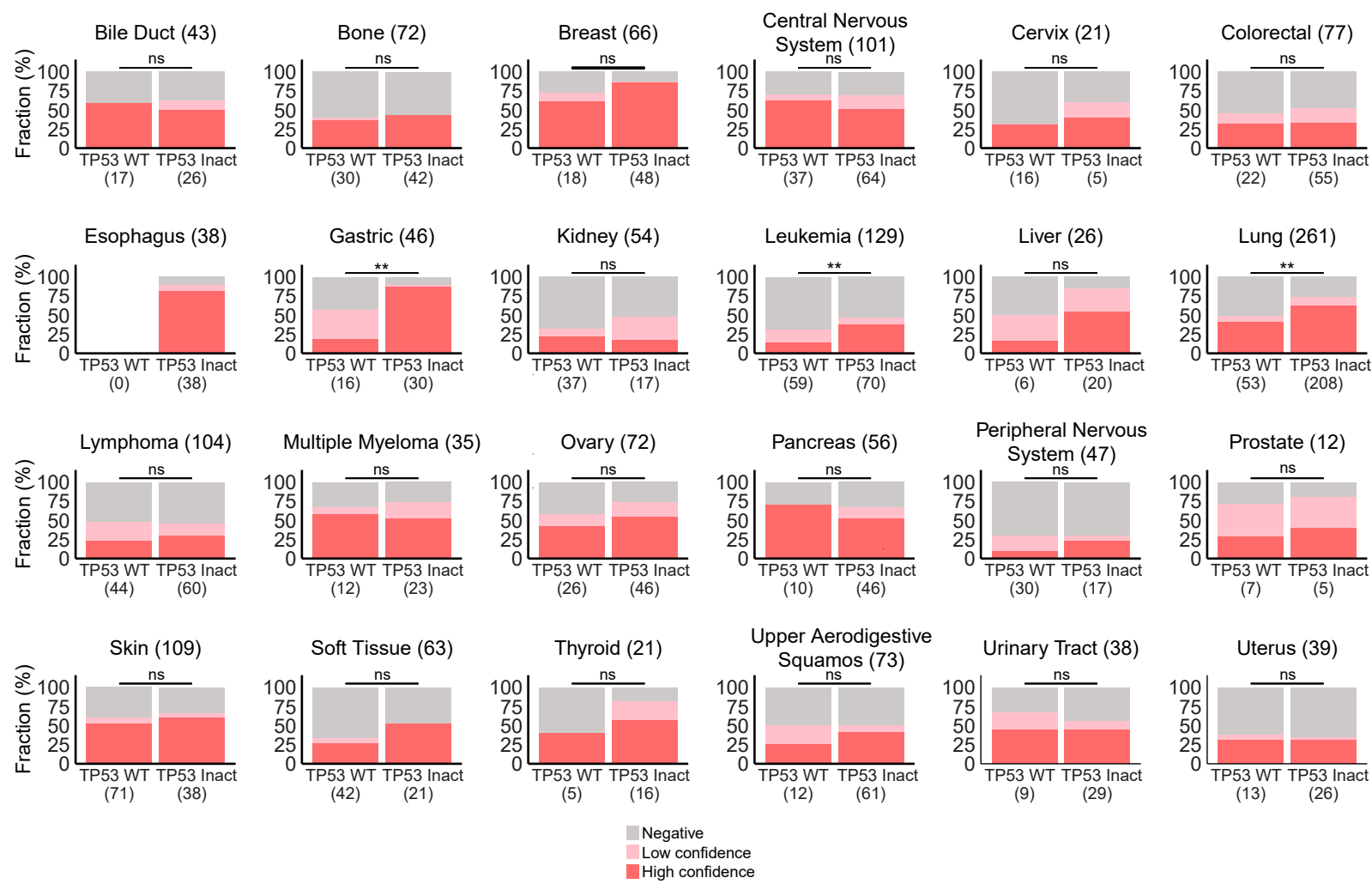

Extended Data Fig. 13

a

b

c

d

Extended Data Fig. 14

a

b

c

d

e

f

g

h

Extended Data Fig. 15

Extended Data Fig. 16

Extended Data Fig. 17

Extended Data Fig. 18

Extended Data Fig. 19

a

b

c

d

Extended Data Fig. 20
